# An autism spectrum disorder mutation in Topoisomerase 3β causes accumulation of covalent mRNA intermediates by disrupting metal binding within the zinc finger domain

**DOI:** 10.1101/2025.04.11.647616

**Authors:** Julia E. Warrick, Durga Attili, Trevor van Eeuwen, Benjamin Pastore, Sarah E. Hoffmann-Weitsman, Nicholas C. Forsyth, Wen Tang, Sami J. Barmada, Michael G. Kearse

## Abstract

The loss and mutation of Topoisomerase 3β (TOP3B), the only known eukaryotic topoisomerase with the ability to catalyze RNA strand passage reactions, is linked to schizophrenia, autism, and intellectual disability. Uniquely, TOP3B primarily localizes to the cytoplasm and has been shown to regulate translation and stability of a subset of mRNA transcripts. Three neurological disease-linked *de novo TOP3B* point mutations outside of the active site have been identified but their impact on TOP3B activity in cells remains poorly understood. Upon establishing a new Neuro2A cell-based TOP3B activity assay, we provide genetic and biochemical evidence that the autism-linked C666R mutation causes accumulation of unresolved TOP3B•mRNA covalent intermediates by directly disrupting metal coordination via an atypical D1C3-type metal binding motif within the zinc finger domain. Furthermore, we show that primary neurons are sensitive to TOP3B•mRNA covalent intermediates, including those formed by the C666R mutant TOP3B, and that such adducts are capable of causing ribosome collisions. Together, these data identify a previously underappreciated role of the zinc finger domain and how non-active site disease-linked mutations affect TOP3B activity and neuronal toxicity.

## INTRODUCTION

Topoisomerases are essential in all domains of life. Their canonical cellular role is to relieve topological strain that arises during DNA replication and transcription by performing strand passage reactions (1–4). Recent work has identified that metazoan Topoisomerase 3β (TOP3B), a type IA topoisomerase, is catalytically active on both DNA and RNA *in vitro* and in cells (5–8). This dual activity is physiologically important as multiple neurological disorders, including schizophrenia, autism, and intellectual disability, are linked to *TOP3B* deletion and mutation (6,9–12). Furthermore, multiple animal models present with measurable phenotypes often related to neurophysiology and cognition. For example, *TOP3B* null *Drosophila* demonstrate neuromuscular junction abnormalities (7,9,13), which is a known phenotype in *FmrI* null flies and in patients with Fragile X syndrome, a genetic disorder that also causes intellectual disability (14,15). *TOP3B* null mice (which mimics the most common deletion observed in the human population) display reduced fertility and life span, autoimmunity, impaired synapse formation, abnormal adult hippocampal neurogenesis, and behavioral abnormalities tied to cognitive impairment and psychiatric disorders (7,16–21).

Despite harboring a catalytic core that is conserved among type IA topoisomerases (22), multiple pieces of evidence support that mRNA is most likely the primary target of TOP3B. Unlike other mammalian topoisomerases, TOP3B is predominately localized to the cytoplasm, contains a *bona fide* RNA-binding RGG box domain, and co-sediments with the active translation machinery on polysomes (6,7,23). Deletion of the C-terminal RGG box domain severely decreases the ability of TOP3B to bind RNA in cells (9,24); the same mutation also robustly reduces recombinant TOP3B binding and catalytic function on RNA substrates *in vitro* (7,25). Mechanistic studies defining the biochemical requirements of TOP3B have reported that RNA is most likely the more favorable catalytic substrate due to the lower dependence on divalent ions (*i.e.*, Mg^2+^, Mn^2+^) and other active site residues to drive the reaction forward (25). However, increased R-loop formation, genome instability, and transcriptional changes have also been reported in cells and mice lacking TOP3B (20,26–29); therefore, it cannot be ignored that pathology linked to TOP3B could at least partially be influenced by its activity in the nucleus on chromatin.

In contrast to nearly all other RNA-binding proteins, TOP3B becomes covalently linked to its substrate during its natural catalytic cycle (2,30). Upon binding RNA substrates with aid from the C-terminal RGG box domain and proper positioning within the active site, the Y336 residue of TOP3B (human isoform 1 numbering) performs a nucleophilic attack on the phosphodiester backbone, forming a covalently linked TOP3B•RNA intermediate. After protein conformational changes allow strand passage, the substrate is subsequently rejoined and TOP3B presumably dissociates before starting another cycle or repeats the cycle multiple times during one binding event to fully resolve a structure (2,31–33). Although a consensus binding sequence or structure has not been reported for TOP3B, HITS-CLIP and eCLIP studies have identified that mammalian TOP3B primarily binds to the open reading frame (ORF) of mRNAs (7,34). Such enriched binding to ORFs is indicative of a role in regulating mRNA translation and/or metabolism. Indeed, a recent study of cells harboring wildtype or catalytically-inactive (Y336F) alleles concluded that TOP3B regulates mRNA translation and stability by two distinct modes: i) by acting as more of a traditional RNA-binding protein where catalytic activity is not required, or ii) by acting as an RNA topoisomerase where catalytic activity is necessary (34).

In addition to the more common pathogenic deletion of the *TOP3B* locus, three *de novo* point mutations have been reported in individuals presenting with neurological disorders (35–37). In this report, we evaluated the impact of all three non-active site point mutations on the ability of TOP3B to function on mRNA in Neuro2A cells. Our data show that the autism-linked C666R mutation results in accumulation of unresolved TOP3B•mRNA covalent intermediates. Further genetic and biochemical analyses support that the C666 residue is directly involved in metal coordination within the zinc finger domain. Using primary rat cortical neurons as a model, we further define how various levels of TOP3B activity and the ability to cause unresolved TOP3B•mRNA covalent intermediates affect neuron survival. Lastly, we show that robust unresolved TOP3B•mRNA covalent intermediates interfere with mRNA translation by causing ribosome collisions. Together, these data provide critical new insights into how neurological disease mutations affect TOP3B and how unresolved TOP3B•mRNA covalent intermediates impact neuronal toxicity and translational control.

## MATERIALS AND METHODS

### Plasmids

pCMV6/TOP3B(WT)-FLAG plasmid that encodes isoform 1 of human TOP3B (RefSeq: NM_003935) was obtained from Origene Technologies (catalog # RC223204). All mutations, including replacing the FLAG tag with a 3xFLAG tag, were made using the Q5 Site Directed Mutagenesis Kit (NEB # E0554S). pcDNA3.1(+)/mEGFP that was used as a transfection control was a kind gift from Jeremy Wilusz (Baylor College of Medicine). pcDNA3.1(+)/3xV5-Ubiquitin was generated by cloning the synthesized 3xV5-Ubiquitin (WT and the indicated mutants) open reading frame (Integrated DNA Technologies) into pcDNA3.1(+) (Thermo Fisher # V79020).

The 3xFLAG-mEGFP-NES-TOP3B(WT) open reading frame was synthesized by Genscript and subcloned into pcDNA5/FRT/TO (for expression in Flp-In T-Rex HEK293 cells) or pGW1 (for expression in primary neurons). pcDNA5/FRT/TO vector and pOG44 plasmid were obtained as part of the Flp-In T-REx Core Kit (Thermo Fisher # K650001). Point mutations were introduced by Q5 Site Directed Mutagenesis.

Recombinant protein expression of MBP-TOP3B ZnF fusion proteins in ExpiSf9 insect cells used the pFastBac Dual vector (Thermo Fisher # 10712024). mEGFP was PCR amplified and cloned downstream of the p10 promoter, which allowed for the easy identification of transfected and transduced cells. FLAG-MBP-TOP3B ZnF (G618-S818)-twinSTII was PCR amplified and cloned downstream of the polyhedrin promoter. The twin StrepII (twinSTII) tag with flexible linkers (GSG<u>WSHPQFEK</u>GS<u>WSHPQFEK</u>GSS; based from Shah *et al*. (38)) was inserted directly upstream of the stop codon of TOP3B. Recombinant expression of full-length TOP3B in ExpiSf9 insect cells used the pFastBac1 vector (Thermo Fisher # 10360014); His6-TOP3B-FLAG was PCR amplified and cloned downstream of the polyhedrin promoter. Point mutations were introduced by Q5 Site Directed Mutagenesis.

All plasmids were propagated in TOP10 *E. coli* (ThermoFisher # C404006) and purified using the PureYield Miniprep (Promega # A1222) or the ZymoPURE II Plasmid Midiprep Kit (Zymo Research # D4201). All plasmids were validated by Sanger Sequencing at The Ohio State University Comprehensive Cancer Center Genomics Shared Resource or by whole plasmid sequencing through Plasmidsaurus. Nucleotide sequences of all proteins used in this study are provided in **Supplementary Data**. All plasmids are available upon request.

### Cell culture, transfection, and drug treatments

Neuro2A cells were obtained from ATCC (catalog # CCL-131) and maintained in high glucose DMEM (Thermo Fisher # 11995065) supplemented with 10% heat-inactivated Fetal Bovine Serum (Cytiva # SH30396.03HI) and 1% Penicillin-Streptomycin (Thermo Fisher # 15140122) at 37°C and 5% CO_2_ in standard tissue culture treated plates. Neuro2A cells were seeded in 12-well and 6-well plates for Western blot and slot blot experiments, respectively. 24 hrs later, at ∼50% confluency, cells were transfected using Fugene 4K (Promega # E5911) using a 4:1 ratio. Briefly, 4 µL of Fugene 4K was mixed with 1 µg total plasmid (500 ng TOP3B + 500 ng GFP or 400 ng TOP3B + 400 ng 3xV5-Ubiquitin + 200 ng GFP) in 100 µL Opti-MEM (ThermoFisher # 31985070). After 10 min at room temperature, the transfection mix was diluted 1:2 with Opti-MEM and added dropwise to cells (45 µL and 95 µL were added to each well of 12-well and 6-well plates, respectively). 24 hours after transfection, media was aspirated, cells were gently washed with 1 mL ice-cold PBS, and then lysed in well with either 600 µL RIPA buffer + protease inhibitors (for 12-well plates and Western blots) or 500 µL Oligo dT Lysis Buffer (100 mM Tris-HCl, 500 mM LiCl, 0.5% LDS, 1 mM EDTA, 5 mM DTT; pH 7.5) (for 6-well plates and slot blot experiments). Plates were incubated with rocking for 10 min at 4°C (12-well plates) or room temperature (6-well plates), and then lysates were transferred to RNase- and protease-free microcentrifuge tubes and stored at −80°C or immediately used.

When using TAK243 and MG132 to inhibit the ubiquitin-proteasome system, fresh media containing 10 µM TAK243 (Selleckchem # S8341; 10 mM stock in DMSO) or 10 µM MG132 (Sigma # M7449; 10 mM stock in DMSO) was exchanged 21 hrs after transfection and placed back at 37°C, 5% CO_2_. 3 hrs later, lysates were harvested as described above.

### TOP3B•mRNA activity assay and slot blotting

mRNA isolation for the TOP3B•mRNA activity assay was performed using the Magnetic mRNA Isolation Kit (NEB # S1550S) with minor changes of the manufacturer’s protocol. We found it was more cost effective to purchase the Oligo dT(25) magnetic beads (NEB # S1419S) alone and make buffers in house (which were made with RNase-free milliQ water, filtered through a 0.2 µm cellulose nitrate membrane (Corning # 430758), and stored at room temperature). We routinely observed that the buffers detailed below performed better when made within 3 months of use. DTT in the below buffers was added fresh to working solutions on the day of use.

Cell lysates were syringed 1X with a 28G needle to reduce sample viscosity. For each sample, 100 µL of bead slurry was equilibrated with Oligo dT Lysis Buffer (100 mM Tris-HCl, 500 mM LiCl, 0.5% (w/v) LDS, 1 mM EDTA, 5 mM DTT; pH 7.4) in an RNase-free microcentrifuge tube. Syringed cell lysates were incubated with equilibrated beads for 30 minutes at room temperature with gentle end-over-end rotation (15 rpm). Beads were then separated using a magnetic rack for 2 min and the supernatant was carefully removed to avoid disturbing the beads. Samples were washed twice with 500 µL of Oligo dT Wash I Buffer (20 mM Tris-HCl, 500 mM LiCl, 0.1% (w/v) LDS, 1 mM EDTA, 5 mM DTT, pH 7.4), twice with 500 µL of Oligo dT Wash II Buffer (20 mM Tris-HCl, 500 mM LiCl, 1 mM EDTA; pH 7.4), and once with 500 µL of Oligo dT Low Salt Buffer (20 mM Tris-HCl, 200 mM LiCl, 1 mM EDTA; pH 7.4). For each wash, beads were gently resuspended by inversion and then separated using a magnetic rack for 2 min. mRNA was eluted by adding 200 µL TE, briefly vortexing to resuspend the beads, and heating at 50°C for 2 min. Beads were then separated using a magnetic rack for 2 min and the supernatant was transferred to an RNase-free centrifuge tube. Samples were then centrifuged at 18,000 rcf for 5 min at room temperature to remove the small amount of residual beads that were not removed by the magnetic rack. 185 µL of the supernatant was transferred to an RNase-free microcentrifuge tube as the final eluate, quantified using a NanoDrop One Spectrophotometer, and used immediately or stored at −80°C. When confirming the dependence on RNA capture (vs. DNA contamination) by slot blotting (**Supplementary Figure S1A**), following the Oligo dT Low Salt Buffer wash, beads were incubated with 500 µL of Oligo dT Low Salt RNase Buffer (20 mM Tris-HCl, 200 mM LiCl, 10 mM MgCl_2_; pH 7.4) with or without 10 µL RNase I_f_ (500 units total; a recombinant protein fusion of *E. coli* RNase I; NEB # M0243S) for 30 minutes at 25°C at 500 rpm. Beads were then washed once more with Oligo dT Low Salt Buffer and continued with elution as described above.

Slot blot samples were prepared by diluting 2 µg mRNA with sterile PBS in a total volume of 200 µL and heating at 70°C for 15 min. We empirically found that heat denaturing alone was far superior to SDS and urea-based treatments for the subsequent immunodetection. A Hoefer PR 648 Slot Blot Blotting Manifold or Bio-Rad Bio-Dot Slot Format (SF) Microfiltration Apparatus was used for all slot blots with standard building vacuum power (19-25 inHg). 0.2 µm nitrocellulose membrane (BioRad # 1620112) and 0.2 µm Brightstar-Plus nylon (Thermo Fisher # AM10100) membrane were cut to size and first equilibrated in 20 mM Tris-HCl, pH 7.4 for 1 hour at room temperature prior to slot blotting. The Hoefer PR 648 Slot Blot Blotting Manifold apparatus with a 4 mm bore Nalgene three-way stopcock (Thermo Fisher # 64700004) was assembled with two nylon membranes underneath one nitrocellulose membrane, and the Bio-Rad Bio-Dot Slot Format (SF) Microfiltration Apparatus was assembled with two pieces of filter paper underneath both membranes. For all subsequent steps, washes and samples were added without vacuum; only once all washes and samples were placed in each slot was vacuum applied. After assembly, membranes were washed twice with 195 µL of 20 mM Tris-HCl, pH 7.4 in each slot. 195 µL of prepared sample was added to each slot and then washed once again with 195 µL of 20 mM Tris-HCl, pH 7.4. The slot blot apparatus was then disassembled and the top nylon membrane was UV crosslinked using the Stratagene Stratalinker UV 1800 crosslinker with 120,000 µJ/cm^2^ (Auto Crosslink function). RNA was subsequently detected by staining with 0.2% (w/v) methylene blue (in 0.4 M sodium acetate and 0.4 M acetic acid) for 20 min at room temperature with orbital shaking, washed with distilled water, and then imaged on the BioRad GelDoc Go Imaging System using the Coomassie Blue setting. In parallel, the nitrocellulose membrane was incubated in 5% (w/v) non-fat dry milk (NFDM) in TBST (1X Tris-buffered saline with 0.1% (v/v) Tween 20) for 30 min at room temperature with orbital shaking, washed 3X in TBST, and incubated in primary antibody overnight at 4°C with rocking. Mouse anti-FLAG M2 (Sigma # F3165-1mg) and rabbit anti-V5 (Cell Signaling # 13202; clone D3H8Q) were used at 1:1,000 dilution in TBST with 0.02% (w/v) sodium azide. Subsequent steps for immunodetection were performed as described below for Western blotting.

### Western blotting

Samples were prepared by adding 4X reducing Laemmli Sample Buffer (Bio-Rad # 1610747) with βME as a reducing agent, heating at 70°C for 15 min, and syringing samples with 28G needle. 15 µL sample was then separated by 4-20% Tris-glycine SDS-PAGE (Invitrogen # XP04200BOX), transferred onto 0.2 µm PVDF membrane (Thermo Fisher # 88520), and blocked with 5% (w/v) NFDM in TBST for 30 min at room temperature. Membranes were washed 3X with TBST and incubated in primary antibody overnight at 4°C with rocking/rotation. Membranes were then washed 3X for 10 min with TBST, incubated with HRP-conjugated secondary antibody in TBST for 1 hour shaking at room temperature, and again washed 3X for 10 min with TBST. Chemiluminescence detection was performed using SuperSignal West Pico PLUS (Thermo Fisher # 34577) and imaged using an Azure Sapphire Biomolecular Imager.

When assaying sucrose gradient fractions, 150 μL of the collected fraction was mixed with 3 μL of 1.4 μΜ recombinant MBP-mEGFP, which served as a spike-in loading control. 47 μL of 4X reducing SDS sample buffer (Bio-Rad # 1610747) was then added, mixed, and heated at 70°C for 15 min. 20 µL (or 40µL for anti-RPS10 blots in Supplementary Data) was then separated by 4-20% Tris-Glycine SDS-PAGE and Western blotting proceeded as described above.

Mouse anti-FLAG M2 (Sigma # F3165-1mg), rabbit anti-V5 (Cell Signaling # 13202; clone D3H8Q), rabbit anti-Ubiquitin (Cell Signaling # 43124; clone E4I2J), rabbit anti-GFP (Cell Signaling # 2956; clone D5.1), mouse anti-alpha (α) Tubulin (Sigma # T9026), rabbit anti-RPS6 (Cell Signaling # 2217; clone 5G10), rabbit anti-RPL7 (abcam # ab72550), and rabbit anti-RPS10 (Abcam # ab151550;EPR8545) were used at 1:1,000 in TBST with 0.02% (w/v) sodium azide. HRP-conjugated goat anti-rabbit IgG (H+L) (Thermo Fisher # 31460) was used at 1:10,000 for anti-V5, -Ubiquitin, and -GFP for Neuro2A cell lysates and anti-RPS10 for sucrose gradient fractions, and 1:30,000 for anti-GFP, -RPS6 and -RPL7 for sucrose gradient fractions. HRP-conjugated goat anti-mouse IgG (H+L) (Thermo Fisher # 31430) was used at 1:10,000.

### RT-qPCR

250 ng of RNA sample extracted from the TOP3B•mRNA activity assay was spiked with 0.2 ng pcDNA3.1(+)/mEGFP plasmid (which served as the internal control for RT-qPCR) and converted to cDNA in 20 µL reactions using iScript Reverse Transcription Supermix with or without reverse transcriptase (+RT and -RT, respectively) (Bio-Rad # 1708841) following the manufacturer’s protocol. Samples were then diluted 10-fold in nuclease-free water and stored at −20°C or used immediately. RT-qPCR was performed in 15 μl reactions using iTaq Universal SYBR Green Supermix (Bio-Rad # 1725124) in a Bio-Rad CFX Connect Real-Time PCR Detection System with 1.5 μl diluted cDNA and 250 nM (final concentration) primers. Relative RNA abundance was calculated between +RT and -RT samples using the spiked-in mEGFP plasmid as the reference gene and the Bio-Rad CFX Maestro software with the standard ΔΔCt method. The following primers were used: F_Mm Actin: 5′-CAGCCTTCCTTCTTGGGTATG-3′; R_Mm Actin 5′-GGCATAGAGGTCTTTACGGATG-3′; F_Mm GAPDH: 5′-AACAGCAACTCCCACTCTTC-3′; R_Mm GAPDH 5′-CCTGTTGCTGTAGCCGTATT-3′; F_Mm Tubulin: 5′-GTGTTCGTAGACCTGGAACC-3′; R_Mm Tubulin 5′-TATTGGCAGCATCCTCCTTG-3′; F_mEGFP: 5′-AGCTGACCCTGAAGTTCATCTG-3′; R_mEGFP 5′-AAGTCGTGCTGCTTCATGTG-3′

### Recombinant protein expression and purification

Insect cell derived recombinant proteins (FLAG-MBP-twinSTII, FLAG-MBP-TOP3B ZnF (G618-S818)-twinSTII, FLAG-MBP-TOP3B ZnF (G618-S818)+C666R-twinSTII, FLAG-MBP-TOP3B ZnF (G618-S818)+D669A-twinSTII, FLAG-MBP-TOP3B ZnF (G618-S818)+ C688A/C691A-twinSTII, His6-TOP3B(WT)-FLAG, and His6-TOP3B(Y336F)-FLAG) were expressed in ExpiSf9 cells using the ExpiSf9 Expression System Starter Kit (ThermoFisher # A39112) following the manufacturer’s protocol. Briefly, bacmids were produced by transforming Max Efficiency DH10Bac Competent Cells (Thermo Fisher # 10361012) with pFastBac plasmids. Integrated transformants were then selected on LB Agar supplemented with 50 µg/mL kanamycin, 7 µg/mL gentamicin, 10 µg/mL tetracycline, 200 µg/mL Bluo-Gal, and 40 µg/mL IPTG for 24h at 37°C. Single integrated transformants (white colonies) were then streaked on new selective LB Agar plates and grown for 24 hr at 37°C. Bacmids were then screened by colony PCR by using 2 µL of overnight LB cultures that were inoculated with single integrated transformants, supplemented with 50 µg/mL kanamycin, 7 µg/mL gentamicin, and 10 µg/mL tetracycline, and incubated for 16 hr at 37°C, 250 rpm. For positive bacmids, 50 µL of the same overnight culture was used to inoculate 100 mL of LB media supplemented with 50 µg/mL kanamycin, 7 µg/mL gentamicin, and 10 µg/mL tetracycline in a 250 mL baffled flask, and then incubated for 16 hr at 37°C, 250 rpm. The culture was split into three equal volumes (∼33 mL), pelleted, and stored at −20°C or immediately used. Bacmids were isolated using the PureLink HiPure Plasmid Midiprep Kit (ThermoFisher # K210004) following the manufacture’s protocol.

To generate P0 baculovirus, 25 mL of ExpiSf9 cells at 2.5 × 10^6^ cells/mL in ExpiSf CD medium in a 125 mL non-baffled vented PETG flask (ThermoFisher # 4115-1000) was transfected with 12.5 µg bacmid using 30 µL ExpiFectamine Sf Transfection Reagent in 1 mL Opti-MEM I Reduced Serum Media and incubated at 27°C, 125 rpm. After 5 days, which typically coincided with ∼60-80% cell death, P0 baculovirus was collected from the ExpiSf9 culture supernatant. Baculovirus was then stored at either 4°C (short term; < 1 week) or aliquoted, snap frozen in liquid nitrogen, and stored at −80°C (long term).

Following the manufacture’s recommendation, 240 mL of ExpiSf9 cells at 5 × 10^6^ cells/mL in ExpiSf CD media in a 1L non-baffled vented PETG flask were treated with 800 µL ExpiSf Enhancer for 22 h at 27°C, 125 rpm. 3 mL of P0 baculovirus stock was then used to transduce the culture for 72-96 hrs at 27°C, 125 rpm. 50 mL culture pellets were then harvested (at 300 x g for 5 min at 4°C), snap frozen in liquid nitrogen, and stored at −80°C.

For purification of insect-derived recombinant proteins, one 50 mL cell pellet was lysed in 16 mL (∼4x cell pellet volume) of ice-cold Lysis Buffer (25 mM HEPES, 500 mM KCl, 10% (v/v) glycerol, 0.5% (v/v) Igepal CA-630, 1 mM DTT; pH 7.5) supplemented with EDTA-free protease inhibitors (Thermo Fisher # A32955), phosphatase inhibitors (Thermo Fisher # A32957), 250U of Pierce Universal Nuclease for Cell Lysis (Thermo Fisher # 88702), and incubated for 15 min at room temperature with gentle end-over-end rotation (15 rpm). Lysates were cleared by centrifugation at ∼215,000 rcf, 4°C for 15 min in S55A rotor using a Sorvall Discovery M120 SE Micro-Ultracentrifuge and added to 1 mL of equilibrated Pierce Anti-DYKDDDDK Affinity Resin slurry (Thermo Fisher # A36803) in a Pierce centrifugation column (Thermo Fisher # 89898) for 2 hrs at 4°C with gentle end-over-end rotation. Columns were then washed with 4 column volumes of ice-cold Wash Buffer (25 mM HEPES, 500 mM KCl, 10% (v/v) glycerol, 0.5% (v/v) Igepal CA-630, 1 mM DTT; pH 7.5) and 1 column volume of room temperature Elution Buffer (25 mM HEPES, 200 mM KCl, 10% (v/v) glycerol; pH 7.5) with centrifugation at 700 x g for 2 min for each wash. FLAG-tagged proteins were eluted with 4 resin-bed volumes of Elution Buffer containing 2.5 mg/mL 3xFLAG peptide (synthesized by Genscript; DYKDHDGDYKDHDIDYKDDDDK) for 15 min at room temperature with gentle end-over-end rotation (15 rpm), and then collected by centrifugation at 700 x g for 2 min.

For twinSTII-tagged proteins, FLAG-eluted protein was then incubated with 0.5 mL of equilibrated Strep-Tactin Superflow Plus slurry (Qiagen # 30004) for 1 h at 4°C with gentle end-over-end rotation (15 rpm). Columns were washed with 2 column volumes of ice-cold Wash Buffer by gravity flow. Recombinant protein was then eluted by incubating 2 resin-bed volumes of Elution Buffer supplemented with 50 mM Biotin (Fisher Scientific # BP2321; 100 mM stock in 100 mM Tris-HCl, pH 8.0) with gentle shaking for 10 min at room temperature, then allowing the eluate to drip by gravity flow.

For His6-tagged proteins, FLAG-eluted protein was then incubated with 0.5 mL of equilibrated HisPur cobalt resin (Thermo Fisher # 89965) for 30 min at 4°C with gentle end-over-end rotation (15 rpm). Columns were washed with 6 resin-bed volumes of ice-cold HisPur Wash Buffer (50 mM sodium phosphate, 300 mM sodium chloride, 10 mM imidazole; pH 7.4) by gravity flow. Recombinant protein was then eluted with 2 resin-bed volumes of HisPur Elution Buffer (50 mM sodium phosphate, 300 mM sodium chloride, 150 mM imidazole; pH 7.4) by gravity flow.

Recombinant protein was then desalted into Elution Buffer (25 mM HEPES, 200 mM KCl, 10% (v/v) glycerol; pH 7.5) using a 5 mL Zeba Spin Desalting Column (7K MWCO for MBP-TOP3B ZnF, 40K MWCO for full-length TOP3B) (Thermo Fisher # 89892, # A57763) following the manufacturer’s protocol. When needed, proteins were concentrated using the Amicon Ultra-2mL Centrifugal Filter (10K MWCO for FLAG-MBP-TOP3B ZnF (G618-S818)-twinSTII or 50K MWCO for His6-TOP3B-FLAG proteins; Millipore # UFC201024 and # UFC205024, respectively) by centrifugation at 3,000 x g for 5 min at room temperature. Recombinant protein concentration was determined by the Pierce Detergent Compatible Bradford Assay Kit (Thermo Fisher # 23246) with BSA standards diluted in Elution Buffer, and purity was analyzed by SDS-PAGE and Coomassie staining. Samples were then aliquoted into single use volumes, snap frozen in liquid nitrogen, and stored at −80°C.

### Inductively coupled plasma mass spectrometry (ICP-MS)

Recombinant proteins in Elution Buffer (25 mM HEPES, 200 mM KCl, 10% (v/v) glycerol; pH 7.5) were aliquoted in metal-free microcentrifuge tubes (Labcon # 3014-870-000-9) and stored at −80°C before being analyzed by ICP-MS at the OHSU Elemental Analysis Core to measure Zn, Fe, Cu, and Mn. 50 µL of each recombinant protein was transferred to a 15 mL metal-free polypropylene tube (VWR # 89049-170) and diluted 1:24 or 1:40 (as indicated) 1% HNO3 (this dilution factor was considered for the final concentration calculations). ICP-MS analysis was performed using an Agilent 8900 triple quad equipped with an SPS autosampler. The system was operated at a radio frequency power of 1550 W, an Ar plasma gas flow rate of 15 L/min, Ar carrier gas flow rate of 0.9 L/min. Elemental concentrations were acquired in kinetic energy discrimination mode (Mn, Fe, Cu, and Zn) using He gas (at 5 mL/min). Data were quantified using weighed, serial dilutions of a multi-element standard (CEM 2, (VHG Labs, VHG-SM70B-100) Mn, Fe, Cu, and Zn). For each biological replicate (n=4 per recombinant protein), data were acquired in triplicates and averaged. A coefficient of variance (CoV) was determined from frequent measurements of a sample containing approximately 10 ppb of Mn, Fe, Cu, and Zn. An internal standard (Sc, Ge, Bi) continuously introduced with the sample was used to correct for detector fluctuations and to monitor plasma stability. Results for the NIST water SRM measured within 95-99%. A *NIST SRM 1683f* was prepared at 8x dilution (3.5 mL of 1% HNO3 and 500 μL of NIST SRM 1683f) to ensure accuracy of the calibration curve. Stoichiometry of each metal to each recombinant protein was calculated by first subtracting the average metal concentration of the Buffer Alone sample from all other samples, then dividing the final metal concentration by the known recombinant protein concentration.

### Structural Modeling and Comparisons

AlphaFold 3 (39) was used for structural prediction of the full length TOP3B using the canonical sequence (UniProtKB: O95985-1). UCSF ChimeraX (40) was used for visualization of cryo-EM density and (Root Mean Square Deviation) RMSD calculations were determined by pairwise alignment of residues in ChimeraX. Structural figures were generated in ChimeraX or PyMol (Schrödinger, Inc.).

### TOP3B Quencher release assay

TOP3B quencher release reactions were prepared in 10 µL volumes by adding 8 µL master mixes containing 100 nM (final) His6-TOP3B(WT)-FLAG or His6-TOP3B(Y336F)-FLAG recombinant protein and 1x Reaction Buffer (5 mM HEPES, 100 mM potassium glutamate, 0.02% (v/v) Tween-20, 0.5 mM MgCl_2_, 2 mM DTT; pH 7.5) to nuclease-free PCR reaction tubes. 2 µL of 100 nM FAM-oligo1DNA-Q (56-FAM/TTTGGGATTATTGAACTGTTGTTCAAGCGTGGT/3IABkFQ) (20 nM final) was then added to the side of the reaction tubes and all tubes were spun down simultaneously to begin the time course. This sequence was previously reported as a favorable *in vitro* TOP3B cleavage substrate (6). A control oligo only reaction containing 1x Reaction Buffer, Elution Buffer (25 mM HEPES, 200 mM KCl, 10% (v/v) glycerol; pH 7.5) instead of recombinant protein, and 20 nM FAM-oligo1DNA-Q was used and only collected at timepoint 0. For timepoint 0 reactions, 20 µL of 2x Quenching Buffer (10 mM EDTA, 0.2% SDS) was added prior to addition of DNA substrate. Reactions were then incubated at 30°C and reactions were quenched by addition of 20 µL 2x Quenching Buffer at 15 sec, 30 sec, 1 min, 2 min, and every minute until 10 min. Upon quenching, reactions were kept in dark at room temperature until all samples were collected. The 10 min timepoint was incubated in dark at room temperature for 5 min after quenching. 20 µL of quenched reactions were then added to a 384-well low flange black flat bottom polystyrene non-binding surface microplate (Corning # 3575) and FAM fluorescence intensity was measured using a Tecan Spark. Rate constants (*k*) were calculated using a non-linear regression, one-phase association with GraphPad Prism.

### Primary cortical neuron culturing and longitudinal fluorescence microscopy

Cortices from embryonic day (E)19-20 Long-Evans rat embryos were dissected and disassociated, and primary neurons were plated at a density of 6×10^5^ cells/mL in 96-well plates or on coverslips. At *in vitro* day (DIV) 4 or 5, neurons were transfected with 100 ng of a control fluorescent plasmid (mApple) to mark cell bodies and 100 ng of mEGFP-NES or mEGFP-NES labeled TOP3B variants using Lipofectamine 2000 as previously described (41–44). Following transfection, cells were cultured in Neurobasal Complete Media (Gibco Neurobasal Medium (Thermo Fisher # 21103049), Gibco 1x B27 Supplement (Thermo Fisher # 17504044), Gibco 1x GlutaMAX Supplement (Thermo Fisher # 35050061), 100 units/mL Penicillin-Streptomycin), and incubated at 37°C in 5% CO_2_.

Neurons were imaged as described previously (43–46) using a Nikon Eclipse Ti inverted microscope with PerfectFocus2, a 20X objective lens, and either an Andor iXon3 897 EMCCD camera or Andor Zyla 4.2 (+) sCMOS camera. A Lambda 421 multi-LED light source (Sutter) with 5 mm liquid light guide (Sutter) was used to illuminate samples, and custom scripts written in Beanshell for use in micromanager controlled all stage movements, shutters, and filters. For automated analyses of primary neuron survival, custom Fiji ImageJ macros and Python scripts were used to identify neurons and draw cellular regions of interest (ROIs) based upon size, morphology, and fluorescence intensity. Custom Python scripts were used to track ROIs over time, and cell death marked a set of criteria that include rounding of the soma, loss of fluorescence and degeneration of neuritic processes (46,47). The ‘survival’ package in R was used to produce Cox proportional hazard baseline plots from the resulting data.

### Generation and maintenance of stable cell lines

Parental Flp-In T-REx HEK 293 cells (ThermoFisher # R78007) were maintained in high glucose DMEM (ThermoFisher # 11995065) supplemented with 10% heat-inactivated Fetal Bovine Serum (Cytiva # SH30396.03HI) and 1% Penicillin-Streptomycin (Thermo Fisher # 15140122), 100 µg/mL Zeocin (ThermoFisher # R25001) and 15 µg/mL Blasticidin S HCL (ThermoFisher # A1113903) at 37°C and 5% CO_2_ in standard tissue culture treated plates. To generate stable cell lines, cells were seeded in a 6-well dish and a single well was transfected using Viafect (Promega # E4981) at a 3:1 ratio with 1 µg total plasmid (500 ng pcDNA5/FRT/TO plasmid + 500 ng pOG44) in 100 µl Opti-MEM. 48 hours post-transfection, cells were split into 10 cm plates with media supplemented with 15 µg/mL Blasticidin S HCl and 100 µg/mL Hygromycin B (ThermoFisher # 10687010). Cells were selected until reaching ∼60-70% confluency with media changes every 3-4 days. After selection, cells were split on a regular schedule with media containing 7.5 µg/mL Blasticidin S HCl and 50 µg/mL Hygromycin B. Stable cell lines were seeded in 12-well plates, without selection drugs, for Western blot analysis and 6-well plates for slot blot analysis and protein expression was induced by adding 1 µg/mL Doxycycline (1 mg/mL stock in water; MP Biomedicals # 195004) for the indicated amount of time. Cells were collected as described above.

### Live-cell imaging of stable cell lines

Flp-In T-REx 293 cells (parent and 3xFLAG-mEGFP-NES-TOP3B) were seeded onto 8-well Nunc Lab-Tek II Chambered Coverglass (ThermoFisher # 55409PK) in high glucose DMEM (Thermo Fisher # 11995065) supplemented with 10% heat-inactivated Fetal Bovine Serum (Cytiva # SH30396.03HI) and 1% Penicillin-Streptomycin (Thermo Fisher # 15140122), without selection drugs. 24 hrs later, media was changed and expression was induced by adding 1 µg/mL Doxycycline. 48 hrs later, cells were stained with 1 µg/mL Hoechst 33342 (ThermoFisher # H3570) in CO_2_ Independent Medium (ThermoFisher # 18045088) supplemented with 10% heat-inactivated Fetal Bovine Serum, 1% Penicillin-Streptomycin, and 20 mM Glutamine (ThermoFisher # A2916801) for 10 min at 37°C and 5% CO_2_. Excess Hoechst 33342 was removed by changing media (complete, supplemented CO_2_ Independent Medium). Confocal imaging was performed using a Nikon Ti2 inverted microscope equipped with an X-Light V3 spinning disk confocal unit (CrestOptics), a Hamamatsu Orca Fusion C14440-20UP detector, a PlanApo 60X water objective with identical laser power and exposure times between −/+ Dox for each genotype. Micrographs were analyzed identically using Fiji ImageJ.

### Sucrose gradient ultracentrifugation

Flp-In T-REx 293 samples were prepared for sucrose gradient ultracentrifugation by seeding cells into 15 cm plates, without selection drugs, to be ∼70-80% confluent on day of collection. To avoid overcrowding and contact inhibition, cells were initially seeded in 10 cm plates, allowed to proliferate for 4 days, and then transferred to 15 cm plates for 3 days. TOP3B expression was induced in the indicated samples for 7 days by addition of 1 µg/mL Doxycycline to cell media, which was changed every 2 days. For control cells treated with anisomycin, media was exchanged with fresh media containing 0.2 µg/mL anisomycin (Sigma # A5862; stock at 10 mg/mL in DMSO) and incubated at 37°C for 15 min prior to cell collection. For collection, cells were placed on a bed of ice and gently washed with 6 mL ice-cold PBS containing 100 µg/mL cycloheximide (Sigma # C1988). With an additional 6 mL ice-cold PBS with 100 µg/mL cycloheximide added to the plate, cells were collected using a cell lifter and then transferred to a pre-chilled 15 mL tube. Cells were pelleted by centrifugation at 500 x g for 5 min in a pre-chilled centrifuge, excess PBS was rigorously removed, and then snap frozen in liquid nitrogen and stored at −80°C.

Cells were lysed in 450 µL Polysome Lysis Buffer (20 mM Tris-HCl, 140 mM KCl, 10 mM MgCl_2_, 1 mM DTT, 1% (v/v) Triton X-100, 100 µg/mL cycloheximide; pH 7.4) and incubated on ice for 10 min. Lysates were cleared by centrifugation at 13,000 rcf for 10 min at 4°C. For RNase treated samples, 0.5 mM CaCl_2_ (final) followed by 150 U of S7 micrococcal nuclease (Roche # 10107921001; stock at 30 U/µL in PBS) was added to cleared lysates, incubated at 25°C for 10 min, and then quenched by adding 1 mM EGTA (final). 400 µL of sample was then loaded onto linear 10-50% buffered sucrose gradients (20 mM Tris-HCl, 140 mM KCl, 10 mM MgCl_2_, 1 mM DTT; pH 7.4) in a 14 mm × 89 mm thin-wall Ultra-Clear tube (Beckman # 344059) that was formed using a BioComp Instruments Gradient Master. Gradients were centrifuged at 35 K rpm for 2 hours at 4°C with max acceleration and no brake in a SW-41Ti rotor using a Beckman Optima L-90 Ultracentrifuge. Gradients were then fractionated into 0.5 mL volumes using a BioComp Instruments piston fractionator with a TRIAX flow cell recording a continuous A260nm trace.

## RESULTS

### The autism-linked C666R mutation causes accumulation of unresolved TOP3B•mRNA covalent intermediates

Three *de novo* TOP3B mutations have been identified in patients diagnosed with intellectual disability, schizophrenia, and autism spectrum disorder (P378Q, R472Q, and C666R, respectively) (35–37). The P378Q and R472Q mutations are within the catalytic core domain, and the C666R mutation is in the zinc finger domain (**Figure 1A**). To our knowledge, only the R472Q and C666R mutations have been previously assessed for their impact on TOP3B, with the latter showing partial loss of topoisomerase activity *in vitro* (9,48). In *TOP3B* null *Drosophila*, the orthologous C666R mutation showed reduced ability to rescue the number of synaptic branches and boutons compared to wildtype (WT), suggesting that TOP3B-C666R is not fully functional (9).

**Figure 1.**
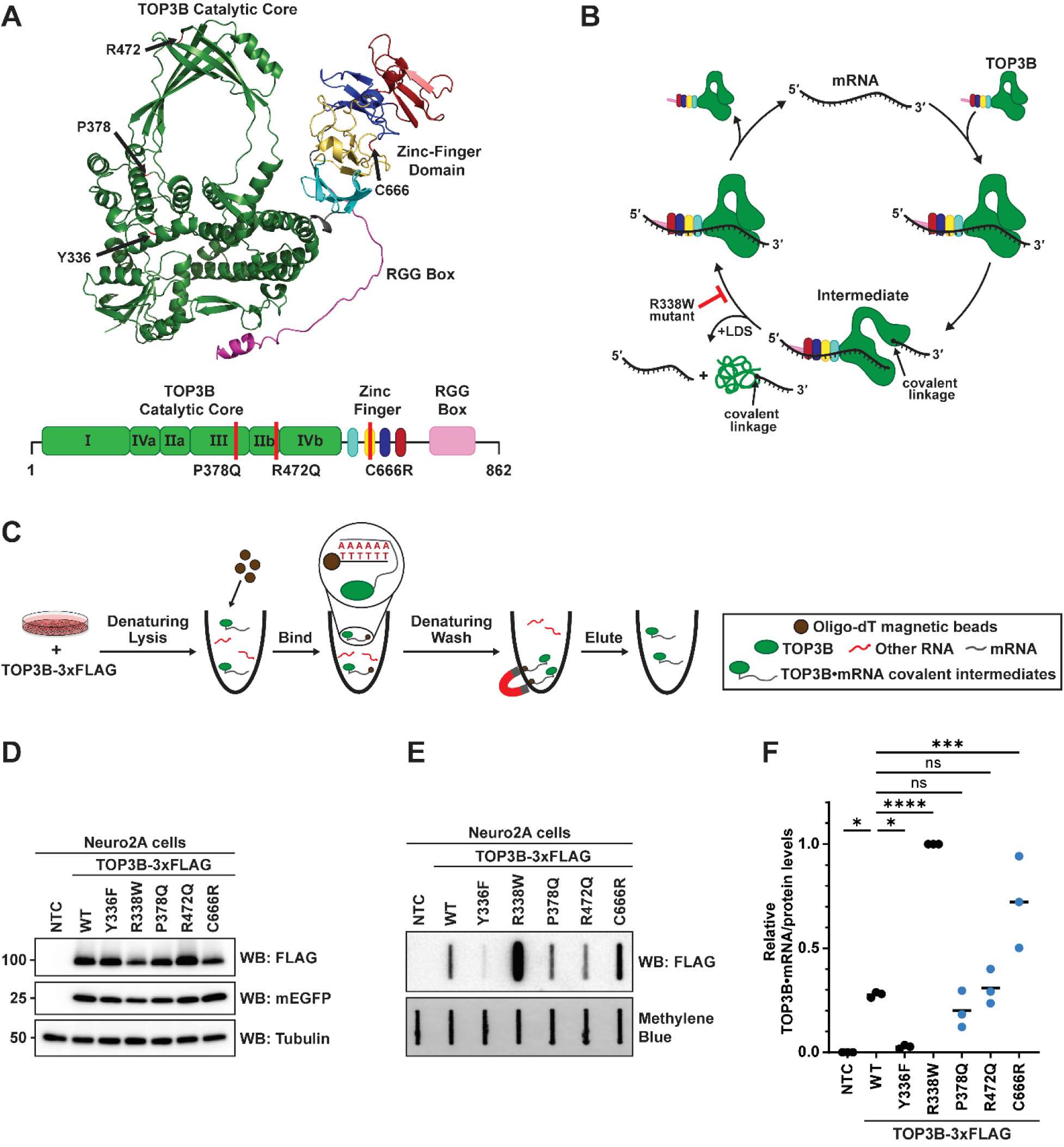
The autism-linked C666R mutation in human TOP3B causes an accumulation of TOP3B•mRNA covalent intermediates. A) AlphaFold predicted structure of full-length human TOP3B (UniProtKB: O95985) with disease-linked mutation residues (P378, R472, and C666) and the catalytic active site tyrosine (Y336) labeled with black arrows (top). Diagram of TOP3B functional domains with disease-linked mutations labeled with red lines (bottom). B) Schematic of TOP3B topoisomerase cycle on mRNA. The synthetic “self-trapping” R338W mutation blocks the mRNA rejoining step, leading to the accumulation of TOP3B•mRNA covalent intermediates. Addition of LDS denatures non-covalently linked TOP3B from substrate mRNAs. C) Schematic of Neuro2A cell-based TOP3B activity assay. Oligo-dT magnetic beads are used to isolate mRNA under strong protein denaturing lysis and wash conditions (*i.e.*, 0.5% (w/v) LDS and 500 mM LiCl) allowing for selective isolation of TOP3B•mRNA covalent intermediates. D) Anti-FLAG Western blot of WT and mutant TOP3B-3xFLAG in Neuro2A cells. mEGFP was used as a transfection control and tubulin was used as a loading control. NTC = no template control. E) Anti-FLAG slot blot of WT and mutant TOP3B•mRNA covalent intermediates isolated from Neuro2A cells (nitrocellulose membrane). Free mRNA was stained with methylene blue (positively charged nylon membrane) and served as a loading control. F) TOP3B activity levels were assessed by quantifying TOP3B•mRNA covalent intermediate levels (signal in panel E) normalized by steady state protein levels (signal in panel D). TOP3B-FLAG protein levels were first normalized by the mEGFP transfection control. Data were then set relative to the R338W mutant. n = 3 biological replicates. Comparisons were made using a one-way ANOVA with Dunnett’s multiple comparisons. * = p<0.05, *** = p≤0.001, **** = p≤0.0001. ns = not significant. Exact p-values are reported in **Supplementary Table S1**.

Unfortunately, published TOP3B activity assays (at least until the time of this publication) are unable to distinguish between the RNA-binding and topoisomerase activity, and do not clearly distinguish the class of RNA being assayed. As most data in the field supports that TOP3B primarily functions in the cytoplasm on mRNA, it is critical to decipher the effect of disease mutations on TOP3B activity on this RNA species. To achieve this, we developed a cell-based TOP3B activity assay that takes advantage of measuring the levels of TOP3B•mRNA covalent intermediates (**Figure 1B,C**). Neuro2A cells were transfected with plasmids encoding WT or mutant TOP3B-3xFLAG, and oligo-dT magnetic beads were used to isolate mRNA under strong protein denaturing lysis and wash conditions (*i.e.*, 0.5% (w/v) LDS and 500 mM LiCl) to allow for selective isolation of TOP3B•mRNA covalent intermediates (**Figure 1C**), which were detected by slot blotting on nitrocellulose membranes and subsequent immunodetection with anti-FLAG. A positively charged nylon membrane was placed underneath to capture free mRNA. Additionally, a plasmid encoding mEGFP was co-transfected as a transfection control when determining steady state protein levels by Western blot.

To confirm that these capture conditions selectively isolated TOP3B•mRNA covalent intermediates over bound but non-covalently linked TOP3B, we included two control mutants. A previously reported catalytically inactive TOP3B mutant, Y336F, that can bind but cannot cleave mRNA was used as a negative control. A synthetic “self-trapping” R338W mutant that prevents substrate rejoining and traps TOP3B in the intermediate state after substrate cleavage (**Figure 1B**) served as a positive control (5,49). As it was previously shown that TOP3B within unresolved covalent intermediates is targeted for degradation by the ubiquitin-proteasome pathway (5), we used protein levels (**Figure 1D**) to normalize the levels of intermediates detected by slot blot (**Figure 1E**) to assess the activity of WT and each mutant (**Figure 1F**).

Both control mutants were detected as expected. Catalytically inactive TOP3B-Y336F produced barely measurable levels of TOP3B•mRNA covalent intermediates, but “self-trapping” TOP3B-R338W robustly produced detectable intermediates. Importantly, including RNase I during oligo-dT mRNA isolation completely abolished the robust signal produced by TOP3B-R338W (**Supplementary Figure S1A**). Additionally, when the final eluate was assayed by RT-qPCR with and without reverse transcriptase, relative levels of actin, GAPDH, and tubulin mRNAs were reduced 2-3 orders of magnitude in the no reverse transcriptase controls (**Supplementary Figure 1B**). Together, these data validate that there is no significant DNA contamination from this cell-based TOP3B activity assay. Lastly, we confirmed the influence of both the RGG box and zinc finger domain using the R338W mutant (**Supplementary Figure S2**). Together, these data demonstrate the ability to selectively assess the catalytic activity of TOP3B on mRNA in cells.

Using this established assay, we next tested the effect of each *de novo* TOP3B mutation compared to WT (**Figure 1D-F**). The P378Q (intellectual disability) and R472Q (schizophrenia) mutants produced similar levels of TOP3B•mRNA covalent intermediates, suggesting the overall TOP3B topoisomerase cycle was unaffected by either mutation. However, a significant increase in TOP3B•mRNA covalent intermediates were detected with the autism-linked C666R mutant, partially phenocopying the “self-trapping” R338W mutant. These data suggest that the C666R mutation may similarly, albeit to a lesser degree, inhibit substrate rejoining and cause accumulation of unresolved covalent intermediates.

In HCT116 cells (human colorectal carcinoma) and HEK293 cells, stable TOP3B covalent intermediates that remain unresolved are reported to be targeted by the E3 ubiquitin (Ub) ligase TRIM41 and subsequently degraded by the proteasome. Tyrosyl DNA phosphodiesterase 2 (TDP2) then cleaves the remaining phosphodiester bond between the substrate RNA and Y336 residue for complete degradation (5) (**Figure 2A**). It should be noted that in this report, the class of RNA being assayed was not identified. Nevertheless, we sought to modulate this targeting system to further validate the C666R mutation partially phenocopying the “self-trapping” R338W mutant. Despite testing multiple antibodies for both TRIM41 and TDP2, we were unable to confirm the expression of either in Neuro2A cells; additionally, the anti-TRIM41 antibody used in previous work is now discontinued. We also attempted to overexpress both effectors individually, but we again were unable to confirm expression using commercially available antibodies. Therefore, we relied on the ability to acutely inhibit the Ub-proteasome pathway at two separate points (**Figure 2A and Supplementary Figure S3A**), similar to previous work with TOP3B. Saha *et al*. reported that inhibitors of the E1 ubiquitin activating enzyme and the proteasome—TAK243 and MG132, respectively—cause accumulation of unresolved TOP3B covalent intermediates produced by the R338W mutation, with a lesser effect on WT TOP3B (5). Both TAK243 (**Figure 2B,C**) and MG132 (**Figure 2D,E**) treatments caused a ∼2.5 fold increase in TOP3B-C666R•mRNA covalent intermediate levels, compared to WT. The impact of both treatments on the R338W mutant was blunted due to the near saturated signal of the untreated condition; however, most replicates do show a small but measurable increase compared to WT.

**Figure 2.**
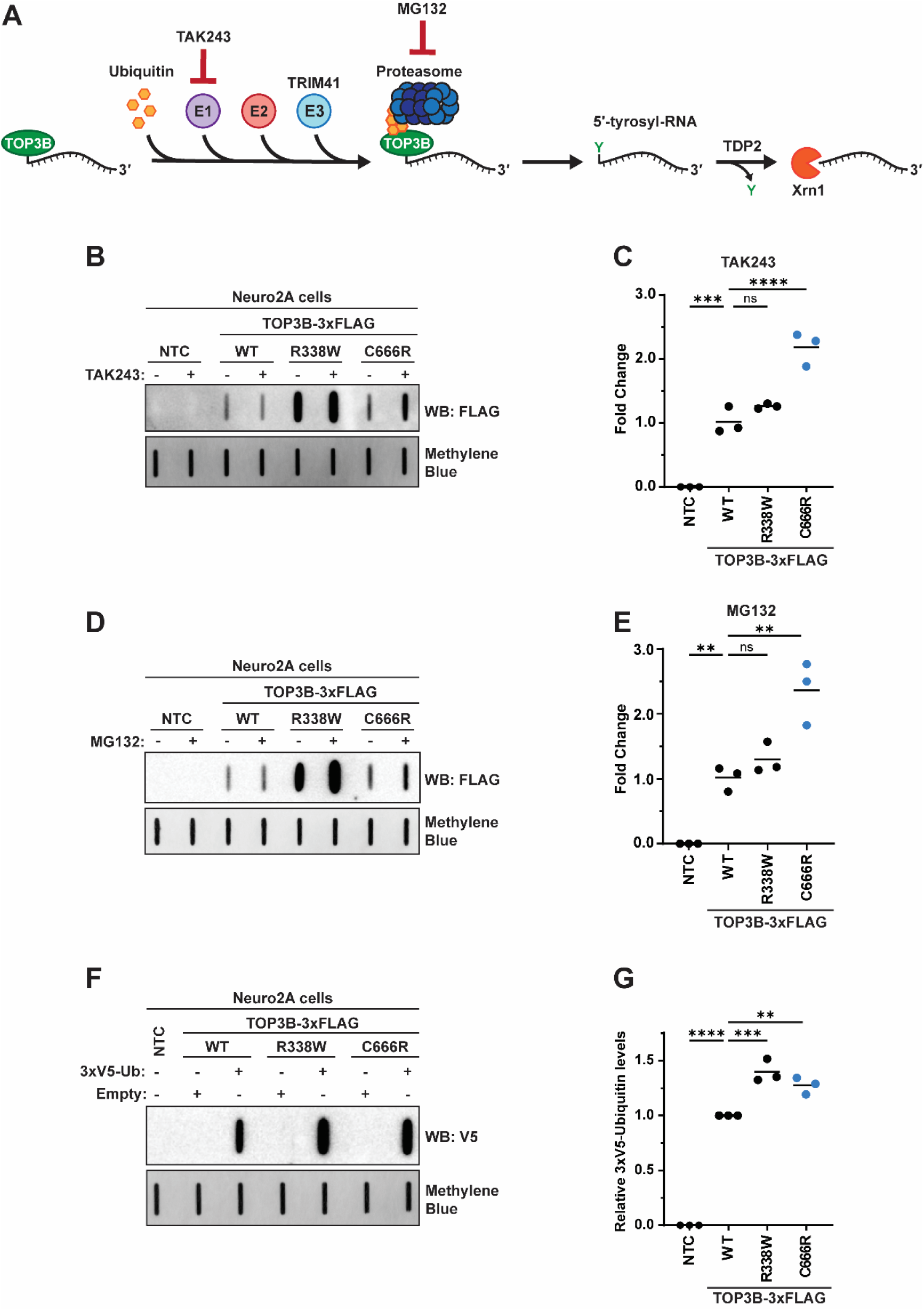
TOP3B-C666R•mRNA covalent intermediates are sensitive to inhibiting the ubiquitin-proteasome pathway, phenocopying the “self-trapping” R338W mutant. A) The reported proteasomal degradation pathway of unresolved TOP3B•RNA covalent intermediates that are sensitive to TAK243 (ubiquitin E1 activating enzyme inhibitor) and MG132 (proteasome inhibitor). B) Anti-FLAG slot blot of WT and mutant TOP3B•mRNA covalent intermediates isolated from Neuro2A cells treated with vehicle or 10 µM TAK243 for 3 hrs. NTC = no template control. C) Quantification of panel B. n = 3 biological replicates. D) Anti-FLAG slot blot of WT and mutant TOP3B•mRNA covalent intermediates isolated from Neuro2A cells treated with vehicle or 10 µM MG132 for 3 hrs. E) Quantification of panel D. n = 3 biological replicates. F) Anti-V5 slot blot of WT and mutant TOP3B•mRNA covalent intermediates isolated from Neuro2A cells co-transfected with either an empty plasmid control (Empty) or a plasmid encoding 3xV5-Ubiquitin (3xV5-Ub). G) Quantification of panel F. n = 3 biological replicates. Free mRNA was stained with methylene blue and served as a loading control for all experiments. Comparisons were made using a one-way ANOVA with Dunnett’s multiple comparisons. ** = p≤0.01, *** = p≤0.001, **** = p≤0.0001. ns = not significant. Exact p-values are reported in **Supplementary Table S2**.

To further validate these findings, we co-expressed 3xV5-Ub with WT and mutant TOP3B and measured the amount of tagged Ub that was detected in the cell-based TOP3B activity assay. Importantly, 3xV5-Ub was functional as it is incorporated into polyubiquitin chains that are sensitive to TAK243 and MG132 (**Supplementary Figure S3B**). 3xV5-Ub was detected at higher levels by slot blot when co-expressed with TOP3B compared to an empty plasmid control (**Supplementary Figure S3C**); this increase in 3xV5-Ub signal was also seen in a mutant that is only capable of making K11- and K48-independent linkages, suggesting WT TOP3B can be tagged with Ub that is not canonically associated with proteasome targeting (**Supplementary Figure S3C**), similar to published data (5). The detection of 3xV5-Ub with empty plasmid (without exogenous TOP3B-WT) is most likely due to endogenous TOP3B. Importantly, our data show that 3xV5-Ub is detected at higher levels when co-expressed with both the R338W and C666R TOP3B mutants compared to WT (**Figure 2F,G**). Together, these data support that the *de novo* C666R mutation results in unresolved TOP3B•mRNA covalent intermediates, partially phenocopying the “self-trapping” R338W mutation.

### The C666R mutation directly interferes with metal coordination within the zinc finger domain

During the majority of our investigation, the exact positioning of the C666 residue within the structure of the zinc finger (ZnF) domain was complicated by two factors. First, there is conflicting literature regarding the predicted number of zinc binding motifs (four vs. five) within the ZnF domain. Based on sequence conservation, it has been proposed that the ZnF domain consists of four cysteine (C4)-type zinc binding motifs, each coordinating a single Zn^2+^ ion (50) (**Figure 3A**). Secondly, at least at the time of this manuscript, an empirically determined structure of full-length TOP3B has not been reported. A solved X-ray crystallography structure of C-terminally truncated TOP3B is available, but the recombinant protein used lacks the ZnF and RGG box domains (48). Therefore, we used AlphaFold to predict the structure of full-length TOP3B with multiple Zn^2+^ ions to investigate how the C666R mutation could affect the ZnF domain (**Figure 3A**). This prediction yielded a structure with very high to high confidence for the catalytic core and ZnF domains, but low confidence for the RGG box domain (**Supplementary Figure 4A**). The AlphaFold 3 prediction shows motifs 1, 3, & 4 each binding a single Zn^2+^ ion with four cysteines (C4) in optimal distance for coordination. The three identified motifs matched with the C4 sequence conservation prediction previously reported. However, we observed disparity in motif 2. Specifically, the reported C4 sequence conservation prediction suggested that residues C688, C691, C706, & C709 make up motif 2; however, AlphaFold predicted that two separate Zn^2+^ are coordinated in this region—one via a D1C3 motif (residues C666, D669, C688, & C691) and one via a C3 motif (residues C706, C709, & C714) (**Figure 3A**)—supporting the presence of five zinc binding motifs within the ZnF domain. Interestingly, this prediction suggests that the C666R mutation would directly interfere with Zn^2+^ coordination and potentially the ZnF domain structure. It should be noted that the crystal structure of full-length *E*. *coli* Topoisomerase I, also a type IA topoisomerase and a homolog of human TOP3B, identified that the orthologous C666 residue (C662 in *E. coli*) is part of a *bona fide* C4-type Zn^2+^ coordination site (51) (**Supplementary Figure S4B,C**). Additionally, during the preparation of this manuscript, a cryo-EM structure using full-length TOP3B (bound to TDRD3, with approximately half of the ZnF domain resolved) was reported (PDB: 9CAH) and confirms the AlphaFold prediction of both the D1C3 and C3 zinc binding motifs (30). Therefore, we next tested the hypothesis that the C666R mutation directly affects metal coordination via two orthogonal approaches.

**Figure 3.**
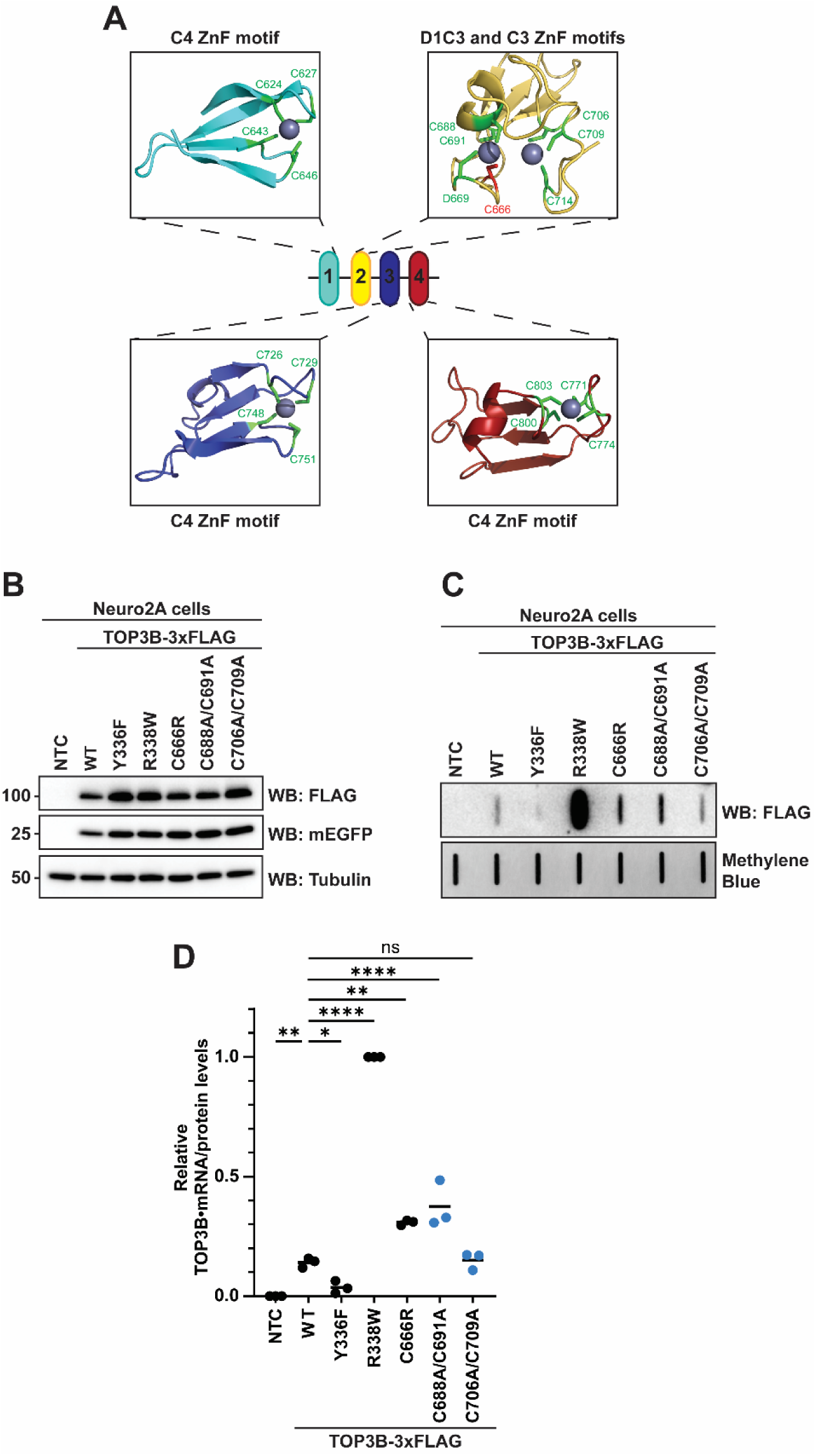
C666R disrupts a D1C3-type metal binding motif within the ZnF domain. A) AlphaFold 3 predicted structure of human TOP3B depicting putative zinc binding motifs and Zn^2+^ coordination (UniProtKB: O95985). The C666 residue is predicted to coordinate Zn^2+^ as part of a D1C3 motif. B) Anti-FLAG Western blot of WT and mutant TOP3B-3xFLAG isolated from Neuro2A cells. mEGFP was used as a transfection control and tubulin was used as a loading control. NTC = no template control. C) Anti-FLAG slot blot of WT and mutant TOP3B•mRNA covalent intermediates isolated from Neuro2A cells. Free mRNA was stained with methylene blue and served as a loading control. D) TOP3B activity levels were assessed by quantifying TOP3B•mRNA covalent intermediate levels (signal in panel C) normalized by steady state protein levels (signal in panel B). TOP3B-3xFLAG protein levels were first normalized by the mEGFP transfection control. Data were then set relative to the R338W mutant. n = 3 biological replicates. Comparisons were made using a one-way ANOVA with Dunnett’s multiple comparisons. * = p<0.05, ** = p≤0.01, **** = p≤0.0001. ns = not significant. Exact p-values are reported in **Supplementary Table S3**.

First, we rationalized that if the C666R mutation was interfering with the role of C666 as part of the D1C3 motif that coordinates a metal, then mutating C688 & C691 (*i.e.*, C688A/C691A) would phenocopy the C666R mutant in the cell-based TOP3B assay. However, if C688, C691, C706, & C709 were truly functioning together within a C4 zinc binding motif, then C688A/C691A and C706A/C709A mutants would have similar phenotypes in the assay. Indeed, and further supporting our hypothesis, the C688A/C691A mutant phenocopied the C666R mutant, causing increased levels of TOP3B•mRNA covalent intermediates compared to WT; the C706A/C709A mutant showed similar levels to WT (**Figure 3B-D**). Additionally, mutating the D669 residue (*i.e.*, D669A) similarly phenocopied the C666R mutant (**Supplementary Figure 5**), supporting the importance of metal coordination by the D1C3 motif in the TOP3B catalytic cycle on mRNA.

Secondly, we measured Zn and other metals bound to the ZnF domain in both WT and mutant contexts. This was achieved by expressing and purifying a tagged version of the TOP3B ZnF domain (residues G618-S818) from insect cells and then measuring stably bound metals by inductively coupled plasma mass spectrometry (ICP-MS). Unfortunately, expression of full-length TOP3B-C666R yielded insoluble protein when expressed at high levels in the baculovirus-insect cell system. Nevertheless, MBP-tagged versions of the ZnF domain were soluble and were able to be purified to very high homogeneity (**Figure 4A,B**). The Tag Alone control (FLAG-MBP-twinSTII) had little detectable Zn, but the WT ZnF domain was measured to have ∼4 Zn ions stably bound per molecule (**Figure 4C**). This stoichiometry was only minimally affected by the C666R mutation (**Figure 4C**). However, upon measuring other metals, we noticed a stark decrease in stably bound Fe. ICP-MS suggests that the WT ZnF domain has ∼1 Fe ion stably bound per molecule, which was essentially eliminated in the C666R ZnF domain (**Figure 4D**). Other metals that are known to bind proteins (*i.e.,* Cu and Mn) were detected at very sub-stochiometric levels close to background levels (**Supplementary Table S5**), suggesting that the Zn and Fe detection is specific to the TOP3B ZnF domain. D669A and C688A/C691A mutations also greatly reduced Fe levels, but they also affected Zn levels more, resulting in ∼3 Zn ions per molecule (**Figure 4C,D**). This could be due to these residues being closer to or part of the more structured region of motif 2 and therefore affecting the folding of the D1C3 motif or downstream metal binding motifs.

**Figure 4.**
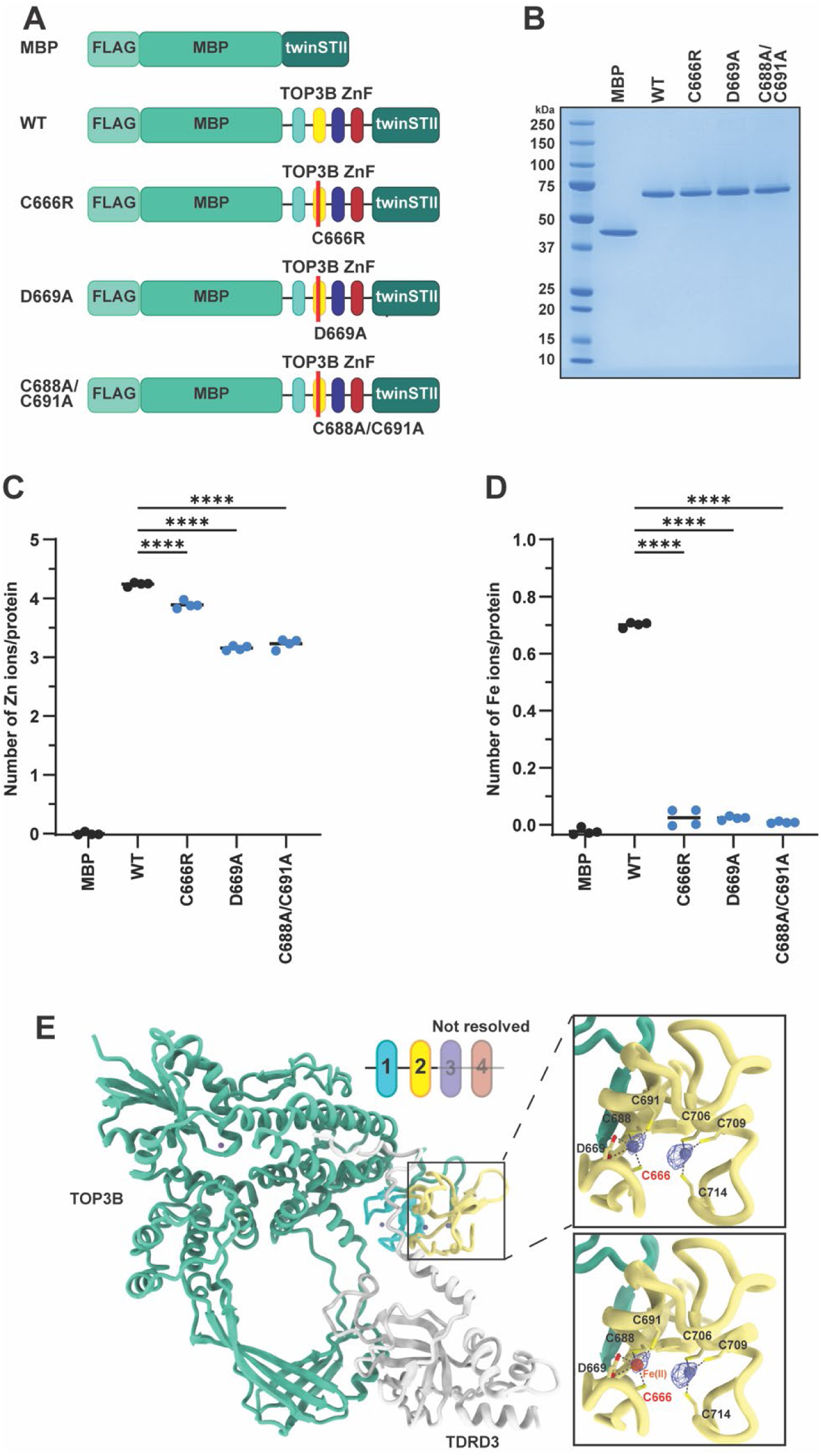
The C666R mutation leads to loss of metal coordination in the ZnF domain. A) Schematic of recombinant FLAG-MBP-twinSTII, FLAG-MBP-ZnF(WT)-twinSTII, and mutant FLAG-MBP-ZnF-twinSTII proteins. B) SDS-PAGE and Coomassie stain of recombinant proteins. C-D) Stoichiometry of Zn (C) and Fe (D) and each indicated recombinant protein as determined by ICP-MS. n = 4 separate ICP-MS samples that were each analyzed in triplicate. E) The published cryo-EM structure of TOP3B bound to TDRD3 (PDB: 9CAH) overlayed with the reported density for the metals in motif 2 with either a Zn (top) or Fe (bottom) being coordinated. Both metals fit into the reported density within the D1C3 motif that consists of C666. Comparisons were made using a one-way ANOVA with Dunnett’s multiple comparisons. **** = p≤0.0001. Exact p-values are reported in **Supplementary Table S4**. ICP-MS data is also found in **Supplementary Table S5**.

Inspection of the published cryo-EM map (EMDB: 45391) shows two clear densities at very high signal to noise, consistent with bound metals, and without additional refinement, we were able to fit both Zn and Fe into said density coordinated by the D1C3 motif that consists of C666 (**Figure 4E**). Additionally, available metal-macromolecule complex validation tools (52,53) equally assess Zn and Fe to be the top ranked metals coordinated by the D1C3 motif in this cryo-EM structure; specifically, the approximated ranking was Zn = Fe = Co > Mn = Ni = Cu > Na = Mg = K >>> Ca. The AlphaFold 3 model and the cryo-EM model (PDB: 9CAH) are highly similar through the core domain of TOP3B (residues 1-606; RMSD = 0.561“Å) but have an all atom RMSD of 9.316 Å due to the flexible nature of the zinc fingers, as predicted by the AlphaFold 3 Predicted Alignment Error (PAE) plot (**Supplementary Figure S4A**). Focusing on the D1C3 motif consisting of C666, the two models are highly similar with near identical arrangements (**Supplementary Figure S4D**).

While five bound metals (4 Zn and 1 Fe) would be consistent with the five predicted metal binding sites from AlphaFold 3, we investigated the possibly that C688, C691, C706, & C709 make up an iron-sulfur cluster that would not have been stabilized during the aerobic conditions of the protein purification. Using both the AlphaFold 3 and the published cryo-EM models, we assessed the distance between the modeled Fe and Zn ions and found that their distance is inconsistent with known Fe-S clusters. Specifically, we kept the modeled Fe coordinated by C666, D669, C688, & C691 and modeled in Fe instead of Zn coordinated by C706, C709, & C714. The distance between the two modeled Fe ions is 5.7 Å. However, the distance between Fe ions in known 2Fe-2S clusters is only ∼2.7-3 Å (*e.g.*, PDB: 6TG9, 2Y5C, and 3P1M). The distance between Fe ions in 4Fe-4S clusters are similar, ∼2.7-3.2 Å (*e.g.*, PDB: 6TG9, 3Q36, and 6NZU). Therefore, all available structural data (prediction and empirically derived) do not support an iron-sulfur cluster in motif 2.

Consistent with ICP-MS results using the MBP-tagged ZnF domain, active full-length WT TOP3B was found to stably bind both Zn and Fe, with Mg and Cu at background levels (**Supplementary Table S6**). Using full-length TOP3B harboring a Y336F (catalytically inactive) mutation as a negative control, we confirmed that full-length WT TOP3B protein was catalytically active in a quencher release assay using a previously published TOP3B substrate (6). In this assay, the substrate has a stem-loop that harbors a 5′ FAM and 3′ Iowa Black FQ quencher that are separated upon TOP3B cleavage, resulting in an increase in FAM fluorescence (**Supplementary Figure S6**). Additionally, the recently published cryo-EM studies of full-length TOP3B obtained reconstructions of TOP3B bound to dsDNA and R-loop substrates; these studies placed the ZnF domain binding upstream of the cleavage site of the dsDNA substrate (30). While the reconstructions were not of a sufficient resolution to build an atomic model, the density ascribed to the ZnF domain clearly moves away from the TOP3B core, consistent with the predicted relative flexibility of this domain, to bind upstream of the cleavage site. The authors thus suggest a role for the ZnF domain in formation and resolution of TOP3B covalent complexes through direct contact with target nucleic acids (30). Together, these data support that the C666R mutation directly disrupts metal coordination within the D1C3 motif of the ZnF domain, thereby impairing the ability of TOP3B to resolve catalytic intermediates.

### Neuronal toxicity is induced by unresolved TOP3B-C666R•mRNA covalent intermediates

Overexpression of the “self-trapping” TOP3B-R338W mutant in HEK293 and HCT116 cells induced cell death and reduced proliferation, with notable signs of DNA damage (5). As TOP3B is primarily cytoplasmic and binds to ORFs of mRNAs, we prioritized testing the effect of accumulating unresolved TOP3B-C666R•mRNA covalent intermediates in neurons. Therefore, we next asked whether TOP3B-C666R, as well as other control mutants, induced neuronal toxicity in primary rat cortical neurons. To mitigate mislocalization to the nucleus upon ectopic expression as much as possible, we generated mEGFP tagged TOP3B constructs that harbored a previously reported and validated nuclear export signal (NES) (54,55) (**Supplementary Figure S7**). Using single cell longitudinal fluorescence microscopy (41–47) (**Figure 5A,B**), we compared the effect of the mEGFP-NES control to mEGFP-NES-TOP3B-WT, -Y336F, -R338W, and -C666R on neuronal survival over a period of 10 days by imaging individual transfected neurons every 24 hrs. While TOP3B-WT did not statistically increase the risk of death compared to the mEGFP control (**Figure 5C**), the catalytically inactive (Y336F) mutant was slightly toxic, indicating a possible dominant negative effect. As expected, due to its ability to generate the highest levels of unresolved TOP3B•mRNA covalent intermediates, primary neurons were most sensitive to the “self-trapping” R338W mutant, as the risk of death for neurons expressing TOP3B-R338W was significantly greater than for those expressing mEGFP-NES control, TOP3B-WT, or TOP3B-Y366F (**Figure 5C**). Consistent with the ability of TOP3B-C666R to partially phenocopy TOP3B-R338W, in addition to the unresolved TOP3B•mRNA covalent intermediates observed with TOP3B-C666R (**Figure 1 and Figure 3**), this mutant also significantly increased the risk of death in primary neurons compared to the mEGFP-NES control (**Figure 5C**). These data highlight a previously-unappreciated neurotoxicity phenotype in association with the autism-linked TOP3B-C666R mutation, suggesting that primary neurons are highly sensitive to unresolved TOP3B-C666R•mRNA covalent intermediates.

**Figure 5.**
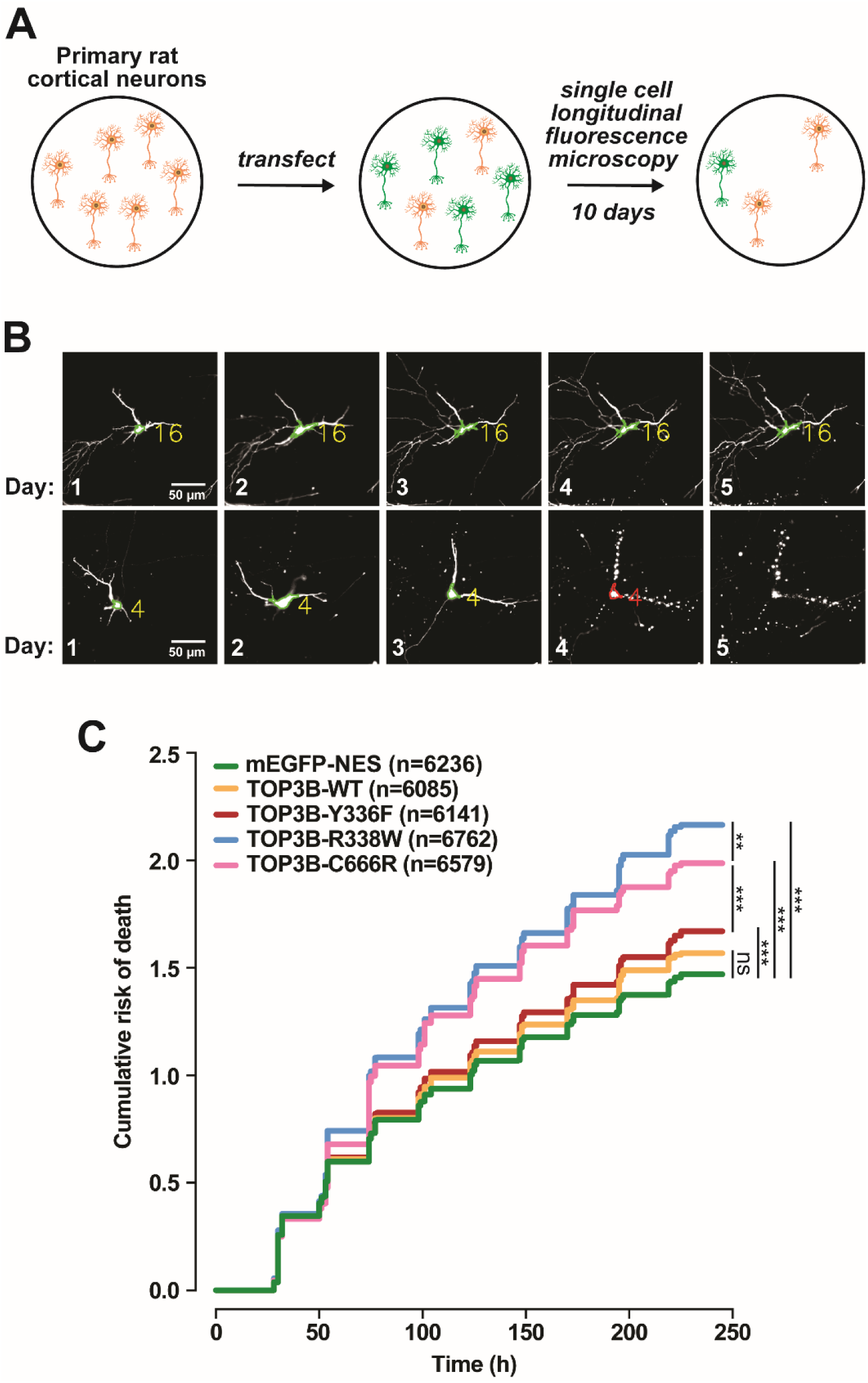
Unresolved TOP3B•mRNA covalent intermediates are toxic to primary neurons. A) Schematic of single cell longitudinal fluorescence microscopy of primary rat cortical neurons. B) Representative images of primary neurons co-transfected with mApple to mark cell bodies (green outline). Transfected primary neurons are imaged taken every 24 hrs; for simplicity, only days 1-5 are shown. Neuron 16 (top) survives the entire time course, whereas neuron 4 (bottom) death can be seen at day 4 (red). C) Cumulative risk of death of primary cortical neurons expressing the mEGFP-NES or the indicated version of mEGFP-NES-TOP3B. Data were combined and stratified from 6 separate experiments imaged every 24 hrs for 10 days. n = total number of neurons imaged. The indicated comparisons were made using a Cox proportional hazard test. *** = p≤0.001. ns = not significant. Exact p-values and all possible comparisons are reported in **Supplementary Table S7**.

### Unresolved TOP3B•mRNA covalent intermediates cause ribosome collisions

The ability of TOP3B to be covalently linked to the mRNA presents a challenge for elongating ribosomes. It is now well established that when an elongating ribosome is selectively inhibited (*e.g.*, translating damaged mRNA, intermediate doses of translation inhibitors) within the ORF, or at the end of a truncated mRNA that lacks a stop codon, it acts as a roadblock for trailing ribosomes and ultimately causes ribosome collisions that potentially stimulate no-go mRNA decay and apoptosis (56–59). As TOP3B primarily binds to ORFs and associates with actively translating polyribosomes, we hypothesized that unresolved TOP3B•mRNA covalent intermediates would directly block ribosomes and cause ribosome collisions (**Figure 6A**). To test this hypothesis, we first generated isogenic stable Flp-In T-REx HEK 293 cell lines that express WT, catalytically inactive Y336F, and “self-trapping” R338W 3xFLAG-mEGFP-NES-TOP3B under doxycycline regulation (**Figure 6B**). The inducible expression system is necessary as prolonged expression of the R338W mutant causes cell death and reduced proliferation in HEK293 and HCT116 cells. Similar to previously described, an NES was included to prevent the accumulation of ectopic TOP3B in the nucleus (**Supplementary Figure S8**). As seen in Neuro2A cells (**Figure 1**), increased TOP3B•mRNA covalent intermediates were observed with TOP3B-R338W compared to WT and the catalytically inactive Y336F control (**Figure 6C**).

**Figure 6.**
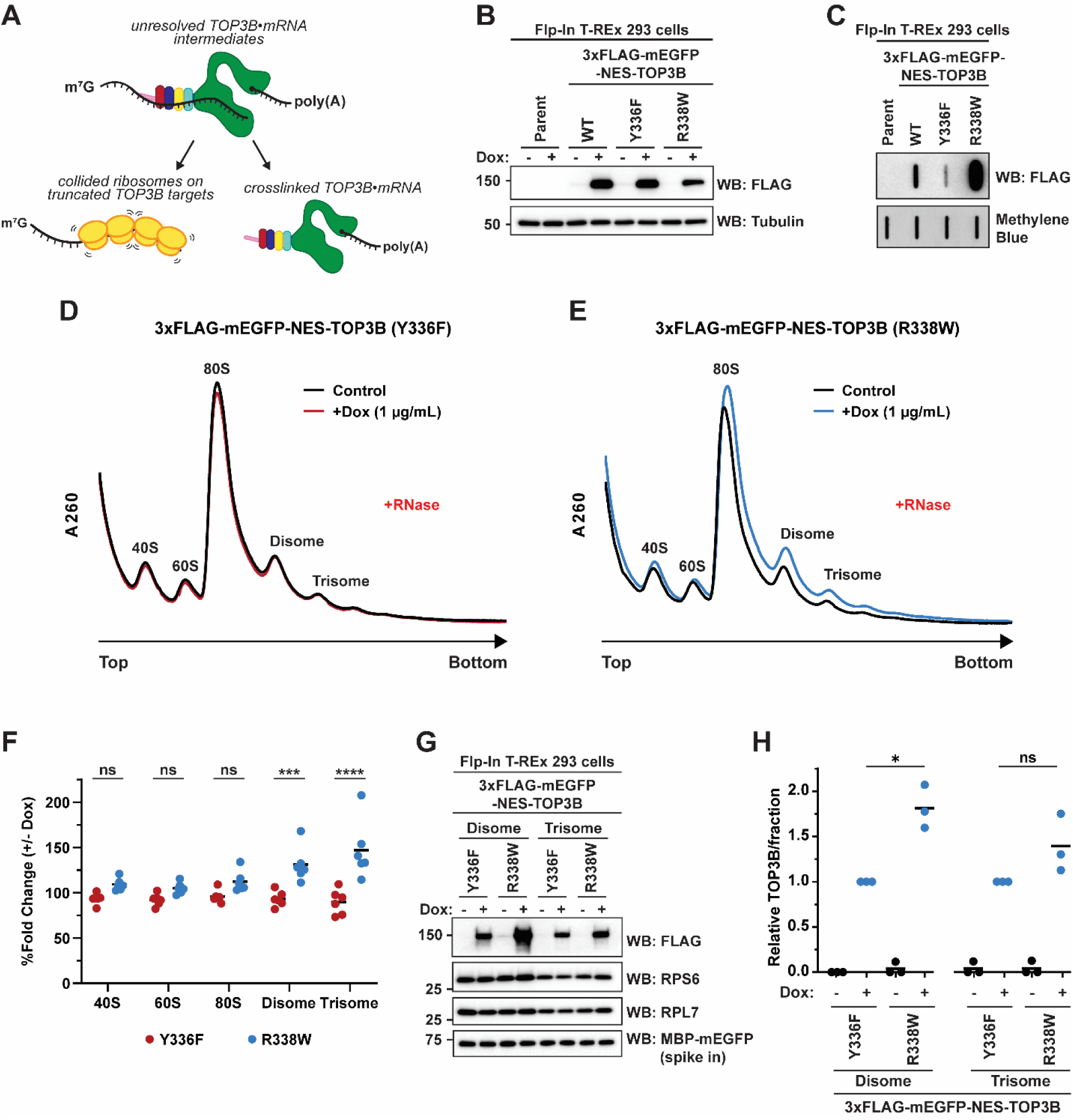
Unresolved TOP3B•mRNA covalent intermediates cause ribosome collisions. A) Schematic of unresolved TOP3B•mRNA covalent intermediates leading to collided ribosomes. B) Anti-FLAG Western blot confirming inducible expression of the indicated Flp-In T-REx 293 cell lines after 24 hrs vehicle or doxycycline (Dox; 1 µg/mL final). Tubulin was used as a loading control. C) Anti-FLAG slot blot of TOP3B•mRNA covalent intermediates isolated from the indicated Flp-In T-REx 293 cell lines after 24 hrs doxycycline (1 µg/mL final). Free mRNA was stained with methylene blue and served as a loading control. D-E) Polysome analysis (10-50% (w/v) sucrose gradients) of the S7 micrococcal nuclease (RNase)-treated lysates of catalytically inactive Y336F mutant (D) or “self-trapping” R338W mutant (E) Flp-In T-REx 293 cell lines after 7-day vehicle or doxycycline (1 µg/mL final). F) Quantification of each ribosomal species reported as the % fold change between + and - Dox induction in the indicated TOP3B mutant. n=6 biological replicates. Comparisons were made using a two-way ANOVA with Sidak’s multiple comparisons. G) Anti-FLAG, anti-RPS6 and anti-RPL7 Western blots of nuclease-resistant disomes (fractions 10) and trisomes (fraction 12). Recombinant MBP-mEGFP was spiked in as a loading control. H) Quantification of panel G 3xFLAG-TOP3B levels were first normalized by RPS6. Data were then set relative to Y336F + Dox in each fraction (disome or trisome). n=3 biological replicates. Comparisons were made using a two-tailed unpaired t-test with Welch’s correction. * = p≤0.05. ns = not significant. Exact p-values for comparisons in panel F and H are reported in **Supplementary Table S8**.

Collided ribosomes have a hallmark of sedimenting within the polysome fractions of sucrose gradients after RNase treatment. Polysomes, mRNAs with multiple bound ribosomes, typically collapse into 80S monosomes upon RNase treatment as the ribonuclease severs the mRNA connection between translating ribosomes. However, collided ribosomes occlude the RNase from gaining access to the mRNA, forming RNase-resistant disomes and trisomes (57,60). We validated this approach and, as expected, a decrease in polysomes and a concatenate increase in 80S monosomes and disomes was observed with RNase treatment (**Supplementary Figure S9A**). Furthermore, addition of an intermediate dose of the translation elongation inhibitor, anisomycin, resulted in the accumulation of RNase-resistant disomes and trisomes compared to the vehicle control (**Supplementary Figure S9B**).

Consistent with our hypothesis that unresolved TOP3B•mRNA covalent intermediates cause ribosome collisions, doxycycline induction of TOP3B-R338W, but not TOP3B-Y336F, resulted in an increase in both nuclease-resistant disomes and trisomes compared to the vehicle control (**Figure 6D-F**). The near indistinguishable signals seen with catalytically inactive TOP3B-Y336F support that the ribosome collisions are not caused by TOP3B binding mRNA, but rather due to the formation of unresolved TOP3B•mRNA covalent intermediates within the ORF. We observed ubiquitylation of RPS10, a secondary hallmark of ribosome collisions (60,61), in the disome and trisome fractions of both cell lines, supporting that the observed accumulation of nuclease-resistant ribosomes are due to ribosome collisions rather than insufficient S7 nuclease treatment (**Supplementary Figure S9C**). Both TOP3B mutants are observed in the disome and trisome gradient fractions by Western blot, suggesting that TOP3B is present at sites of ribosome collisions. Although only statistically significant in the disome fraction, the R338W mutant is seen at higher levels on average than the Y336F mutant in both fractions (**Figure 6G,H**). Together, these data support that unresolved TOP3B•mRNA covalent intermediates cause ribosome collisions.

## DISCUSSION

Locus deletions and *de novo* point mutations linked to neurological disorders highlight the biological importance of encoding a functional TOP3B (6,10–12,62,63). Many of the cellular and behavioral phenotypes connected to cognitive impairment and psychiatric disorders are also observed in multiple animal models harboring TOP3B deletion or mutations (7,20). However, the fundamental function of TOP3B that is affected which ultimately contributes to disease is still unclear. Answering this question is further complicated by the ability of TOP3B to be catalytically active on DNA and RNA, as well as by the atypical predominant cytoplasmic localization for a topoisomerase. Several observations have been reported pointing to the primary role of TOP3B being in the cytoplasm on translating mRNA (6,7,34). The strong bias of TOP3B binding to the open reading frame, as observed in CLIP data sets, strongly supports a role in regulating translation and is uncharacteristic of an RNA-binding protein that would be involved in mRNA metabolism in the nucleus. However, transcriptional, R-loop, and genomic instability changes have been noted in cells deficient of TOP3B (5,20,26–29), raising the possibility that pathology could at least be partially influenced or tied to these changes.

Disease linked mutations can be informative for identifying critical residues or domains of the mutated gene’s functional product. In this report, we investigated three TOP3B mutations—P378Q, R472Q, C666R—and find that the C666R interferes with metal coordination within the ZnF domain, resulting in the increased abundance of TOP3B•mRNA covalent intermediates. Our investigation using a combination of genetic and biochemical approaches interrogating the AlphaFold 3 predicted structure of TOP3B shows that the ZnF domain was previously mischaracterized when using C4-type zinc binding motif predictions based on the primary sequence (**Figure 3,4**), in alignment with a recent cryo-EM structure (30).

A critical aspect of completing a full TOP3B topoisomerase reaction is the realignment of the cleaved substrate in the active site after strand passage, to allow for reverse phosphoryl transfer to rejoin the 3′ OH and 5′ scissile phosphate of the cleaved mRNA (32,33,64). Multiple active site residues can influence this proper realignment, including E9, K10, D117, D119, and R338 (25,30,48). As recent work has shown that the ZnF domain directly binds a DNA bubble and R-loop substrate *in vitro* (30), it is possible that the C666R mutated ZnF domain affects the proper realignment of the cleaved substrate—resulting in the observed increase in covalent intermediates. However, the exact position of the complete ZnF domain unbound and bound to substrate still remains to be reported. Recent cryo-EM studies were only able to confidently resolve approximately the first half of the ZnF domain and provided evidence of two separate metals being coordinated by the D1C3-type motif (formed partially by C666) and the C3-type motif that were predicted by AlphaFold (30) (**Figure 3**). In these structural studies, the ZnF domain begins interacting with the substrate only ∼9 nt from the scissile phosphate in the active site. The adjacent position of the ZnF domain to the catalytic core may explain why C666R only partially phenocopies the “self-trapping” R338W mutation that is located in the active site.

Residues that correspond to disease-causing missense mutations are typically highly conserved (65). However, for the three *de novo* TOP3B point mutations, only P378 and C666 are conserved from *Drosophila* to humans. R472 is highly variable in metazoans (50) and the schizophrenia-linked R472Q mutant having no observable effect in our cell-based TOP3B activity assay is consistent with other reports (9). Ahmed *et al*. demonstrate no difference in protein expression or mRNA binding in cells, and recombinant TOP3B-R472Q had similar topoisomerase activity *in vitro* compared to WT (9). Goto-Ito *et al.* also showed that the R472Q mutation did not affect the tertiary structure of TOP3B (48). As TOP3B has been shown to also regulate mRNA metabolism independent of topoisomerase activity (Y336F mutation) (34), it cannot be ruled out that these mutations also affect this facet of TOP3B-mediated regulation, as well as its reported functions in the nucleus.

*TOP3B* locus deletions are the most commonly reported mutation (6,10–12). As strong data supports that TOP3B is catalytically functional on both DNA and RNA in cells (5–7,25), a remaining key question for the field is identifying whether pathology is driven more by the loss of activity on one substrate over another. Given its predominant cytoplasmic localization, it may be initially presumed that changes in transcription and chromatin are secondary or minor to the altered mRNA metabolism in the cytoplasm. Animal models harboring RGG box domain deletions to robustly reduce RNA targeting, as well as identify the most TOP3B-regulated targets in neurons, may prove to be powerful in addressing this gap. Indeed, ectopic expression of an RGG box deletion mutant in TOP3B null flies was unable to rescue the observed abnormal neuromuscular junction phenotype (9), suggesting that loss of activity on RNA may more heavily contribute to disease phenotypes. Nevertheless, the ability of TOP3B to function on RNA presents an exciting and biologically important facet of RNA biology.

### Limitations of this study

Due to low detection limits of endogenous TOP3B•mRNA covalent intermediates and reported cell viability issues with long-term expression of TOP3B mutants that cause stable covalent intermediates (5), our studies use ectopic overexpression of WT and mutant TOP3B. We also acknowledge that the metal binding studies were completed using only the ZnF domain of TOP3B and not full-length recombinant TOP3B-C666R protein; this was due to the inability to recover many full-length TOP3B mutants. Additionally, all recombinant proteins were purified under aerobic conditions that do not robustly stabilize iron-sulfur clusters, but follows similar aerobic purification conditions that were used for recombinant TOP3B protein preparation and cryo-EM by other groups (6,7,25,30).

## DATA AVAILABILITY

The data underlying this article are available in the article and in its Supplementary Data.

## ACKNOWLEDGEMENTS

We thank members of the Kearse lab, Dr. Venkat Gopalan, Dr. Karin Musier-Forsyth, and Dr. Kurt Fredrick for stimulating conversations and input. We also thank Dr. Martina Ralle and Ms. Sophia Miller for their input and expertise with ICP-MS. The Ohio State University Comprehensive Cancer Center Genomics Shared Resource (OSUCCC GSR) is supported by NIH grant P30CA016058. ICP-MS measurements were performed in the Oregon Health and Science University Elemental Analysis Core with partial support from NIH grant S10OD028492.

