## Supplementary Data for "An autism spectrum disorder mutation in Topoisomerase 3β causes accumulation of covalent mRNA intermediates by disrupting metal binding within the zinc finger domain"

### SUPPLEMENTARY FIGURES

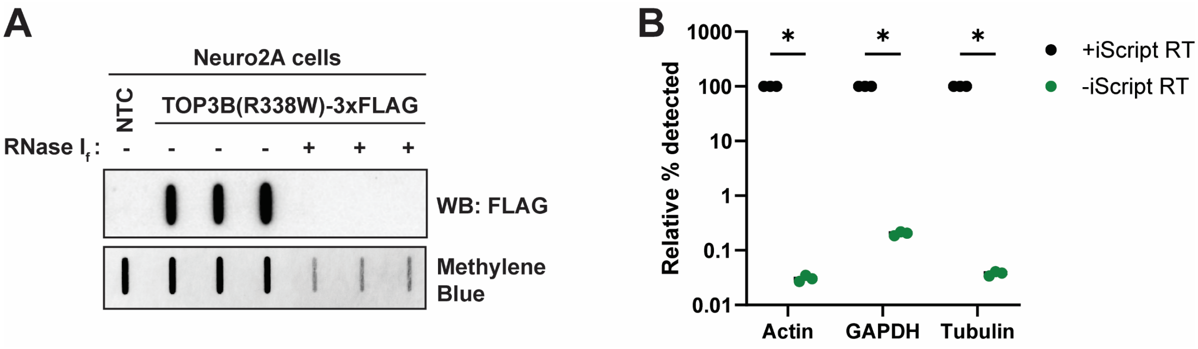


**Supplementary Figure S1. Cell-based TOP3B activity assay is highly sensitive to RNase and is not contaminated with DNA.** A) Anti-FLAG slot blot of TOP3B-R338W•mRNA covalent intermediates isolated from Neuro2A cells via oligo-dT magnetic beads under stringent conditions (**Figure 1C**) without and with addition of RNase $I_f$ during the last wash (see methods for more details) (nitrocellulose membrane). Free mRNA was stained with methylene blue (positively charged nylon membrane) and served as a loading control. The decrease in methylene blue staining in the +RNase $I_f$ lanes is expected due to the RNase activity during isolation; equal volume of eluates were assessed (see methods). B) Relative quantification by RT-qPCR of Actin, GAPDH, and Tubulin mRNA with and without iScript Reverse Transcriptase (RT) added during cDNA synthesis. 0.2 ng of mEGFP plasmid was added prior to cDNA synthesis and used as reference gene that was independent of RT. n = 3 biological replicates. Comparisons were made using an unpaired t test with Welch correction. * = p<0.000001. Exact p-values are reported in **Supplementary Table S9**.

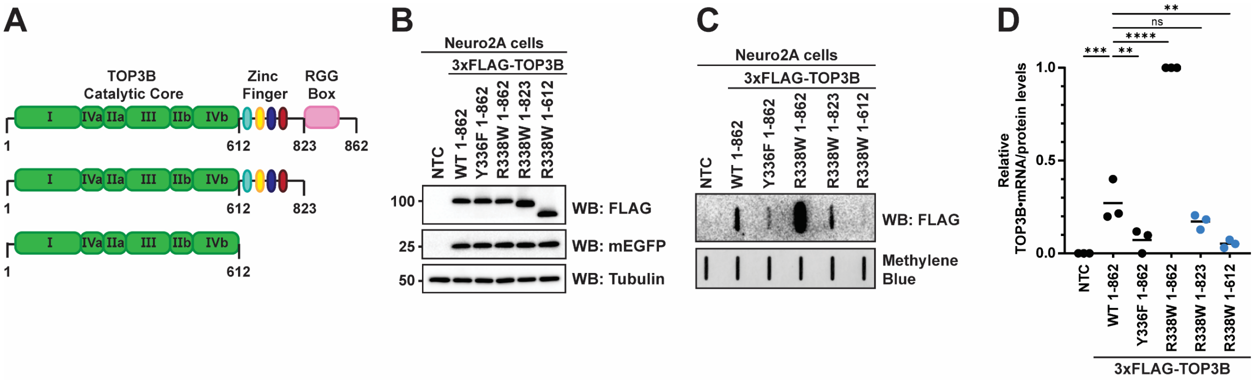


**Supplementary Figure S2. Deletion of C-terminal domains affects the accumulation of TOP3B-R338W•mRNA covalent intermediates.** A) Diagram of full-length and mutant TOP3B harboring domain deletions used. B) Anti-FLAG Western blot of WT and mutant 3xFLAG-TOP3B in Neuro2A cells. mEGFP was used as a transfection control and tubulin was used as a loading control. NTC = no template control. C) Anti-FLAG slot blot of WT and mutant TOP3B•mRNA covalent intermediates isolated from Neuro2A cells (nitrocellulose membrane). Free mRNA was stained with methylene blue (positively charged nylon membrane) and served as a loading control. D) TOP3B activity levels were assessed by quantifying TOP3B•mRNA covalent intermediate levels (signal in panel C) normalized by steady state protein levels (signal in panel B). 3xFLAG-TOP3B protein levels were first normalized by the mEGFP transfection control. Data were then set relative to the R338W mutant. n = 3 biological replicates. Comparisons were made using a one-way ANOVA with Dunnett's multiple comparisons. ** = $p<0.01$, *** = $p\leq0.001$, **** = $p\leq0.0001$. ns = not significant. Exact p-values are reported in **Supplementary Table S9**.

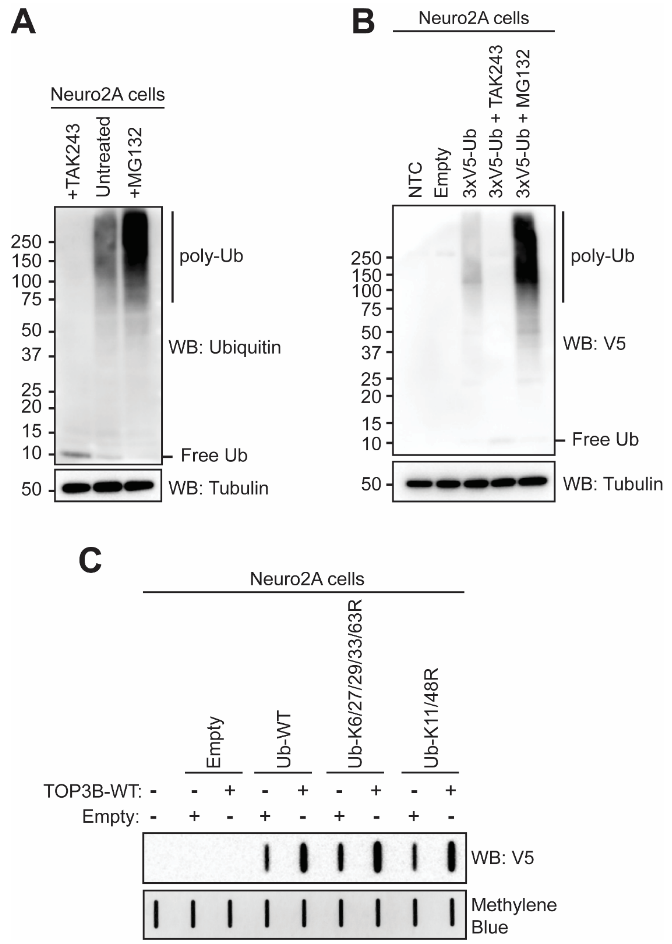


**Supplementary Figure S3. 3xV5-Ubiquitin is functional and incorporated into polyubiquitin chains.** A) Anti-Ubiquitin (Ub) Western blot of Neuro2A cells treated with 10 μM TAK243 or MG132 for 2 hrs. Tubulin was used as a loading control. B) Anti-V5 Western blot from Neuro2A cell lysates transfected with an empty plasmid or a plasmid encoding 3xV5-Ub for 24 hrs and then left untreated or treated with 10 μM TAK243 or MG132 for 2 hrs. Tubulin was used as a loading control. C) Anti-V5 slot blot detecting TOP3B•mRNA covalent intermediates isolated from Neuro2A cells transfected with an empty plasmid control (Empty) and or a plasmid encoding WT TOP3B-3xFLAG. An empty plasmid control (Empty) or a plasmid encoding WT or mutant 3xV5-Ubiquitin (3xV5-Ub) was co-expressed. Free mRNA was stained with methylene blue and served as a loading control.

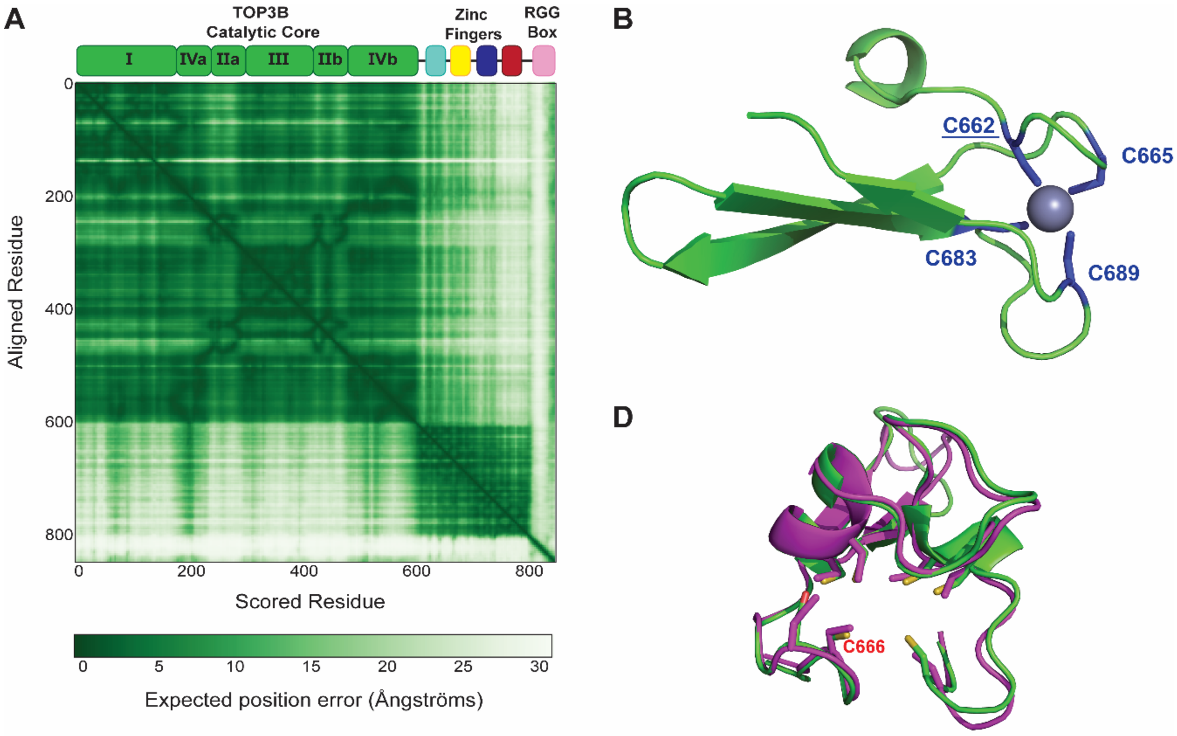


C

```
Hs TOP3B   1    MKTVLMVAEKPSLAQSIAKILSRGSLSSHKGLNGACSVHEYTGTFAGQPVRFKMTSVCGHVMTLDFLGKYNKWDKVDPAE  80
Ec TopoI   1    MGKALVIVESPAKAKTINKYLGSD---------------------------YVVKSSVGHIRDLPTSGSAAKKSADSTST  53

Hs TOP3B   81   LFSQAPTEKKEA--------NP-------------KLNMVKFLQVEGRGCDYIVLWLDCDKEGENICFEVLDAVLPVMNK  139
Ec TopoI   54   KTAKKPKKDERGALVNRMGVDPWHNWEAHYEVLPGKEKVVSELKQLAEKADHIYLATDLDREGEAIAWHLREVI------  127

Hs TOP3B   140  AHGGEKTVFRARFSSITDTDICNAMACLGEPDHNEALSVDARQELDLRIGCAFTRFQTKYFQGKYGDLDSSLISFGPCQT  219
Ec TopoI   128  -GGDDARYSRVVFNEITKNAIRQAFNKPGELNIDRVNAQQARRFMDRVVGYMVSPLLWKKI--------ARGLSAGRVQS  198

Hs TOP3B   220  PTLGFCVERHDKIQSFKPETYWVLQAKVNTDKDRSLLLD-----WDRVRVFDREIAQMFLNMTKLEKEAQVEATSRKEKA  294
Ec TopoI   199  VAVRLVVEREREIKAFVPEEFWEVDASTTTPSGEALALQVTHQNDKPFRPVNKEQTQAAVSLLE-KARYSVLEREDKPTT  277

Hs TOP3B   295  KQRPLALNTVEMLRVASSSLGMGPQHAMQTAERLYTQGYISYPRTETTHYPENFDLKGSLRQQANHPYWADTVKRLLAEG  374
Ec TopoI   278  SKPGAPFITSTLQQAASTRLGFGVKKTMMMAQRLYEAGYITYMRTDSTNLSQDAVNMVRGYISDNFG------KKYLPES  351

Hs TOP3B   375  INR-PRKGHDAGDHPPITPMKSATEAE----LGGDAWRLYEYITRHFIATVSHDCKYLQSTISFRIGPELFTCSGKTVLS  449
Ec TopoI   352  PNQYASKENSQEAHEAIRPSDVNVMAESLKDMEADAQKLYQLIWRQFVACQMTPAKYDSTTLTVGAGDFRLKARGRILRF  431

Hs TOP3B   450  PGFTEVMPWQSVPLEES-LPTCQRGDAFPVGEVKMLEKQTNPPDYLTEAELITLMEKHGIGTDASIPVHINNICQRNYVT  528
Ec TopoI   432  DGWTKVMPALRKGDEDRILPAVNKGDALTLVELTPAQHFTKPPARFSEASLVKELEKRGIGRPSTYASIISTIQDRGYVR  511

Hs TOP3B   529  VESGRRLKPTNLGIVLVHGYYKIDAELVLPTIRSAVEKQLNLIAQGKADYRQVLGHTLDVFKRKFHYFVDSIAGMDELME  608
Ec TopoI   512  VEN-RRFYAEKMGEIVTDRLEENFRELMNYDFTAQMENSLDQVANHEAEWKAVLDHFFSDFTQQ----LDKAEKDPEEGG  586

Hs TOP3B   609  VSFSPLAATGKPLSRCGKCHRFMKYIQAKPSRLHCS--------HCDETYTL-PQNGTIKLYK-----------ELRCPL  668
Ec TopoI   587  MRPNQMVLTSIDCPTCGR--KMGIRTASTGVFLGCSGYALPPKERCKTTINLVPENEVLNVLEGEDAETNALRAKRRCP-  663

Hs TOP3B   669  DDFELVLWSSGSRGKSYPLCP-----YCYNHPP----------FRDMKKGMGCNECTHPSCQHSLSMLGIGQCVECESGV  733
Ec TopoI   664  --------KCGTAMDSYLIDPKRKLHVCGNNPTCDGYEIEEGEFRIKGYDGPIVECEKCGSEMHLKMGRFGKYMACTNEE  735

Hs TOP3B   734  LVLDPTSGPKWKVACNKCNVVAH----CFENAHRVRVSADTCSVCEAALLDVDFNKAKSPLPGDETQHMGCVFCDPVFQE  809
Ec TopoI   736  CKNTRKILRNGEVAPPKEDPVPLPELPCEKSDAYFVLRDGAAGVFLAANTFPKSRETRAPLVEELYR-----FRDRLPEK  810

Hs TOP3B   810  LVELKHAASCHPMHRGGPGRRQGRGRGRARRPPGKPNPRRPKD-KMSALAAYFV----------  862
Ec TopoI   811  LRYLADAPQQDP--------EGNKTMVRFSRKTKQQYVSSEKDGKATGWSAFYVDGKWVEGKK  865
```

**Supplementary Figure S4. Assessment of the AlphaFold predicted structure of TOP3B and conservation of the C666 residue within a metal binding motif.** A) Predicted Alignment Error (PAE) plot for the predicted structure of TOP3B (UniProtKB: O95985) shows high accuracy (low error) for the TOP3B catalytic core region. Zinc fingers show low positional accuracy relative to each other but high optional error relative to the catalytic core, and the RGG motif shows low positional accuracy. B-C) The homologous C666 residue in human TOP3B is conserved in *E. coli* TopoI within a *bona fide* C4-type zinc binding motif. The solved X-ray crystallography structure of full-length *E. coli* TopoI (PDB: 4RUL) demonstrates that C662 (underlined) contributes to a C4-type zinc finger motif; for simplicity, only residues 655-696 are shown (B). Alignment of human TOP3B and *E. coli* TopoI using COBALT (Constraint-based Multiple Alignment Tool) from NCBI with sequence identity shown in red (C). The Human TOP3B (Hs TOP3B) sequence is from NCBI Reference Sequence NP_003926.1 and the *E. coli* TopoI (Ec TopoI) sequence is from NCBI Reference Sequence NP_003926.1. The C666 residue in human TOP3B and the homologous C662 residue in *E. coli* TopoI are underlined. D) Pymol structural alignment of the AlphaFold 3 prediction (green) and the published cryo-EM structure (magenta) for TOP3B (PDB: 9CAH); only residues 663-713 are shown for simplicity and to compare the D1C3 motif that consists of C666 (labeled in red as a reference point).

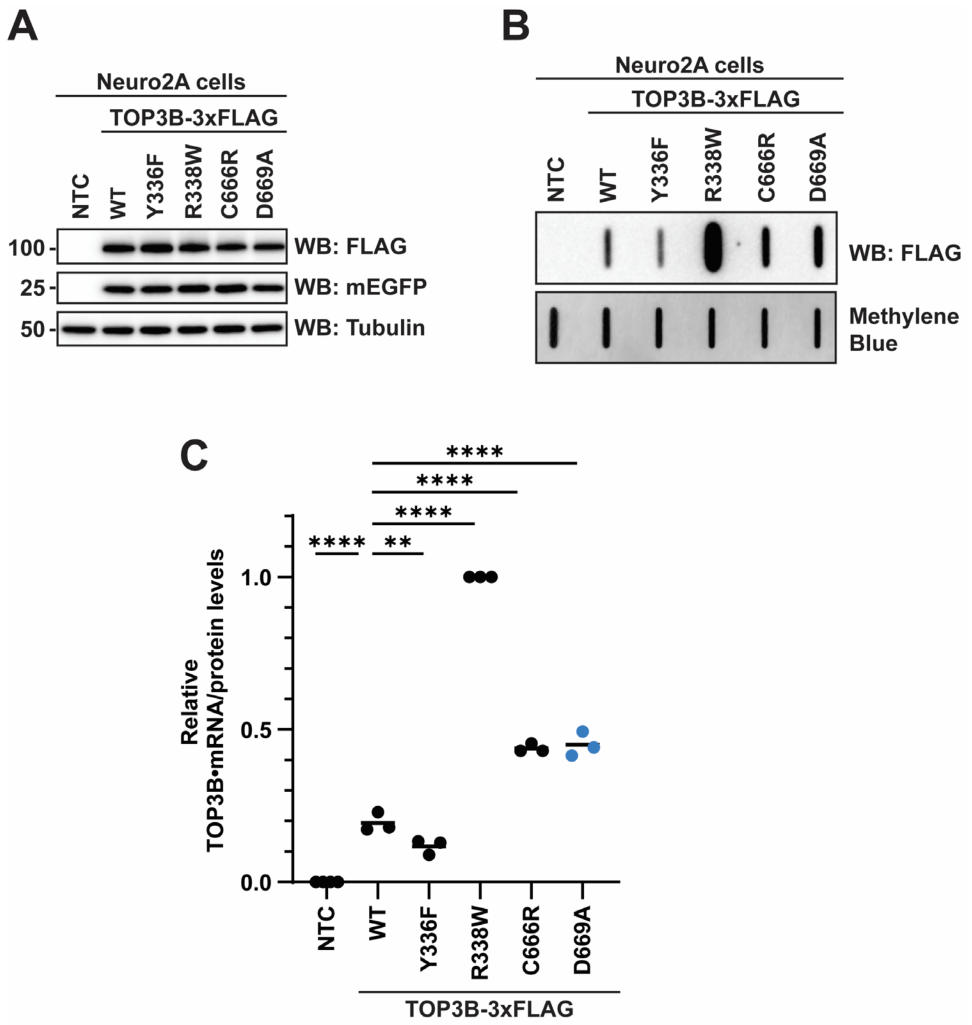


**Supplementary Figure S5. Mutating the D669 residue within the D1C3 motif phenocopies the C666R mutation.** A) Anti-FLAG Western blot of WT and mutant TOP3B-3xFLAG in Neuro2A cells. mEGFP was used as a transfection control and tubulin was used as a loading control. NTC = no template control. B) Anti-FLAG slot blot of WT and mutant TOP3B•mRNA covalent intermediates isolated from Neuro2A cells (nitrocellulose membrane). Free mRNA was stained with methylene blue (positively charged nylon membrane) and served as a loading control. C) TOP3B activity levels were assessed by quantifying TOP3B•mRNA covalent intermediate levels (signal in panel B) normalized by steady state protein levels (signal in panel A). TOP3B-3xFLAG protein levels were first normalized by the mEGFP transfection control. Data were then set relative to the R338W mutant. n = 3 biological replicates. Comparisons were made using a one-way ANOVA with Dunnett's multiple comparisons. ** = $p<0.01$, **** = $p \leq 0.0001$. Exact p-values are reported in **Supplementary Table S9**.

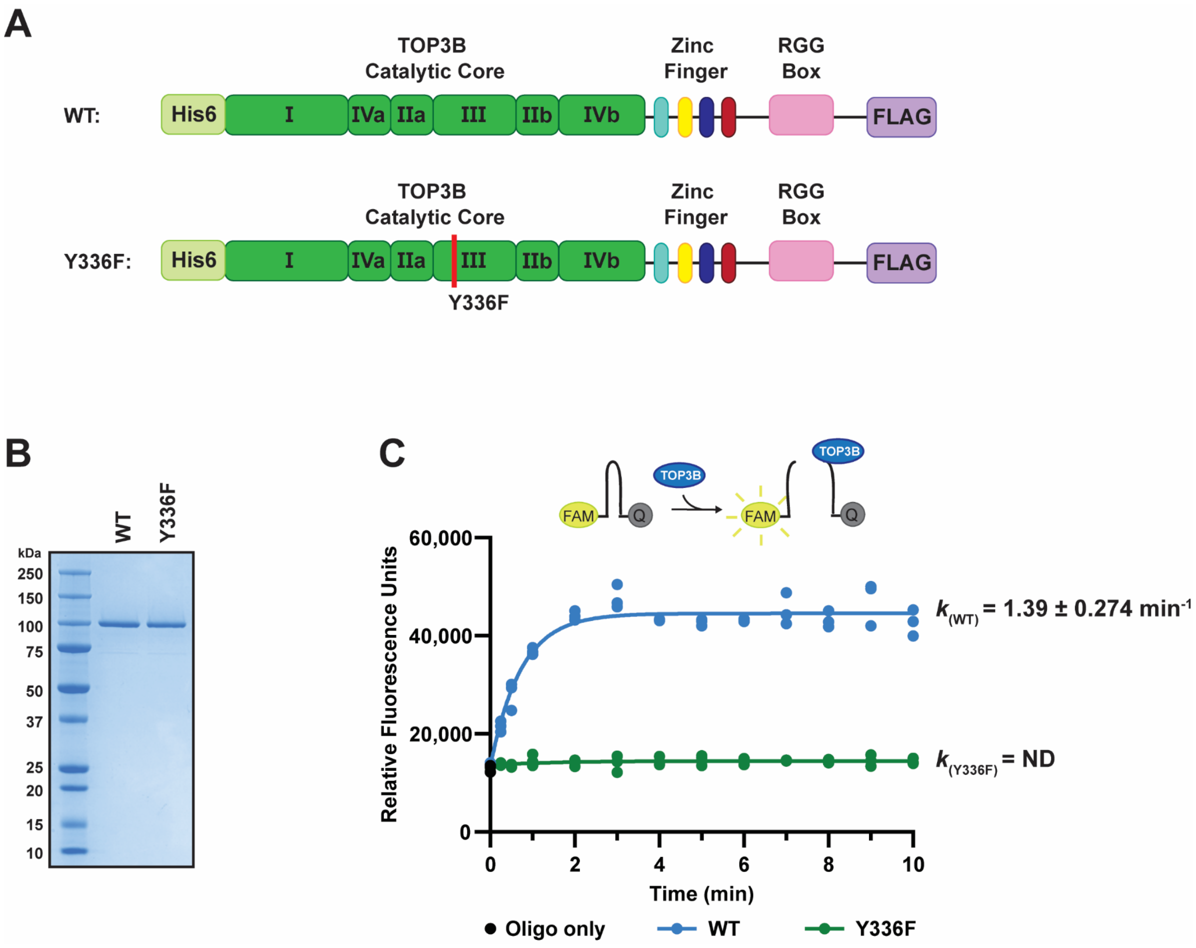


**Supplementary Figure S6. Recombinant full-length TOP3B is catalytically active.** A) Schematic of recombinant full-length His6-TOP3B(WT)-FLAG and His6-TOP3B(Y336F)-FLAG expressed and purified using a baculovirus-insect cell system. B) SDS-PAGE and Coomassie stain of the indicated recombinant full-length TOP3B. C) Diagram of quencher release assay demonstrating that catalytic cleavage of stem-loop DNA substrate by TOP3B separates the quencher and fluorophore to allow FAM fluorescence (top). Quencher release assay demonstrating recombinant TOP3B-WT is catalytically active (bottom). n = 3 biological replicates.

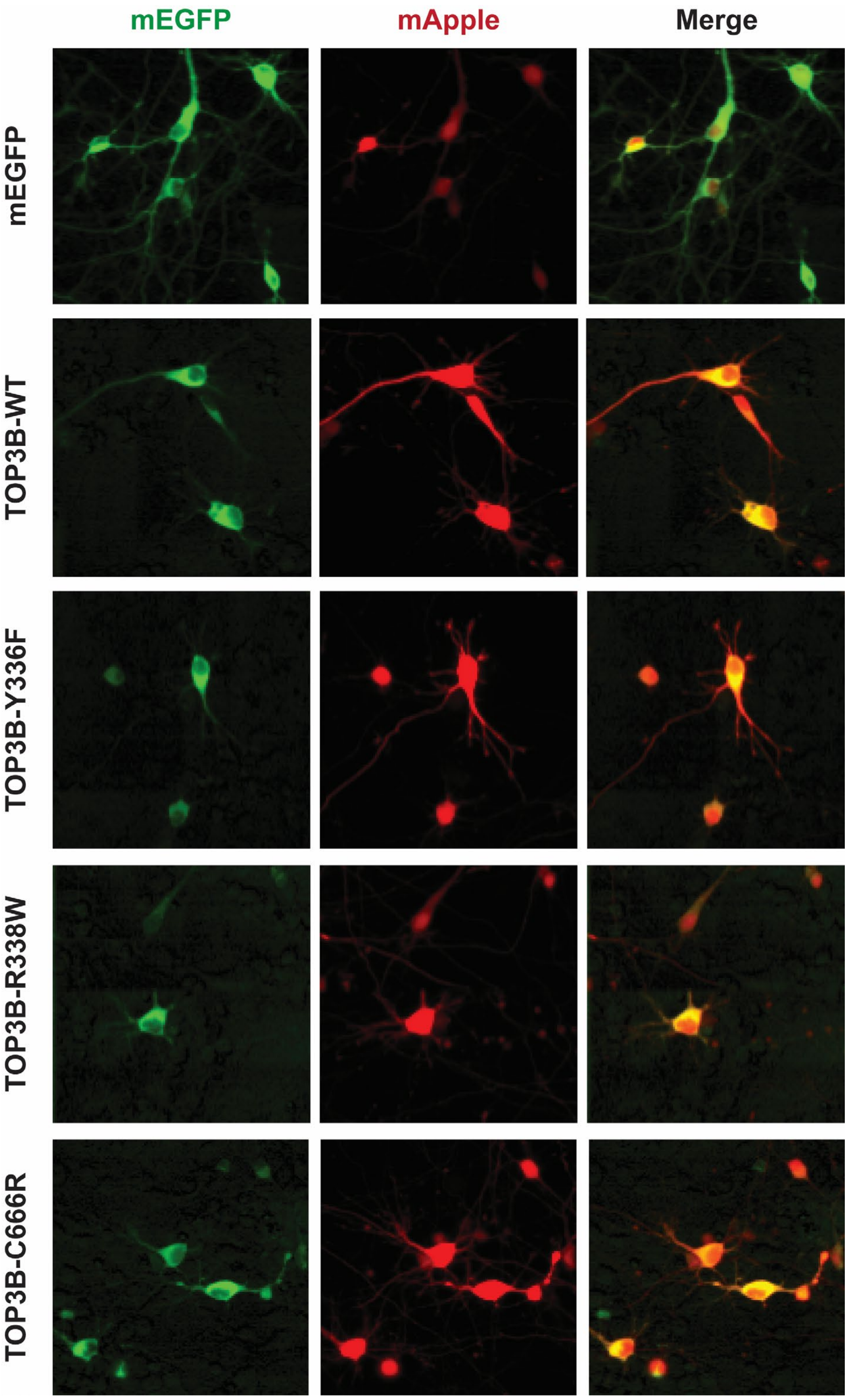


**Supplementary Figure S7. mEGFP-NES and mEGFP-NES-TOP3B fusion proteins are predominantly cytoplasmic in primary neurons.** Fluorescence microscopy of primary rat cortical neurons expressing mEGFP-NES and mEGFP-NES-TOP3B-WT, -Y336F, -R338W, and -C666R. mApple is co-transfected to mark the neuron body.

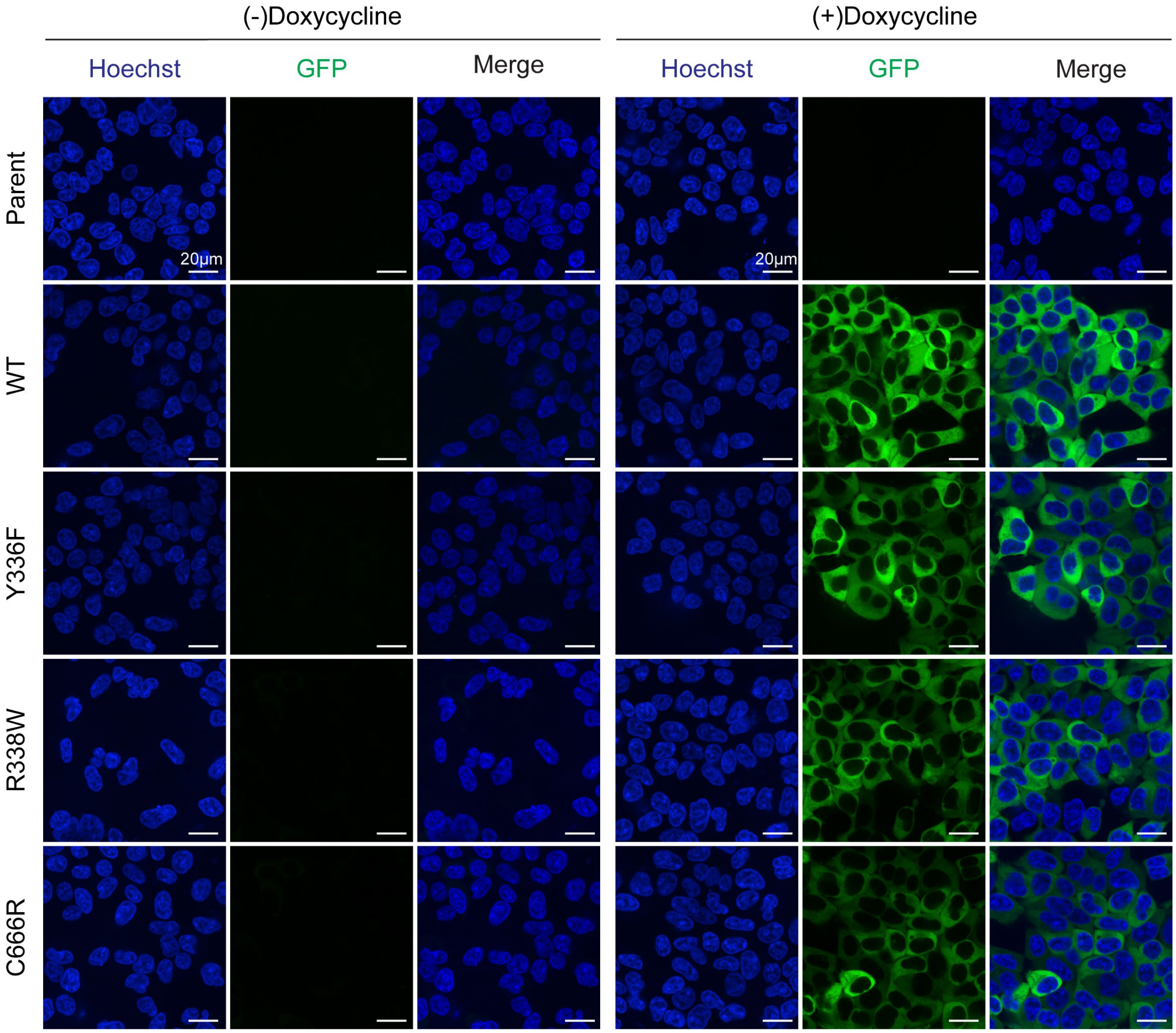


**Supplementary Figure S8. Doxycycline-inducible TOP3B is primarily cytoplasmic.** Spinning disk confocal microscopy of the parental Flp-In T-REx 293 cell line and stable cell lines with inducible expression of 3xFLAG-mEGFP-NES-TOP3B (WT, Y336F, and R338W, and C666R). Where indicated, expression was induced with doxycycline (Dox; 1 µg/mL final) for 48 hrs. DNA was stained using Hoechst 33342 (Hoechst; 1 µg/mL final). Images were taken at 60X magnification.

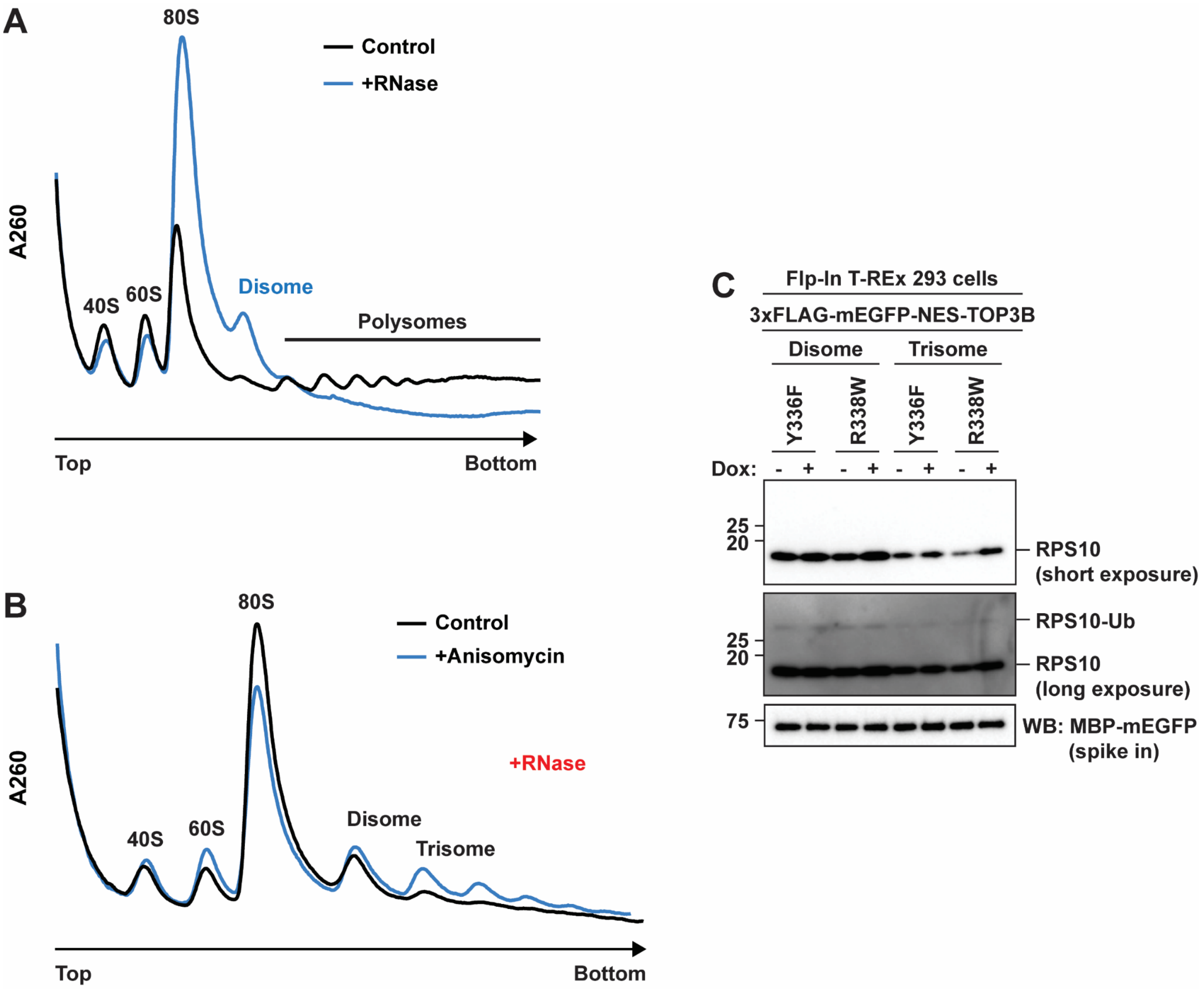


**Supplementary Figure S9. Micrococcal nuclease treatment effectively eliminates polysomes and allows for the detection of nuclease-resistant collied ribosomes.** A) Polysome analysis (10-50% (w/v) sucrose gradient) of uninduced 3xFLAG-mEGFP-NES-TOP3B (Y336F) Flp-In T-REx 293 cells. Lysates were either mock untreated (Control) or treated with S7 micrococcal nuclease (+RNase). B) Polysome analysis of uninduced 3xFLAG-mEGFP-NES-TOP3B (Y336F) Flp-In T-REx 293 cells treated with vehicle or intermediate dose of anisomycin (0.2 μg/mL final) for 15 min prior to harvesting. Both lysates were treated with S7 micrococcal nuclease for detection of nuclease-resistant ribosome collisions. C) Anti-RPS10 Western blot showing the expected ubiquitylation of RPS10 caused by collided ribosomes.

**OPEN READING FRAMES USED IN THIS MANUSCRIPT**

Full length TOP3B sequence
Point mutation
MYC tag
3xFLAG tag
mEGFP
V5 tag
Ubiquitin
NES
MBP
TEV
FLAG tag
Zinc finger only of TOP3B
Strep tag
His6 tag

### ORFs in plasmids for transfection of Neuro2A cells

**TOP3B-3xFLAG WT**
ATGAAGACTGTGCTCATGGTTGCTGAAAAGCCGTCCTTGGCACAGTCAATTGCCAAAATCCT
CTCTAGAGGGAGCCTGTCCTCACACAAAGGGCTGAACGGGGCCTGCTCAGTCCACGAGTA
CACTGGGACCTTTGCTGGCCAGCCAGTGCGCTTCAAGATGACGTCTGTCTGTGGTCACGT
GATGACCCTGGATTTCCTGGGAAAATACAACAAATGGGACAAAGTGGACCCCGCAGAACTG
TTCAGCCAAGCTCCCACGGAGAAGAAAGAAGCTAACCCCAAGCTGAACATGGTGAAGTTCC
TGCAGGTGGAGGGCAGAGGCTGCGACTACATCGTGCTGTGGCTGGACTGCGACAAGGAG
GGGGAGAACATCTGCTTTGAGGTTCTTGATGCTGTTCTGCCCGTCATGAACAAGGCCCATG
GTGGCGAGAAGACCGTGTTCCGGGCCAGGTTTAGCTCCATCACGGACACAGACATCTGTA
ATGCCATGGCCTGCCTAGGCGAGCCTGACCACAACGAGGCGCTCTCAGTGGATGCTCGCC
AGGAGCTGGACCTGCGAATCGGCTGTGCATTCACCAGGTTTCAGACTAAATATTTCCAGGG
GAAATACGGTGATTTAGACAGCTCTCTCATCTCCTTTGGGCCGTGTCAGACTCCAACCCTG
GGATTCTGTGTGGAGAGACATGATAAAATCCAGTCCTTCAAACCAGAGACCTACTGGGTGC
TGCAGGCCAAGGTTAACACTGACAAAGACAGATCTCTCCTTTTGGACTGGGACCGAGTAAG
AGTGTTTGACCGGGAGATCGCACAGATGTTTTTAAACATGACAAAGCTGGAGAAGGAAGCC
CAGGTGGAGGCCACAAGCAGGAAAGAAAAGGCCAAGCAGAGGCCCCTGGCCCTGAACAC
TGTGGAGATGCTGCGTGTGGCCAGCTCTTCTCTGGGCATGGGGCCGCAGCACGCCATGCA
GACGGCTGAGCGGCTCTACACGCAAGGCTACATCAGCTACCCACGGACAGAGACCACCCA
CTACCCTGAGAACTTTGACCTGAAGGGCTCTCTGCGGCAGCAGGCCAACCACCCCTACTG
GGCCGACACGGTGAAGCGGTTGTTAGCAGAAGGTATCAACCGCCCGCGGAAAGGCCATGA
CGCCGGCGACCATCCCCCCATCACCCCCATGAAGTCTGCCACAGAGGCCGAATTAGGGGG
TGACGCGTGGCGGCTCTATGAGTACATCACCAGACACTTCATCGCCACGGTCAGCCATGAC
TGCAAGTACCTGCAGAGCACCATCTCCTTCAGAATTGGGCCCGAGCTCTTCACCTGCTCCG
GGAAGACCGTCCTCTCACCAGGCTTCACGGAGGTCATGCCCTGGCAGAGCGTGCCCCTG
GAGGAGAGCCTGCCCACTTGCCAGCGGGGTGATGCCTTCCCTGTGGGCGAGGTGAAGAT
GCTGGAGAAGCAGACGAACCCACCCGACTACCTGACGGAGGCCGAGCTCATCACGCTCAT
GGAGAAGCATGGCATCGGCACGGATGCCAGCATCCCTGTGCATATCAACAACATCTGCCAG
CGCAACTATGTCACGGTGGAGAGCGGGCGCCGGCTCAAGCCCACCAACCTCGGCATCGT
CCTGGTGCACGGCTACTATAAGATTGATGCAGAGCTGGTGCTCCCCACCATCCGCAGTGCA
GTGGAGAAGCAGCTGAACCTGATCGCCCAGGGCAAGGCCGACTACCGCCAGGTCCTGGG

CCACACCCTGGACGTGTTCAAGAGGAAGTTCCACTACTTTGTCGACTCCATTGCTGGCATG
GATGAGTTGATGGAGGTGTCTTTCTCGCCCCTGGCGGCCACAGGCAAGCCCCTCTCACGC
TGTGGGAAGTGCCACCGCTTCATGAAGTACATCCAGGCCAAGCCAAGCCGCCTGCACTGC
TCCCACTGCGATGAGACCTACACGCTCCCCCAGAACGGCACCATCAAGCTCTACAAGGAG
CTCCGCTGCCCTCTGGATGACTTCGAGCTGGTCCTGTGGTCATCAGGCTCTCGGGGCAAG
AGCTACCCGCTGTGCCCCTACTGCTACAACCACCCACCCTTCCGAGACATGAAGAAAGGCA
TGGGCTGCAACGAGTGTACGCACCCCTCCTGCCAGCACTCGCTGAGCATGCTGGGCATCG
GCCAGTGCGTGGAATGTGAGAGCGGGGTGCTGGTGCTGGACCCCACCTCGGGCCCCAAG
TGGAAGGTGGCCTGCAACAAGTGCAACGTGGTAGCGCACTGCTTCGAGAACGCCCACCGC
GTGCGGGTGTCCGCCGACACCTGCAGTGTCTGTGAGGCCGCCTTGCTTGATGTGGACTTC
AACAAGGCCAAGTCCCCACTCCCGGGCGATGAGACGCAGCACATGGGCTGCGTCTTTTGT
GACCCCGTCTTCCAGGAGCTGGTGGAGCTGAAGCATGCGGCCTCCTGCCACCCCATGCAC
CGCGGTGGACCAGGGAGAAGGCAGGGTCGAGGGCGGGGCCGGGCCAGGAGGCCCCCT
GGGAAGCCCAACCCCAGACGGCCCAAGGACAAGATGTCAGCCCTGGCCGCCTACTTTGTA
AGCGGACCGACGCGTACGCGGCCGCTCGAGCAGAAACTCATCTCAGAAGAGGATCTGGCA
GCAAATGATATCCTGgactacaaagaccatgacggtgattataaagatcatgacatcgattacaaggatgacgatgacaag
GTTTAA

**TOP3B-3xFLAG Y336F**

ATGAAGACTGTGCTCATGGTTGCTGAAAAGCCGTCCTTGGCACAGTCAATTGCCAAAATCCT
CTCTAGAGGGAGCCTGTCCTCACACAAAGGGCTGAACGGGGCCTGCTCAGTCCACGAGTA
CACTGGGACCTTTGCTGGCCAGCCAGTGCGCTTCAAGATGACGTCTGTCTGTGGTCACGT
GATGACCCTGGATTTCCTGGGAAAATACAACAAATGGGACAAAGTGGACCCCGCAGAACTG
TTCAGCCAAGCTCCCACGGAGAAGAAAGAAGCTAACCCCAAGCTGAACATGGTGAAGTTCC
TGCAGGTGGAGGGCAGAGGCTGCGACTACATCGTGCTGTGGCTGGACTGCGACAAGGAG
GGGGAGAACATCTGCTTTGAGGTTCTTGATGCTGTTCTGCCCGTCATGAACAAGGCCCATG
GTGGCGAGAAGACCGTGTTCCGGGCCAGGTTTAGCTCCATCACGGACACAGACATCTGTA
ATGCCATGGCCTGCCTAGGCGAGCCTGACCACAACGAGGCGCTCTCAGTGGATGCTCGCC
AGGAGCTGGACCTGCGAATCGGCTGTGCATTCACCAGGTTTCAGACTAAATATTTCCAGGG
GAAATACGGTGATTTAGACAGCTCTCTCATCTCCTTTGGGCCGTGTCAGACTCCAACCCTG
GGATTCTGTGTGGAGAGACATGATAAAATCCAGTCCTTCAAACCAGAGACCTACTGGGTGC
TGCAGGCCAAGGTTAACACTGACAAAGACAGATCTCTCCTTTTGGACTGGGACCGAGTAAG
AGTGTTTGACCGGGAGATCGCACAGATGTTTTTAAACATGACAAAGCTGGAGAAGGAAGCC
CAGGTGGAGGCCACAAGCAGGAAAGAAAAGGCCAAGCAGAGGCCCCTGGCCCTGAACAC
TGTGGAGATGCTGCGTGTGGCCAGCTCTTCTCTGGGCATGGGGCCGCAGCACGCCATGCA
GACGGCTGAGCGGCTCTACACGCAAGGCTACATCAGCtttCCACGGACAGAGACCACCCACT
ACCCTGAGAACTTTGACCTGAAGGGCTCTCTGCGGCAGCAGGCCAACCACCCCTACTGGG
CCGACACGGTGAAGCGGTTGTTAGCAGAAGGTATCAACCGCCCGCGGAAAGGCCATGACG
CCGGCGACCATCCCCCCATCACCCCCATGAAGTCTGCCACAGAGGCCGAATTAGGGGGTG
ACGCGTGGCGGCTCTATGAGTACATCACCAGACACTTCATCGCCACGGTCAGCCATGACTG
CAAGTACCTGCAGAGCACCATCTCCTTCAGAATTGGGCCCGAGCTCTTCACCTGCTCCGGG
AAGACCGTCCTCTCACCAGGCTTCACGGAGGTCATGCCCTGGCAGAGCGTGCCCCTGGAG
GAGAGCCTGCCCACTTGCCAGCGGGGTGATGCCTTCCCTGTGGGCGAGGTGAAGATGCT
GGAGAAGCAGACGAACCCACCCGACTACCTGACGGAGGCCGAGCTCATCACGCTCATGGA
GAAGCATGGCATCGGCACGGATGCCAGCATCCCTGTGCATATCAACAACATCTGCCAGCGC
AACTATGTCACGGTGGAGAGCGGGCGCCGGCTCAAGCCCACCAACCTCGGCATCGTCCTG
GTGCACGGCTACTATAAGATTGATGCAGAGCTGGTGCTCCCCACCATCCGCAGTGCAGTGG
AGAAGCAGCTGAACCTGATCGCCCAGGGCAAGGCCGACTACCGCCAGGTCCTGGGCCAC
ACCCTGGACGTGTTCAAGAGGAAGTTCCACTACTTTGTCGACTCCATTGCTGGCATGGATG
AGTTGATGGAGGTGTCTTTCTCGCCCCTGGCGGCCACAGGCAAGCCCCTCTCACGCTGTG
GGAAGTGCCACCGCTTCATGAAGTACATCCAGGCCAAGCCAAGCCGCCTGCACTGCTCCC

ACTGCGATGAGACCTACACGCTCCCCCAGAACGGCACCATCAAGCTCTACAAGGAGCTCC
GCTGCCCTCTGGATGACTTCGAGCTGGTCCTGTGGTCATCAGGCTCTCGGGGCAAGAGCT
ACCCGCTGTGCCCCTACTGCTACAACCACCCACCCTTCCGAGACATGAAGAAAGGCATGG
GCTGCAACGAGTGTACGCACCCCTCCTGCCAGCACTCGCTGAGCATGCTGGGCATCGGCC
AGTGCGTGGAATGTGAGAGCGGGGTGCTGGTGCTGGACCCCACCTCGGGCCCCAAGTGG
AAGGTGGCCTGCAACAAGTGCAACGTGGTAGCGCACTGCTTCGAGAACGCCCACCGCGTG
CGGGTGTCCGCCGACACCTGCAGTGTCTGTGAGGCCGCCTTGCTTGATGTGGACTTCAAC
AAGGCCAAGTCCCCACTCCCGGGCGATGAGACGCAGCACATGGGCTGCGTCTTTTGTGAC
CCCGTCTTCCAGGAGCTGGTGGAGCTGAAGCATGCGGCCTCCTGCCACCCCATGCACCGC
GGTGGACCAGGGAGAAGGCAGGGTCGAGGGCGGGGCCGGGCCAGGAGGCCCCCTGGG
AAGCCCAACCCCAGACGGCCCAAGGACAAGATGTCAGCCCTGGCCGCCTACTTTGTAAGC
GGACCGACGCGTACGCGGCCGCTCGAGCAGAAACTCATCTCAGAAGAGGATCTGGCAGCA
AATGATATCCTGgactacaaagaccatgacggtgattataaagatcatgacatcgattacaaggatgacgatgacaagGTT
TAA

**TOP3B-3xFLAG R338W**

ATGAAGACTGTGCTCATGGTTGCTGAAAAGCCGTCCTTGGCACAGTCAATTGCCAAAATCCT
CTCTAGAGGGAGCCTGTCCTCACACAAAGGGCTGAACGGGGCCTGCTCAGTCCACGAGTA
CACTGGGACCTTTGCTGGCCAGCCAGTGCGCTTCAAGATGACGTCTGTCTGTGGTCACGT
GATGACCCTGGATTTCCTGGGAAAATACAACAAATGGGACAAAGTGGACCCCGCAGAACTG
TTCAGCCAAGCTCCCACGGAGAAGAAAGAAGCTAACCCCAAGCTGAACATGGTGAAGTTCC
TGCAGGTGGAGGGCAGAGGCTGCGACTACATCGTGCTGTGGCTGGACTGCGACAAGGAG
GGGGAGAACATCTGCTTTGAGGTTCTTGATGCTGTTCTGCCCGTCATGAACAAGGCCCATG
GTGGCGAGAAGACCGTGTTCCGGGCCAGGTTTAGCTCCATCACGGACACAGACATCTGTA
ATGCCATGGCCTGCCTAGGCGAGCCTGACCACAACGAGGCGCTCTCAGTGGATGCTCGCC
AGGAGCTGGACCTGCGAATCGGCTGTGCATTCACCAGGTTTCAGACTAAATATTTCCAGGG
GAAATACGGTGATTTAGACAGCTCTCTCATCTCCTTTGGGCCGTGTCAGACTCCAACCCTG
GGATTCTGTGTGGAGAGACATGATAAAATCCAGTCCTTCAAACCAGAGACCTACTGGGTGC
TGCAGGCCAAGGTTAACACTGACAAAGACAGATCTCTCCTTTTGGACTGGGACCGAGTAAG
AGTGTTTGACCGGGAGATCGCACAGATGTTTTTAAACATGACAAAGCTGGAGAAGGAAGCC
CAGGTGGAGGCCACAAGCAGGAAAGAAAAGGCCAAGCAGAGGCCCCTGGCCCTGAACAC
TGTGGAGATGCTGCGTGTGGCCAGCTCTTCTCTGGGCATGGGGCCGCAGCACGCCATGCA
GACGGCTGAGCGGCTCTACACGCAAGGCTACATCAGCTACCCAtggACAGAGACCACCCAC
TACCCTGAGAACTTTGACCTGAAGGGCTCTCTGCGGCAGCAGGCCAACCACCCCTACTGG
GCCGACACGGTGAAGCGGTTGTTAGCAGAAGGTATCAACCGCCCGCGGAAAGGCCATGAC
GCCGGCGACCATCCCCCCATCACCCCCATGAAGTCTGCCACAGAGGCCGAATTAGGGGGT
GACGCGTGGCGGCTCTATGAGTACATCACCAGACACTTCATCGCCACGGTCAGCCATGACT
GCAAGTACCTGCAGAGCACCATCTCCTTCAGAATTGGGCCCGAGCTCTTCACCTGCTCCGG
GAAGACCGTCCTCTCACCAGGCTTCACGGAGGTCATGCCCTGGCAGAGCGTGCCCCTGGA
GGAGAGCCTGCCCACTTGCCAGCGGGGTGATGCCTTCCCTGTGGGCGAGGTGAAGATGC
TGGAGAAGCAGACGAACCCACCCGACTACCTGACGGAGGCCGAGCTCATCACGCTCATGG
AGAAGCATGGCATCGGCACGGATGCCAGCATCCCTGTGCATATCAACAACATCTGCCAGCG
CAACTATGTCACGGTGGAGAGCGGGCGCCGGCTCAAGCCCACCAACCTCGGCATCGTCCT
GGTGCACGGCTACTATAAGATTGATGCAGAGCTGGTGCTCCCCACCATCCGCAGTGCAGTG
GAGAAGCAGCTGAACCTGATCGCCCAGGGCAAGGCCGACTACCGCCAGGTCCTGGGCCA
CACCCTGGACGTGTTCAAGAGGAAGTTCCACTACTTTGTCGACTCCATTGCTGGCATGGAT
GAGTTGATGGAGGTGTCTTTCTCGCCCCTGGCGGCCACAGGCAAGCCCCTCTCACGCTGT
GGGAAGTGCCACCGCTTCATGAAGTACATCCAGGCCAAGCCAAGCCGCCTGCACTGCTCC
CACTGCGATGAGACCTACACGCTCCCCCAGAACGGCACCATCAAGCTCTACAAGGAGCTC
CGCTGCCCTCTGGATGACTTCGAGCTGGTCCTGTGGTCATCAGGCTCTCGGGGCAAGAGC
TACCCGCTGTGCCCCTACTGCTACAACCACCCACCCTTCCGAGACATGAAGAAAGGCATGG

GCTGCAACGAGTGTACGCACCCCTCCTGCCAGCACTCGCTGAGCATGCTGGGCATCGGCC
AGTGCGTGGAATGTGAGAGCGGGGTGCTGGTGCTGGACCCCACCTCGGGCCCCAAGTGG
AAGGTGGCCTGCAACAAGTGCAACGTGGTAGCGCACTGCTTCGAGAACGCCCACCGCGTG
CGGGTGTCCGCCGACACCTGCAGTGTCTGTGAGGCCGCCTTGCTTGATGTGGACTTCAAC
AAGGCCAAGTCCCCACTCCCGGGCGATGAGACGCAGCACATGGGCTGCGTCTTTTGTGAC
CCCGTCTTCCAGGAGCTGGTGGAGCTGAAGCATGCGGCCTCCTGCCACCCCATGCACCGC
GGTGGACCAGGGAGAAGGCAGGGTCGAGGGCGGGGCCGGGCCAGGAGGCCCCCTGGG
AAGCCCAACCCCAGACGGCCCAAGGACAAGATGTCAGCCCTGGCCGCCTACTTTGTAAGC
GGACCGACGCGTACGCGGCCGCTCGAGCAGAAACTCATCTCAGAAGAGGATCTGGCAGCA
AATGATATCCTGgactacaaagaccatgacggtgattataaagatcatgacatcgattacaaggatgacgatgacaagGTT
TAA

**TOP3B-3xFLAG P378Q**

ATGAAGACTGTGCTCATGGTTGCTGAAAAGCCGTCCTTGGCACAGTCAATTGCCAAAATCCT
CTCTAGAGGGAGCCTGTCCTCACACAAAGGGCTGAACGGGGCCTGCTCAGTCCACGAGTA
CACTGGGACCTTTGCTGGCCAGCCAGTGCGCTTCAAGATGACGTCTGTCTGTGGTCACGT
GATGACCCTGGATTTCCTGGGAAAATACAACAAATGGGACAAAGTGGACCCCGCAGAACTG
TTCAGCCAAGCTCCCACGGAGAAGAAAGAAGCTAACCCCAAGCTGAACATGGTGAAGTTCC
TGCAGGTGGAGGGCAGAGGCTGCGACTACATCGTGCTGTGGCTGGACTGCGACAAGGAG
GGGGAGAACATCTGCTTTGAGGTTCTTGATGCTGTTCTGCCCGTCATGAACAAGGCCCATG
GTGGCGAGAAGACCGTGTTCCGGGCCAGGTTTAGCTCCATCACGGACACAGACATCTGTA
ATGCCATGGCCTGCCTAGGCGAGCCTGACCACAACGAGGCGCTCTCAGTGGATGCTCGCC
AGGAGCTGGACCTGCGAATCGGCTGTGCATTCACCAGGTTTCAGACTAAATATTTCCAGGG
GAAATACGGTGATTTAGACAGCTCTCTCATCTCCTTTGGGCCGTGTCAGACTCCAACCCTG
GGATTCTGTGTGGAGAGACATGATAAAATCCAGTCCTTCAAACCAGAGACCTACTGGGTGC
TGCAGGCCAAGGTTAACACTGACAAAGACAGATCTCTCCTTTTGGACTGGGACCGAGTAAG
AGTGTTTGACCGGGAGATCGCACAGATGTTTTTAAACATGACAAAGCTGGAGAAGGAAGCC
CAGGTGGAGGCCACAAGCAGGAAAGAAAAGGCCAAGCAGAGGCCCCTGGCCCTGAACAC
TGTGGAGATGCTGCGTGTGGCCAGCTCTTCTCTGGGCATGGGGCCGCAGCACGCCATGCA
GACGGCTGAGCGGCTCTACACGCAAGGCTACATCAGCTACCCACGGACAGAGACCACCCA
CTACCCTGAGAACTTTGACCTGAAGGGCTCTCTGCGGCAGCAGGCCAACCACCCCTACTG
GGCCGACACGGTGAAGCGGTTGTTAGCAGAAGGTATCAACCGCcagCGGAAAGGCCATGAC
GCCGGCGACCATCCCCCCATCACCCCCATGAAGTCTGCCACAGAGGCCGAATTAGGGGGT
GACGCGTGGCGGCTCTATGAGTACATCACCAGACACTTCATCGCCACGGTCAGCCATGACT
GCAAGTACCTGCAGAGCACCATCTCCTTCAGAATTGGGCCCGAGCTCTTCACCTGCTCCGG
GAAGACCGTCCTCTCACCAGGCTTCACGGAGGTCATGCCCTGGCAGAGCGTGCCCCTGGA
GGAGAGCCTGCCCACTTGCCAGCGGGGTGATGCCTTCCCTGTGGGCGAGGTGAAGATGC
TGGAGAAGCAGACGAACCCACCCGACTACCTGACGGAGGCCGAGCTCATCACGCTCATGG
AGAAGCATGGCATCGGCACGGATGCCAGCATCCCTGTGCATATCAACAACATCTGCCAGCG
CAACTATGTCACGGTGGAGAGCGGGCGCCGGCTCAAGCCCACCAACCTCGGCATCGTCCT
GGTGCACGGCTACTATAAGATTGATGCAGAGCTGGTGCTCCCCACCATCCGCAGTGCAGTG
GAGAAGCAGCTGAACCTGATCGCCCAGGGCAAGGCCGACTACCGCCAGGTCCTGGGCCA
CACCCTGGACGTGTTCAAGAGGAAGTTCCACTACTTTGTCGACTCCATTGCTGGCATGGAT
GAGTTGATGGAGGTGTCTTTCTCGCCCCTGGCGGCCACAGGCAAGCCCCTCTCACGCTGT
GGGAAGTGCCACCGCTTCATGAAGTACATCCAGGCCAAGCCAAGCCGCCTGCACTGCTCC
CACTGCGATGAGACCTACACGCTCCCCCAGAACGGCACCATCAAGCTCTACAAGGAGCTC
CGCTGCCCTCTGGATGACTTCGAGCTGGTCCTGTGGTCATCAGGCTCTCGGGGCAAGAGC
TACCCGCTGTGCCCCTACTGCTACAACCACCCACCCTTCCGAGACATGAAGAAAGGCATGG
GCTGCAACGAGTGTACGCACCCCTCCTGCCAGCACTCGCTGAGCATGCTGGGCATCGGCC
AGTGCGTGGAATGTGAGAGCGGGGTGCTGGTGCTGGACCCCACCTCGGGCCCCAAGTGG
AAGGTGGCCTGCAACAAGTGCAACGTGGTAGCGCACTGCTTCGAGAACGCCCACCGCGTG

CGGGTGTCCGCCGACACCTGCAGTGTCTGTGAGGCCGCCTTGCTTGATGTGGACTTCAAC
AAGGCCAAGTCCCCACTCCCGGGCGATGAGACGCAGCACATGGGCTGCGTCTTTTGTGAC
CCCGTCTTCCAGGAGCTGGTGGAGCTGAAGCATGCGGCCTCCTGCCACCCCATGCACCGC
GGTGGACCAGGGAGAAGGCAGGGTCGAGGGCGGGGCCGGGCCAGGAGGCCCCCTGGG
AAGCCCAACCCCAGACGGCCCAAGGACAAGATGTCAGCCCTGGCCGCCTACTTTGTAAGC
GGACCGACGCGTACGCGGCCGCTCGAGCAGAAACTCATCTCAGAAGAGGATCTGGCAGCA
AATGATATCCTGgactacaaagaccatgacggtgattataaagatcatgacatcgattacaaggatgacgatgacaagGTT
TAA

**TOP3B-3xFLAG R472Q**

ATGAAGACTGTGCTCATGGTTGCTGAAAAGCCGTCCTTGGCACAGTCAATTGCCAAAATCCT
CTCTAGAGGGAGCCTGTCCTCACACAAAGGGCTGAACGGGGCCTGCTCAGTCCACGAGTA
CACTGGGACCTTTGCTGGCCAGCCAGTGCGCTTCAAGATGACGTCTGTCTGTGGTCACGT
GATGACCCTGGATTTCCTGGGAAAATACAACAAATGGGACAAAGTGGACCCCGCAGAACTG
TTCAGCCAAGCTCCCACGGAGAAGAAAGAAGCTAACCCCAAGCTGAACATGGTGAAGTTCC
TGCAGGTGGAGGGCAGAGGCTGCGACTACATCGTGCTGTGGCTGGACTGCGACAAGGAG
GGGGAGAACATCTGCTTTGAGGTTCTTGATGCTGTTCTGCCCGTCATGAACAAGGCCCATG
GTGGCGAGAAGACCGTGTTCCGGGCCAGGTTTAGCTCCATCACGGACACAGACATCTGTA
ATGCCATGGCCTGCCTAGGCGAGCCTGACCACAACGAGGCGCTCTCAGTGGATGCTCGCC
AGGAGCTGGACCTGCGAATCGGCTGTGCATTCACCAGGTTTCAGACTAAATATTTCCAGGG
GAAATACGGTGATTTAGACAGCTCTCTCATCTCCTTTGGGCCGTGTCAGACTCCAACCCTG
GGATTCTGTGTGGAGAGACATGATAAAATCCAGTCCTTCAAACCAGAGACCTACTGGGTGC
TGCAGGCCAAGGTTAACACTGACAAAGACAGATCTCTCCTTTTGGACTGGGACCGAGTAAG
AGTGTTTGACCGGGAGATCGCACAGATGTTTTTAAACATGACAAAGCTGGAGAAGGAAGCC
CAGGTGGAGGCCACAAGCAGGAAAGAAAAGGCCAAGCAGAGGCCCCTGGCCCTGAACAC
TGTGGAGATGCTGCGTGTGGCCAGCTCTTCTCTGGGCATGGGGCCGCAGCACGCCATGCA
GACGGCTGAGCGGCTCTACACGCAAGGCTACATCAGCTACCCACGGACAGAGACCACCCA
CTACCCTGAGAACTTTGACCTGAAGGGCTCTCTGCGGCAGCAGGCCAACCACCCCTACTG
GGCCGACACGGTGAAGCGGTTGTTAGCAGAAGGTATCAACCGCCCGCGGAAAGGCCATGA
CGCCGGCGACCATCCCCCCATCACCCCCATGAAGTCTGCCACAGAGGCCGAATTAGGGGG
TGACGCGTGGCGGCTCTATGAGTACATCACCAGACACTTCATCGCCACGGTCAGCCATGAC
TGCAAGTACCTGCAGAGCACCATCTCCTTCAGAATTGGGCCCGAGCTCTTCACCTGCTCCG
GGAAGACCGTCCTCTCACCAGGCTTCACGGAGGTCATGCCCTGGCAGAGCGTGCCCCTG
GAGGAGAGCCTGCCCACTTGCCAGcagGGTGATGCCTTCCCTGTGGGCGAGGTGAAGATG
CTGGAGAAGCAGACGAACCCACCCGACTACCTGACGGAGGCCGAGCTCATCACGCTCATG
GAGAAGCATGGCATCGGCACGGATGCCAGCATCCCTGTGCATATCAACAACATCTGCCAGC
GCAACTATGTCACGGTGGAGAGCGGGCGCCGGCTCAAGCCCACCAACCTCGGCATCGTC
CTGGTGCACGGCTACTATAAGATTGATGCAGAGCTGGTGCTCCCCACCATCCGCAGTGCAG
TGGAGAAGCAGCTGAACCTGATCGCCCAGGGCAAGGCCGACTACCGCCAGGTCCTGGGC
CACACCCTGGACGTGTTCAAGAGGAAGTTCCACTACTTTGTCGACTCCATTGCTGGCATGG
ATGAGTTGATGGAGGTGTCTTTCTCGCCCCTGGCGGCCACAGGCAAGCCCCTCTCACGCT
GTGGGAAGTGCCACCGCTTCATGAAGTACATCCAGGCCAAGCCAAGCCGCCTGCACTGCT
CCCACTGCGATGAGACCTACACGCTCCCCCAGAACGGCACCATCAAGCTCTACAAGGAGC
TCCGCTGCCCTCTGGATGACTTCGAGCTGGTCCTGTGGTCATCAGGCTCTCGGGGCAAGA
GCTACCCGCTGTGCCCCTACTGCTACAACCACCCACCCTTCCGAGACATGAAGAAAGGCAT
GGGCTGCAACGAGTGTACGCACCCCTCCTGCCAGCACTCGCTGAGCATGCTGGGCATCGG
CCAGTGCGTGGAATGTGAGAGCGGGGTGCTGGTGCTGGACCCCACCTCGGGCCCCAAGT
GGAAGGTGGCCTGCAACAAGTGCAACGTGGTAGCGCACTGCTTCGAGAACGCCCACCGC
GTGCGGGTGTCCGCCGACACCTGCAGTGTCTGTGAGGCCGCCTTGCTTGATGTGGACTTC
AACAAGGCCAAGTCCCCACTCCCGGGCGATGAGACGCAGCACATGGGCTGCGTCTTTTGT
GACCCCGTCTTCCAGGAGCTGGTGGAGCTGAAGCATGCGGCCTCCTGCCACCCCATGCAC

CGCGGTGGACCAGGGAGAAGGCAGGGTCGAGGGCGGGGCCGGGCCAGGAGGCCCCCT
GGGAAGCCCAACCCCAGACGGCCCAAGGACAAGATGTCAGCCCTGGCCGCCTACTTTGTA
AGCGGACCGACGCGTACGCGGCCGCTCGAGCAGAAACTCATCTCAGAAGAGGATCTGGCA
GCAAATGATATCCTGgactacaaagaccatgacggtgattataaagatcatgacatcgattacaaggatgacgatgacaag
GTTTAA

**TOP3B-3xFLAG C666R**

ATGAAGACTGTGCTCATGGTTGCTGAAAAGCCGTCCTTGGCACAGTCAATTGCCAAAATCCT
CTCTAGAGGGAGCCTGTCCTCACACAAAGGGCTGAACGGGGCCTGCTCAGTCCACGAGTA
CACTGGGACCTTTGCTGGCCAGCCAGTGCGCTTCAAGATGACGTCTGTCTGTGGTCACGT
GATGACCCTGGATTTCCTGGGAAAATACAACAAATGGGACAAAGTGGACCCCGCAGAACTG
TTCAGCCAAGCTCCCACGGAGAAGAAAGAAGCTAACCCCAAGCTGAACATGGTGAAGTTCC
TGCAGGTGGAGGGCAGAGGCTGCGACTACATCGTGCTGTGGCTGGACTGCGACAAGGAG
GGGGAGAACATCTGCTTTGAGGTTCTTGATGCTGTTCTGCCCGTCATGAACAAGGCCCATG
GTGGCGAGAAGACCGTGTTCCGGGCCAGGTTTAGCTCCATCACGGACACAGACATCTGTA
ATGCCATGGCCTGCCTAGGCGAGCCTGACCACAACGAGGCGCTCTCAGTGGATGCTCGCC
AGGAGCTGGACCTGCGAATCGGCTGTGCATTCACCAGGTTTCAGACTAAATATTTCCAGGG
GAAATACGGTGATTTAGACAGCTCTCTCATCTCCTTTGGGCCGTGTCAGACTCCAACCCTG
GGATTCTGTGTGGAGAGACATGATAAAATCCAGTCCTTCAAACCAGAGACCTACTGGGTGC
TGCAGGCCAAGGTTAACACTGACAAAGACAGATCTCTCCTTTTGGACTGGGACCGAGTAAG
AGTGTTTGACCGGGAGATCGCACAGATGTTTTTAAACATGACAAAGCTGGAGAAGGAAGCC
CAGGTGGAGGCCACAAGCAGGAAAGAAAAGGCCAAGCAGAGGCCCCTGGCCCTGAACAC
TGTGGAGATGCTGCGTGTGGCCAGCTCTTCTCTGGGCATGGGGCCGCAGCACGCCATGCA
GACGGCTGAGCGGCTCTACACGCAAGGCTACATCAGCTACCCACGGACAGAGACCACCCA
CTACCCTGAGAACTTTGACCTGAAGGGCTCTCTGCGGCAGCAGGCCAACCACCCCTACTG
GGCCGACACGGTGAAGCGGTTGTTAGCAGAAGGTATCAACCGCCCGCGGAAAGGCCATGA
CGCCGGCGACCATCCCCCCATCACCCCCATGAAGTCTGCCACAGAGGCCGAATTAGGGGG
TGACGCGTGGCGGCTCTATGAGTACATCACCAGACACTTCATCGCCACGGTCAGCCATGAC
TGCAAGTACCTGCAGAGCACCATCTCCTTCAGAATTGGGCCCGAGCTCTTCACCTGCTCCG
GGAAGACCGTCCTCTCACCAGGCTTCACGGAGGTCATGCCCTGGCAGAGCGTGCCCCTG
GAGGAGAGCCTGCCCACTTGCCAGCGGGGTGATGCCTTCCCTGTGGGCGAGGTGAAGAT
GCTGGAGAAGCAGACGAACCCACCCGACTACCTGACGGAGGCCGAGCTCATCACGCTCAT
GGAGAAGCATGGCATCGGCACGGATGCCAGCATCCCTGTGCATATCAACAACATCTGCCAG
CGCAACTATGTCACGGTGGAGAGCGGGCGCCGGCTCAAGCCCACCAACCTCGGCATCGT
CCTGGTGCACGGCTACTATAAGATTGATGCAGAGCTGGTGCTCCCCACCATCCGCAGTGCA
GTGGAGAAGCAGCTGAACCTGATCGCCCAGGGCAAGGCCGACTACCGCCAGGTCCTGGG
CCACACCCTGGACGTGTTCAAGAGGAAGTTCCACTACTTTGTCGACTCCATTGCTGGCATG
GATGAGTTGATGGAGGTGTCTTTCTCGCCCCTGGCGGCCACAGGCAAGCCCCTCTCACGC
TGTGGGAAGTGCCACCGCTTCATGAAGTACATCCAGGCCAAGCCAAGCCGCCTGCACTGC
TCCCACTGCGATGAGACCTACACGCTCCCCCAGAACGGCACCATCAAGCTCTACAAGGAG
CTCCGCcgcCCTCTGGATGACTTCGAGCTGGTCCTGTGGTCATCAGGCTCTCGGGGCAAGA
GCTACCCGCTGTGCCCCTACTGCTACAACCACCCACCCTTCCGAGACATGAAGAAAGGCAT
GGGCTGCAACGAGTGTACGCACCCCTCCTGCCAGCACTCGCTGAGCATGCTGGGCATCGG
CCAGTGCGTGGAATGTGAGAGCGGGGTGCTGGTGCTGGACCCCACCTCGGGCCCCAAGT
GGAAGGTGGCCTGCAACAAGTGCAACGTGGTAGCGCACTGCTTCGAGAACGCCCACCGC
GTGCGGGTGTCCGCCGACACCTGCAGTGTCTGTGAGGCCGCCTTGCTTGATGTGGACTTC
AACAAGGCCAAGTCCCCACTCCCGGGCGATGAGACGCAGCACATGGGCTGCGTCTTTTGT
GACCCCGTCTTCCAGGAGCTGGTGGAGCTGAAGCATGCGGCCTCCTGCCACCCCATGCAC
CGCGGTGGACCAGGGAGAAGGCAGGGTCGAGGGCGGGGCCGGGCCAGGAGGCCCCCT
GGGAAGCCCAACCCCAGACGGCCCAAGGACAAGATGTCAGCCCTGGCCGCCTACTTTGTA
AGCGGACCGACGCGTACGCGGCCGCTCGAGCAGAAACTCATCTCAGAAGAGGATCTGGCA

GCAAATGATATCCTGgactacaaagaccatgacggtgattataaagatcatgacatcgattacaaggatgacgatgacaag
GTTTAA

**3xFLAG-TOP3B WT**

ATGggttcttctgactacaaagaccatgacggtgattataaagatcatgacatcgattacaaggatgacgatgacaagggttcttctA
AGACTGTGCTCATGGTTGCTGAAAAGCCGTCCTTGGCACAGTCAATTGCCAAAATCCTCTC
TAGAGGGAGCCTGTCCTCACACAAAGGGCTGAACGGGGCCTGCTCAGTCCACGAGTACAC
TGGGACCTTTGCTGGCCAGCCAGTGCGCTTCAAGATGACGTCTGTCTGTGGTCACGTGAT
GACCCTGGATTTCCTGGGAAAATACAACAAATGGGACAAAGTGGACCCCGCAGAACTGTTC
AGCCAAGCTCCCACGGAGAAGAAAGAAGCTAACCCCAAGCTGAACATGGTGAAGTTCCTG
CAGGTGGAGGGCAGAGGCTGCGACTACATCGTGCTGTGGCTGGACTGCGACAAGGAGGG
GGAGAACATCTGCTTTGAGGTTCTTGATGCTGTTCTGCCCGTCATGAACAAGGCCCATGGT
GGCGAGAAGACCGTGTTCCGGGCCAGGTTTAGCTCCATCACGGACACAGACATCTGTAATG
CCATGGCCTGCCTAGGCGAGCCTGACCACAACGAGGCGCTCTCAGTGGATGCTCGCCAG
GAGCTGGACCTGCGAATCGGCTGTGCATTCACCAGGTTTCAGACTAAATATTTCCAGGGGA
AATACGGTGATTTAGACAGCTCTCTCATCTCCTTTGGGCCGTGTCAGACTCCAACCCTGGG
ATTCTGTGTGGAGAGACATGATAAAATCCAGTCCTTCAAACCAGAGACCTACTGGGTGCTGC
AGGCCAAGGTTAACACTGACAAAGACAGATCTCTCCTTTTGGACTGGGACCGAGTAAGAGT
GTTTGACCGGGAGATCGCACAGATGTTTTTAAACATGACAAAGCTGGAGAAGGAAGCCCAG
GTGGAGGCCACAAGCAGGAAAGAAAAGGCCAAGCAGAGGCCCCTGGCCCTGAACACTGT
GGAGATGCTGCGTGTGGCCAGCTCTTCTCTGGGCATGGGGCCGCAGCACGCCATGCAGA
CGGCTGAGCGGCTCTACACGCAAGGCTACATCAGCTACCCACGGACAGAGACCACCCACT
ACCCTGAGAACTTTGACCTGAAGGGCTCTCTGCGGCAGCAGGCCAACCACCCCTACTGGG
CCGACACGGTGAAGCGGTTGTTAGCAGAAGGTATCAACCGCCCGCGGAAAGGCCATGACG
CCGGCGACCATCCCCCCATCACCCCCATGAAGTCTGCCACAGAGGCCGAATTAGGGGGTG
ACGCGTGGCGGCTCTATGAGTACATCACCAGACACTTCATCGCCACGGTCAGCCATGACTG
CAAGTACCTGCAGAGCACCATCTCCTTCAGAATTGGGCCCGAGCTCTTCACCTGCTCCGGG
AAGACCGTCCTCTCACCAGGCTTCACGGAGGTCATGCCCTGGCAGAGCGTGCCCCTGGAG
GAGAGCCTGCCCACTTGCCAGCGGGGTGATGCCTTCCCTGTGGGCGAGGTGAAGATGCT
GGAGAAGCAGACGAACCCACCCGACTACCTGACGGAGGCCGAGCTCATCACGCTCATGGA
GAAGCATGGCATCGGCACGGATGCCAGCATCCCTGTGCATATCAACAACATCTGCCAGCGC
AACTATGTCACGGTGGAGAGCGGGCGCCGGCTCAAGCCCACCAACCTCGGCATCGTCCTG
GTGCACGGCTACTATAAGATTGATGCAGAGCTGGTGCTCCCCACCATCCGCAGTGCAGTGG
AGAAGCAGCTGAACCTGATCGCCCAGGGCAAGGCCGACTACCGCCAGGTCCTGGGCCAC
ACCCTGGACGTGTTCAAGAGGAAGTTCCACTACTTTGTCGACTCCATTGCTGGCATGGATG
AGTTGATGGAGGTGTCTTTCTCGCCCCTGGCGGCCACAGGCAAGCCCCTCTCACGCTGTG
GGAAGTGCCACCGCTTCATGAAGTACATCCAGGCCAAGCCAAGCCGCCTGCACTGCTCCC
ACTGCGATGAGACCTACACGCTCCCCCAGAACGGCACCATCAAGCTCTACAAGGAGCTCC
GCTGCCCTCTGGATGACTTCGAGCTGGTCCTGTGGTCATCAGGCTCTCGGGGCAAGAGCT
ACCCGCTGTGCCCCTACTGCTACAACCACCCACCCTTCCGAGACATGAAGAAAGGCATGG
GCTGCAACGAGTGTACGCACCCCTCCTGCCAGCACTCGCTGAGCATGCTGGGCATCGGCC
AGTGCGTGGAATGTGAGAGCGGGGTGCTGGTGCTGGACCCCACCTCGGGCCCCAAGTGG
AAGGTGGCCTGCAACAAGTGCAACGTGGTAGCGCACTGCTTCGAGAACGCCCACCGCGTG
CGGGTGTCCGCCGACACCTGCAGTGTCTGTGAGGCCGCCTTGCTTGATGTGGACTTCAAC
AAGGCCAAGTCCCCACTCCCGGGCGATGAGACGCAGCACATGGGCTGCGTCTTTTGTGAC
CCCGTCTTCCAGGAGCTGGTGGAGCTGAAGCATGCGGCCTCCTGCCACCCCATGCACCGC
GGTGGACCAGGGAGAAGGCAGGGTCGAGGGCGGGGCCGGGCCAGGAGGCCCCCTGGG
AAGCCCAACCCCAGACGGCCCAAGGACAAGATGTCAGCCCTGGCCGCCTACTTTGTATAA

**3xFLAG-TOP3B Y336F**

ATGggttcttctgactacaaagaccatgacggtgattataaagatcatgacatcgattacaaggatgacgatgacaagggttcttctA
AGACTGTGCTCATGGTTGCTGAAAAGCCGTCCTTGGCACAGTCAATTGCCAAAATCCTCTC
TAGAGGGAGCCTGTCCTCACACAAAGGGCTGAACGGGGCCTGCTCAGTCCACGAGTACAC
TGGGACCTTTGCTGGCCAGCCAGTGCGCTTCAAGATGACGTCTGTCTGTGGTCACGTGAT
GACCCTGGATTTCCTGGGAAAATACAACAAATGGGACAAAGTGGACCCCGCAGAACTGTTC
AGCCAAGCTCCCACGGAGAAGAAAGAAGCTAACCCCAAGCTGAACATGGTGAAGTTCCTG
CAGGTGGAGGGCAGAGGCTGCGACTACATCGTGCTGTGGCTGGACTGCGACAAGGAGGG
GGAGAACATCTGCTTTGAGGTTCTTGATGCTGTTCTGCCCGTCATGAACAAGGCCCATGGT
GGCGAGAAGACCGTGTTCCGGGCCAGGTTTAGCTCCATCACGGACACAGACATCTGTAATG
CCATGGCCTGCCTAGGCGAGCCTGACCACAACGAGGCGCTCTCAGTGGATGCTCGCCAG
GAGCTGGACCTGCGAATCGGCTGTGCATTCACCAGGTTTCAGACTAAATATTTCCAGGGGA
AATACGGTGATTTAGACAGCTCTCTCATCTCCTTTGGGCCGTGTCAGACTCCAACCCTGGG
ATTCTGTGTGGAGAGACATGATAAAATCCAGTCCTTCAAACCAGAGACCTACTGGGTGCTGC
AGGCCAAGGTTAACACTGACAAAGACAGATCTCTCCTTTTGGACTGGGACCGAGTAAGAGT
GTTTGACCGGGAGATCGCACAGATGTTTTTAAACATGACAAAGCTGGAGAAGGAAGCCCAG
GTGGAGGCCACAAGCAGGAAAGAAAAGGCCAAGCAGAGGCCCCTGGCCCTGAACACTGT
GGAGATGCTGCGTGTGGCCAGCTCTTCTCTGGGCATGGGGCCGCAGCACGCCATGCAGA
CGGCTGAGCGGCTCTACACGCAAGGCTACATCAGCtttCCACGGACAGAGACCACCCACTAC
CCTGAGAACTTTGACCTGAAGGGCTCTCTGCGGCAGCAGGCCAACCACCCCTACTGGGCC
GACACGGTGAAGCGGTTGTTAGCAGAAGGTATCAACCGCCCGCGGAAAGGCCATGACGCC
GGCGACCATCCCCCCATCACCCCCATGAAGTCTGCCACAGAGGCCGAATTAGGGGGTGAC
GCGTGGCGGCTCTATGAGTACATCACCAGACACTTCATCGCCACGGTCAGCCATGACTGCA
AGTACCTGCAGAGCACCATCTCCTTCAGAATTGGGCCCGAGCTCTTCACCTGCTCCGGGAA
GACCGTCCTCTCACCAGGCTTCACGGAGGTCATGCCCTGGCAGAGCGTGCCCCTGGAGG
AGAGCCTGCCCACTTGCCAGCGGGGTGATGCCTTCCCTGTGGGCGAGGTGAAGATGCTG
GAGAAGCAGACGAACCCACCCGACTACCTGACGGAGGCCGAGCTCATCACGCTCATGGAG
AAGCATGGCATCGGCACGGATGCCAGCATCCCTGTGCATATCAACAACATCTGCCAGCGCA
ACTATGTCACGGTGGAGAGCGGGCGCCGGCTCAAGCCCACCAACCTCGGCATCGTCCTG
GTGCACGGCTACTATAAGATTGATGCAGAGCTGGTGCTCCCCACCATCCGCAGTGCAGTGG
AGAAGCAGCTGAACCTGATCGCCCAGGGCAAGGCCGACTACCGCCAGGTCCTGGGCCAC
ACCCTGGACGTGTTCAAGAGGAAGTTCCACTACTTTGTCGACTCCATTGCTGGCATGGATG
AGTTGATGGAGGTGTCTTTCTCGCCCCTGGCGGCCACAGGCAAGCCCCTCTCACGCTGTG
GGAAGTGCCACCGCTTCATGAAGTACATCCAGGCCAAGCCAAGCCGCCTGCACTGCTCCC
ACTGCGATGAGACCTACACGCTCCCCCAGAACGGCACCATCAAGCTCTACAAGGAGCTCC
GCTGCCCTCTGGATGACTTCGAGCTGGTCCTGTGGTCATCAGGCTCTCGGGGCAAGAGCT
ACCCGCTGTGCCCCTACTGCTACAACCACCCACCCTTCCGAGACATGAAGAAAGGCATGG
GCTGCAACGAGTGTACGCACCCCTCCTGCCAGCACTCGCTGAGCATGCTGGGCATCGGCC
AGTGCGTGGAATGTGAGAGCGGGGTGCTGGTGCTGGACCCCACCTCGGGCCCCAAGTGG
AAGGTGGCCTGCAACAAGTGCAACGTGGTAGCGCACTGCTTCGAGAACGCCCACCGCGTG
CGGGTGTCCGCCGACACCTGCAGTGTCTGTGAGGCCGCCTTGCTTGATGTGGACTTCAAC
AAGGCCAAGTCCCCACTCCCGGGCGATGAGACGCAGCACATGGGCTGCGTCTTTTGTGAC
CCCGTCTTCCAGGAGCTGGTGGAGCTGAAGCATGCGGCCTCCTGCCACCCCATGCACCGC
GGTGGACCAGGGAGAAGGCAGGGTCGAGGGCGGGGCCGGGCCAGGAGGCCCCCTGGG
AAGCCCAACCCCAGACGGCCCAAGGACAAGATGTCAGCCCTGGCCGCCTACTTTGTATAA

**3xFLAG-TOP3B R338W**

ATGggttcttctgactacaaagaccatgacggtgattataaagatcatgacatcgattacaaggatgacgatgacaagggttcttctA
AGACTGTGCTCATGGTTGCTGAAAAGCCGTCCTTGGCACAGTCAATTGCCAAAATCCTCTC
TAGAGGGAGCCTGTCCTCACACAAAGGGCTGAACGGGGCCTGCTCAGTCCACGAGTACAC
TGGGACCTTTGCTGGCCAGCCAGTGCGCTTCAAGATGACGTCTGTCTGTGGTCACGTGAT
GACCCTGGATTTCCTGGGAAAATACAACAAATGGGACAAAGTGGACCCCGCAGAACTGTTC

AGCCAAGCTCCCACGGAGAAGAAAGAAGCTAACCCCAAGCTGAACATGGTGAAGTTCCTG
CAGGTGGAGGGCAGAGGCTGCGACTACATCGTGCTGTGGCTGGACTGCGACAAGGAGGG
GGAGAACATCTGCTTTGAGGTTCTTGATGCTGTTCTGCCCGTCATGAACAAGGCCCATGGT
GGCGAGAAGACCGTGTTCCGGGCCAGGTTTAGCTCCATCACGGACACAGACATCTGTAATG
CCATGGCCTGCCTAGGCGAGCCTGACCACAACGAGGCGCTCTCAGTGGATGCTCGCCAG
GAGCTGGACCTGCGAATCGGCTGTGCATTCACCAGGTTTCAGACTAAATATTTCCAGGGGA
AATACGGTGATTTAGACAGCTCTCTCATCTCCTTTGGGCCGTGTCAGACTCCAACCCTGGG
ATTCTGTGTGGAGAGACATGATAAAATCCAGTCCTTCAAACCAGAGACCTACTGGGTGCTGC
AGGCCAAGGTTAACACTGACAAAGACAGATCTCTCCTTTTGGACTGGGACCGAGTAAGAGT
GTTTGACCGGGAGATCGCACAGATGTTTTTAAACATGACAAAGCTGGAGAAGGAAGCCCAG
GTGGAGGCCACAAGCAGGAAAGAAAAGGCCAAGCAGAGGCCCCTGGCCCTGAACACTGT
GGAGATGCTGCGTGTGGCCAGCTCTTCTCTGGGCATGGGGCCGCAGCACGCCATGCAGA
CGGCTGAGCGGCTCTACACGCAAGGCTACATCAGCTACCCAtggACAGAGACCACCCACTA
CCCTGAGAACTTTGACCTGAAGGGCTCTCTGCGGCAGCAGGCCAACCACCCCTACTGGGC
CGACACGGTGAAGCGGTTGTTAGCAGAAGGTATCAACCGCCCGCGGAAAGGCCATGACGC
CGGCGACCATCCCCCCATCACCCCCATGAAGTCTGCCACAGAGGCCGAATTAGGGGGTGA
CGCGTGGCGGCTCTATGAGTACATCACCAGACACTTCATCGCCACGGTCAGCCATGACTGC
AAGTACCTGCAGAGCACCATCTCCTTCAGAATTGGGCCCGAGCTCTTCACCTGCTCCGGGA
AGACCGTCCTCTCACCAGGCTTCACGGAGGTCATGCCCTGGCAGAGCGTGCCCCTGGAG
GAGAGCCTGCCCACTTGCCAGCGGGGTGATGCCTTCCCTGTGGGCGAGGTGAAGATGCT
GGAGAAGCAGACGAACCCACCCGACTACCTGACGGAGGCCGAGCTCATCACGCTCATGGA
GAAGCATGGCATCGGCACGGATGCCAGCATCCCTGTGCATATCAACAACATCTGCCAGCGC
AACTATGTCACGGTGGAGAGCGGGCGCCGGCTCAAGCCCACCAACCTCGGCATCGTCCTG
GTGCACGGCTACTATAAGATTGATGCAGAGCTGGTGCTCCCCACCATCCGCAGTGCAGTGG
AGAAGCAGCTGAACCTGATCGCCCAGGGCAAGGCCGACTACCGCCAGGTCCTGGGCCAC
ACCCTGGACGTGTTCAAGAGGAAGTTCCACTACTTTGTCGACTCCATTGCTGGCATGGATG
AGTTGATGGAGGTGTCTTTCTCGCCCCTGGCGGCCACAGGCAAGCCCCTCTCACGCTGTG
GGAAGTGCCACCGCTTCATGAAGTACATCCAGGCCAAGCCAAGCCGCCTGCACTGCTCCC
ACTGCGATGAGACCTACACGCTCCCCCAGAACGGCACCATCAAGCTCTACAAGGAGCTCC
GCTGCCCTCTGGATGACTTCGAGCTGGTCCTGTGGTCATCAGGCTCTCGGGGCAAGAGCT
ACCCGCTGTGCCCCTACTGCTACAACCACCCACCCTTCCGAGACATGAAGAAAGGCATGG
GCTGCAACGAGTGTACGCACCCCTCCTGCCAGCACTCGCTGAGCATGCTGGGCATCGGCC
AGTGCGTGGAATGTGAGAGCGGGGTGCTGGTGCTGGACCCCACCTCGGGCCCCAAGTGG
AAGGTGGCCTGCAACAAGTGCAACGTGGTAGCGCACTGCTTCGAGAACGCCCACCGCGTG
CGGGTGTCCGCCGACACCTGCAGTGTCTGTGAGGCCGCCTTGCTTGATGTGGACTTCAAC
AAGGCCAAGTCCCCACTCCCGGGCGATGAGACGCAGCACATGGGCTGCGTCTTTTGTGAC
CCCGTCTTCCAGGAGCTGGTGGAGCTGAAGCATGCGGCCTCCTGCCACCCCATGCACCGC
GGTGGACCAGGGAGAAGGCAGGGTCGAGGGCGGGGCCGGGCCAGGAGGCCCCCTGGG
AAGCCCAACCCCAGACGGCCCAAGGACAAGATGTCAGCCCTGGCCGCCTACTTTGTATAA

**3xFLAG-TOP3B R338W 1-823**

ATGggttcttctgactacaaagaccatgacggtgattataaagatcatgacatcgattacaaggatgacgatgacaagggttcttctA
AGACTGTGCTCATGGTTGCTGAAAAGCCGTCCTTGGCACAGTCAATTGCCAAAATCCTCTC
TAGAGGGAGCCTGTCCTCACACAAAGGGCTGAACGGGGCCTGCTCAGTCCACGAGTACAC
TGGGACCTTTGCTGGCCAGCCAGTGCGCTTCAAGATGACGTCTGTCTGTGGTCACGTGAT
GACCCTGGATTTCCTGGGAAAATACAACAAATGGGACAAAGTGGACCCCGCAGAACTGTTC
AGCCAAGCTCCCACGGAGAAGAAAGAAGCTAACCCCAAGCTGAACATGGTGAAGTTCCTG
CAGGTGGAGGGCAGAGGCTGCGACTACATCGTGCTGTGGCTGGACTGCGACAAGGAGGG
GGAGAACATCTGCTTTGAGGTTCTTGATGCTGTTCTGCCCGTCATGAACAAGGCCCATGGT
GGCGAGAAGACCGTGTTCCGGGCCAGGTTTAGCTCCATCACGGACACAGACATCTGTAATG
CCATGGCCTGCCTAGGCGAGCCTGACCACAACGAGGCGCTCTCAGTGGATGCTCGCCAG

GAGCTGGACCTGCGAATCGGCTGTGCATTCACCAGGTTTCAGACTAAATATTTCCAGGGGA
AATACGGTGATTTAGACAGCTCTCTCATCTCCTTTGGGCCGTGTCAGACTCCAACCCTGGG
ATTCTGTGTGGAGAGACATGATAAAATCCAGTCCTTCAAACCAGAGACCTACTGGGTGCTGC
AGGCCAAGGTTAACACTGACAAAGACAGATCTCTCCTTTTGGACTGGGACCGAGTAAGAGT
GTTTGACCGGGAGATCGCACAGATGTTTTTAAACATGACAAAGCTGGAGAAGGAAGCCCAG
GTGGAGGCCACAAGCAGGAAAGAAAAGGCCAAGCAGAGGCCCCTGGCCCTGAACACTGT
GGAGATGCTGCGTGTGGCCAGCTCTTCTCTGGGCATGGGGCCGCAGCACGCCATGCAGA
CGGCTGAGCGGCTCTACACGCAAGGCTACATCAGCTACCCAtggACAGAGACCACCCACTA
CCCTGAGAACTTTGACCTGAAGGGCTCTCTGCGGCAGCAGGCCAACCACCCCTACTGGGC
CGACACGGTGAAGCGGTTGTTAGCAGAAGGTATCAACCGCCCGCGGAAAGGCCATGACGC
CGGCGACCATCCCCCCATCACCCCCATGAAGTCTGCCACAGAGGCCGAATTAGGGGGTGA
CGCGTGGCGGCTCTATGAGTACATCACCAGACACTTCATCGCCACGGTCAGCCATGACTGC
AAGTACCTGCAGAGCACCATCTCCTTCAGAATTGGGCCCGAGCTCTTCACCTGCTCCGGGA
AGACCGTCCTCTCACCAGGCTTCACGGAGGTCATGCCCTGGCAGAGCGTGCCCCTGGAG
GAGAGCCTGCCCACTTGCCAGCGGGGTGATGCCTTCCCTGTGGGCGAGGTGAAGATGCT
GGAGAAGCAGACGAACCCACCCGACTACCTGACGGAGGCCGAGCTCATCACGCTCATGGA
GAAGCATGGCATCGGCACGGATGCCAGCATCCCTGTGCATATCAACAACATCTGCCAGCGC
AACTATGTCACGGTGGAGAGCGGGCGCCGGCTCAAGCCCACCAACCTCGGCATCGTCCTG
GTGCACGGCTACTATAAGATTGATGCAGAGCTGGTGCTCCCCACCATCCGCAGTGCAGTGG
AGAAGCAGCTGAACCTGATCGCCCAGGGCAAGGCCGACTACCGCCAGGTCCTGGGCCAC
ACCCTGGACGTGTTCAAGAGGAAGTTCCACTACTTTGTCGACTCCATTGCTGGCATGGATG
AGTTGATGGAGGTGTCTTTCTCGCCCCTGGCGGCCACAGGCAAGCCCCTCTCACGCTGTG
GGAAGTGCCACCGCTTCATGAAGTACATCCAGGCCAAGCCAAGCCGCCTGCACTGCTCCC
ACTGCGATGAGACCTACACGCTCCCCCAGAACGGCACCATCAAGCTCTACAAGGAGCTCC
GCTGCCCTCTGGATGACTTCGAGCTGGTCCTGTGGTCATCAGGCTCTCGGGGCAAGAGCT
ACCCGCTGTGCCCCTACTGCTACAACCACCCACCCTTCCGAGACATGAAGAAAGGCATGG
GCTGCAACGAGTGTACGCACCCCTCCTGCCAGCACTCGCTGAGCATGCTGGGCATCGGCC
AGTGCGTGGAATGTGAGAGCGGGGTGCTGGTGCTGGACCCCACCTCGGGCCCCAAGTGG
AAGGTGGCCTGCAACAAGTGCAACGTGGTAGCGCACTGCTTCGAGAACGCCCACCGCGTG
CGGGTGTCCGCCGACACCTGCAGTGTCTGTGAGGCCGCCTTGCTTGATGTGGACTTCAAC
AAGGCCAAGTCCCCACTCCCGGGCGATGAGACGCAGCACATGGGCTGCGTCTTTTGTGAC
CCCGTCTTCCAGGAGCTGGTGGAGCTGAAGCATGCGGCCTCCTGCCACCCCATGCACTAA

**3xFLAG-TOP3B R338W 1-612**

ATGggttcttctgactacaaagaccatgacggtgattataaagatcatgacatcgattacaaggatgacgatgacaagggttcttctA
AGACTGTGCTCATGGTTGCTGAAAAGCCGTCCTTGGCACAGTCAATTGCCAAAATCCTCTC
TAGAGGGAGCCTGTCCTCACACAAAGGGCTGAACGGGGCCTGCTCAGTCCACGAGTACAC
TGGGACCTTTGCTGGCCAGCCAGTGCGCTTCAAGATGACGTCTGTCTGTGGTCACGTGAT
GACCCTGGATTTCCTGGGAAAATACAACAAATGGGACAAAGTGGACCCCGCAGAACTGTTC
AGCCAAGCTCCCACGGAGAAGAAAGAAGCTAACCCCAAGCTGAACATGGTGAAGTTCCTG
CAGGTGGAGGGCAGAGGCTGCGACTACATCGTGCTGTGGCTGGACTGCGACAAGGAGGG
GGAGAACATCTGCTTTGAGGTTCTTGATGCTGTTCTGCCCGTCATGAACAAGGCCCATGGT
GGCGAGAAGACCGTGTTCCGGGCCAGGTTTAGCTCCATCACGGACACAGACATCTGTAATG
CCATGGCCTGCCTAGGCGAGCCTGACCACAACGAGGCGCTCTCAGTGGATGCTCGCCAG
GAGCTGGACCTGCGAATCGGCTGTGCATTCACCAGGTTTCAGACTAAATATTTCCAGGGGA
AATACGGTGATTTAGACAGCTCTCTCATCTCCTTTGGGCCGTGTCAGACTCCAACCCTGGG
ATTCTGTGTGGAGAGACATGATAAAATCCAGTCCTTCAAACCAGAGACCTACTGGGTGCTGC
AGGCCAAGGTTAACACTGACAAAGACAGATCTCTCCTTTTGGACTGGGACCGAGTAAGAGT
GTTTGACCGGGAGATCGCACAGATGTTTTTAAACATGACAAAGCTGGAGAAGGAAGCCCAG
GTGGAGGCCACAAGCAGGAAAGAAAAGGCCAAGCAGAGGCCCCTGGCCCTGAACACTGT
GGAGATGCTGCGTGTGGCCAGCTCTTCTCTGGGCATGGGGCCGCAGCACGCCATGCAGA

CGGCTGAGCGGCTCTACACGCAAGGCTACATCAGCTACCCAtggACAGAGACCACCCACTA
CCCTGAGAACTTTGACCTGAAGGGCTCTCTGCGGCAGCAGGCCAACCACCCCTACTGGGC
CGACACGGTGAAGCGGTTGTTAGCAGAAGGTATCAACCGCCCGCGGAAAGGCCATGACGC
CGGCGACCATCCCCCCATCACCCCCATGAAGTCTGCCACAGAGGCCGAATTAGGGGGTGA
CGCGTGGCGGCTCTATGAGTACATCACCAGACACTTCATCGCCACGGTCAGCCATGACTGC
AAGTACCTGCAGAGCACCATCTCCTTCAGAATTGGGCCCGAGCTCTTCACCTGCTCCGGGA
AGACCGTCCTCTCACCAGGCTTCACGGAGGTCATGCCCTGGCAGAGCGTGCCCCTGGAG
GAGAGCCTGCCCACTTGCCAGCGGGGTGATGCCTTCCCTGTGGGCGAGGTGAAGATGCT
GGAGAAGCAGACGAACCCACCCGACTACCTGACGGAGGCCGAGCTCATCACGCTCATGGA
GAAGCATGGCATCGGCACGGATGCCAGCATCCCTGTGCATATCAACAACATCTGCCAGCGC
AACTATGTCACGGTGGAGAGCGGGCGCCGGCTCAAGCCCACCAACCTCGGCATCGTCCTG
GTGCACGGCTACTATAAGATTGATGCAGAGCTGGTGCTCCCCACCATCCGCAGTGCAGTGG
AGAAGCAGCTGAACCTGATCGCCCAGGGCAAGGCCGACTACCGCCAGGTCCTGGGCCAC
ACCCTGGACGTGTTCAAGAGGAAGTTCCACTACTTTGTCGACTCCATTGCTGGCATGGATG
AGTTGATGGAGGTGTCTTTCTCGTAA

**TOP3B-3xFLAG ZnF (D669A)**

ATGAAGACTGTGCTCATGGTTGCTGAAAAGCCGTCCTTGGCACAGTCAATTGCCAAAATCCT
CTCTAGAGGGAGCCTGTCCTCACACAAAGGGCTGAACGGGGCCTGCTCAGTCCACGAGTA
CACTGGGACCTTTGCTGGCCAGCCAGTGCGCTTCAAGATGACGTCTGTCTGTGGTCACGT
GATGACCCTGGATTTCCTGGGAAAATACAACAAATGGGACAAAGTGGACCCCGCAGAACTG
TTCAGCCAAGCTCCCACGGAGAAGAAAGAAGCTAACCCCAAGCTGAACATGGTGAAGTTCC
TGCAGGTGGAGGGCAGAGGCTGCGACTACATCGTGCTGTGGCTGGACTGCGACAAGGAG
GGGGAGAACATCTGCTTTGAGGTTCTTGATGCTGTTCTGCCCGTCATGAACAAGGCCCATG
GTGGCGAGAAGACCGTGTTCCGGGCCAGGTTTAGCTCCATCACGGACACAGACATCTGTA
ATGCCATGGCCTGCCTAGGCGAGCCTGACCACAACGAGGCGCTCTCAGTGGATGCTCGCC
AGGAGCTGGACCTGCGAATCGGCTGTGCATTCACCAGGTTTCAGACTAAATATTTCCAGGG
GAAATACGGTGATTTAGACAGCTCTCTCATCTCCTTTGGGCCGTGTCAGACTCCAACCCTG
GGATTCTGTGTGGAGAGACATGATAAAATCCAGTCCTTCAAACCAGAGACCTACTGGGTGC
TGCAGGCCAAGGTTAACACTGACAAAGACAGATCTCTCCTTTTGGACTGGGACCGAGTAAG
AGTGTTTGACCGGGAGATCGCACAGATGTTTTTAAACATGACAAAGCTGGAGAAGGAAGCC
CAGGTGGAGGCCACAAGCAGGAAAGAAAAGGCCAAGCAGAGGCCCCTGGCCCTGAACAC
TGTGGAGATGCTGCGTGTGGCCAGCTCTTCTCTGGGCATGGGGCCGCAGCACGCCATGCA
GACGGCTGAGCGGCTCTACACGCAAGGCTACATCAGCTACCCACGGACAGAGACCACCCA
CTACCCTGAGAACTTTGACCTGAAGGGCTCTCTGCGGCAGCAGGCCAACCACCCCTACTG
GGCCGACACGGTGAAGCGGTTGTTAGCAGAAGGTATCAACCGCCCGCGGAAAGGCCATGA
CGCCGGCGACCATCCCCCCATCACCCCCATGAAGTCTGCCACAGAGGCCGAATTAGGGGG
TGACGCGTGGCGGCTCTATGAGTACATCACCAGACACTTCATCGCCACGGTCAGCCATGAC
TGCAAGTACCTGCAGAGCACCATCTCCTTCAGAATTGGGCCCGAGCTCTTCACCTGCTCCG
GGAAGACCGTCCTCTCACCAGGCTTCACGGAGGTCATGCCCTGGCAGAGCGTGCCCCTG
GAGGAGAGCCTGCCCACTTGCCAGCGGGGTGATGCCTTCCCTGTGGGCGAGGTGAAGAT
GCTGGAGAAGCAGACGAACCCACCCGACTACCTGACGGAGGCCGAGCTCATCACGCTCAT
GGAGAAGCATGGCATCGGCACGGATGCCAGCATCCCTGTGCATATCAACAACATCTGCCAG
CGCAACTATGTCACGGTGGAGAGCGGGCGCCGGCTCAAGCCCACCAACCTCGGCATCGT
CCTGGTGCACGGCTACTATAAGATTGATGCAGAGCTGGTGCTCCCCACCATCCGCAGTGCA
GTGGAGAAGCAGCTGAACCTGATCGCCCAGGGCAAGGCCGACTACCGCCAGGTCCTGGG
CCACACCCTGGACGTGTTCAAGAGGAAGTTCCACTACTTTGTCGACTCCATTGCTGGCATG
GATGAGTTGATGGAGGTGTCTTTCTCGCCCCTGGCGGCCACAGGCAAGCCCCTCTCACGC
TGTGGGAAGTGCCACCGCTTCATGAAGTACATCCAGGCCAAGCCAAGCCGCCTGCACTGC
TCCCACTGCGATGAGACCTACACGCTCCCCCAGAACGGCACCATCAAGCTCTACAAGGAG
CTCCGCTGCCCTCTGgccGACTTCGAGCTGGTCCTGTGGTCATCAGGCTCTCGGGGCAAGA

GCTACCCGCTGTGCCCCTACTGCTACAACCACCCACCCTTCCGAGACATGAAGAAAGGCAT
GGGCTGCAACGAGTGTACGCACCCCTCCTGCCAGCACTCGCTGAGCATGCTGGGCATCGG
CCAGTGCGTGGAATGTGAGAGCGGGGTGCTGGTGCTGGACCCCACCTCGGGCCCCAAGT
GGAAGGTGGCCTGCAACAAGTGCAACGTGGTAGCGCACTGCTTCGAGAACGCCCACCGC
GTGCGGGTGTCCGCCGACACCTGCAGTGTCTGTGAGGCCGCCTTGCTTGATGTGGACTTC
AACAAGGCCAAGTCCCCACTCCCGGGCGATGAGACGCAGCACATGGGCTGCGTCTTTTGT
GACCCCGTCTTCCAGGAGCTGGTGGAGCTGAAGCATGCGGCCTCCTGCCACCCCATGCAC
CGCGGTGGACCAGGGAGAAGGCAGGGTCGAGGGCGGGGCCGGGCCAGGAGGCCCCCT
GGGAAGCCCAACCCCAGACGGCCCAAGGACAAGATGTCAGCCCTGGCCGCCTACTTTGTA
AGCGGACCGACGCGTACGCGGCCGCTCGAGCAGAAACTCATCTCAGAAGAGGATCTGGCA
GCAAATGATATCCTGgactacaaagaccatgacggtgattataaagatcatgacatcgattacaaggatgacgatgacaag
GTTTAA

**TOP3B-3xFLAG ZnF (C688A/C691A)**

ATGAAGACTGTGCTCATGGTTGCTGAAAAGCCGTCCTTGGCACAGTCAATTGCCAAAATCCT
CTCTAGAGGGAGCCTGTCCTCACACAAAGGGCTGAACGGGGCCTGCTCAGTCCACGAGTA
CACTGGGACCTTTGCTGGCCAGCCAGTGCGCTTCAAGATGACGTCTGTCTGTGGTCACGT
GATGACCCTGGATTTCCTGGGAAAATACAACAAATGGGACAAAGTGGACCCCGCAGAACTG
TTCAGCCAAGCTCCCACGGAGAAGAAAGAAGCTAACCCCAAGCTGAACATGGTGAAGTTCC
TGCAGGTGGAGGGCAGAGGCTGCGACTACATCGTGCTGTGGCTGGACTGCGACAAGGAG
GGGGAGAACATCTGCTTTGAGGTTCTTGATGCTGTTCTGCCCGTCATGAACAAGGCCCATG
GTGGCGAGAAGACCGTGTTCCGGGCCAGGTTTAGCTCCATCACGGACACAGACATCTGTA
ATGCCATGGCCTGCCTAGGCGAGCCTGACCACAACGAGGCGCTCTCAGTGGATGCTCGCC
AGGAGCTGGACCTGCGAATCGGCTGTGCATTCACCAGGTTTCAGACTAAATATTTCCAGGG
GAAATACGGTGATTTAGACAGCTCTCTCATCTCCTTTGGGCCGTGTCAGACTCCAACCCTG
GGATTCTGTGTGGAGAGACATGATAAAATCCAGTCCTTCAAACCAGAGACCTACTGGGTGC
TGCAGGCCAAGGTTAACACTGACAAAGACAGATCTCTCCTTTTGGACTGGGACCGAGTAAG
AGTGTTTGACCGGGAGATCGCACAGATGTTTTTAAACATGACAAAGCTGGAGAAGGAAGCC
CAGGTGGAGGCCACAAGCAGGAAAGAAAAGGCCAAGCAGAGGCCCCTGGCCCTGAACAC
TGTGGAGATGCTGCGTGTGGCCAGCTCTTCTCTGGGCATGGGGCCGCAGCACGCCATGCA
GACGGCTGAGCGGCTCTACACGCAAGGCTACATCAGCTACCCACGGACAGAGACCACCCA
CTACCCTGAGAACTTTGACCTGAAGGGCTCTCTGCGGCAGCAGGCCAACCACCCCTACTG
GGCCGACACGGTGAAGCGGTTGTTAGCAGAAGGTATCAACCGCCCGCGGAAAGGCCATGA
CGCCGGCGACCATCCCCCCATCACCCCCATGAAGTCTGCCACAGAGGCCGAATTAGGGGG
TGACGCGTGGCGGCTCTATGAGTACATCACCAGACACTTCATCGCCACGGTCAGCCATGAC
TGCAAGTACCTGCAGAGCACCATCTCCTTCAGAATTGGGCCCGAGCTCTTCACCTGCTCCG
GGAAGACCGTCCTCTCACCAGGCTTCACGGAGGTCATGCCCTGGCAGAGCGTGCCCCTG
GAGGAGAGCCTGCCCACTTGCCAGCGGGGTGATGCCTTCCCTGTGGGCGAGGTGAAGAT
GCTGGAGAAGCAGACGAACCCACCCGACTACCTGACGGAGGCCGAGCTCATCACGCTCAT
GGAGAAGCATGGCATCGGCACGGATGCCAGCATCCCTGTGCATATCAACAACATCTGCCAG
CGCAACTATGTCACGGTGGAGAGCGGGCGCCGGCTCAAGCCCACCAACCTCGGCATCGT
CCTGGTGCACGGCTACTATAAGATTGATGCAGAGCTGGTGCTCCCCACCATCCGCAGTGCA
GTGGAGAAGCAGCTGAACCTGATCGCCCAGGGCAAGGCCGACTACCGCCAGGTCCTGGG
CCACACCCTGGACGTGTTCAAGAGGAAGTTCCACTACTTTGTCGACTCCATTGCTGGCATG
GATGAGTTGATGGAGGTGTCTTTCTCGCCCCTGGCGGCCACAGGCAAGCCCCTCTCACGC
TGTGGGAAGTGCCACCGCTTCATGAAGTACATCCAGGCCAAGCCAAGCCGCCTGCACTGC
TCCCACTGCGATGAGACCTACACGCTCCCCCAGAACGGCACCATCAAGCTCTACAAGGAG
CTCCGCTGCCCTCTGGATGACTTCGAGCTGGTCCTGTGGTCATCAGGCTCTCGGGGCAAG
AGCTACCCGCTGgccCCCTACgccTACAACCACCCACCCTTCCGAGACATGAAGAAAGGCAT
GGGCTGCAACGAGTGTACGCACCCCTCCTGCCAGCACTCGCTGAGCATGCTGGGCATCGG

CCAGTGCGTGGAATGTGAGAGCGGGGTGCTGGTGCTGGACCCCACCTCGGCCCCAAGT
GGAAGGTGGCCTGCAACAAGTGCAACGTGGTAGCGCACTGCTTCGAGAACGCCCACCGC
GTGCGGGTGTCCGCCGACACCTGCAGTGTCTGTGAGGCCGCCTTGCTTGATGTGGACTTC
AACAAGGCCAAGTCCCCACTCCCGGGCGATGAGACGCAGCACATGGGCTGCGTCTTTTGT
GACCCCGTCTTCCAGGAGCTGGTGGAGCTGAAGCATGCGGCCTCCTGCCACCCCATGCAC
CGCGGTGGACCAGGGAGAAGGCAGGGTCGAGGGCGGGGCCGGGCCAGGAGGCCCCCT
GGGAAGCCCAACCCCAGACGGCCCAAGGACAAGATGTCAGCCCTGGCCGCCTACTTTGTA
AGCGGACCGACGCGTACGCGGCCGCTCGAGCAGAAACTCATCTCAGAAGAGGATCTGGCA
GCAAATGATATCCTGgactacaaagaccatgacggtgattataaagatcatgacatcgattacaaggatgacgatgacaag
GTTTAA

**TOP3B-3xFLAG ZnF (C706A/C709A)**

ATGAAGACTGTGCTCATGGTTGCTGAAAAGCCGTCCTTGGCACAGTCAATTGCCAAAATCCT
CTCTAGAGGGAGCCTGTCCTCACACAAAGGGCTGAACGGGGCCTGCTCAGTCCACGAGTA
CACTGGGACCTTTGCTGGCCAGCCAGTGCGCTTCAAGATGACGTCTGTCTGTGGTCACGT
GATGACCCTGGATTTCCTGGGAAAATACAACAAATGGGACAAAGTGGACCCCGCAGAACTG
TTCAGCCAAGCTCCCACGGAGAAGAAAGAAGCTAACCCCAAGCTGAACATGGTGAAGTTCC
TGCAGGTGGAGGGCAGAGGCTGCGACTACATCGTGCTGTGGCTGGACTGCGACAAGGAG
GGGGAGAACATCTGCTTTGAGGTTCTTGATGCTGTTCTGCCCGTCATGAACAAGGCCCATG
GTGGCGAGAAGACCGTGTTCCGGGCCAGGTTTAGCTCCATCACGGACACAGACATCTGTA
ATGCCATGGCCTGCCTAGGCGAGCCTGACCACAACGAGGCGCTCTCAGTGGATGCTCGCC
AGGAGCTGGACCTGCGAATCGGCTGTGCATTCACCAGGTTTCAGACTAAATATTTCCAGGG
GAAATACGGTGATTTAGACAGCTCTCTCATCTCCTTTGGGCCGTGTCAGACTCCAACCCTG
GGATTCTGTGTGGAGAGACATGATAAAATCCAGTCCTTCAAACCAGAGACCTACTGGGTGC
TGCAGGCCAAGGTTAACACTGACAAAGACAGATCTCTCCTTTTGGACTGGGACCGAGTAAG
AGTGTTTGACCGGGAGATCGCACAGATGTTTTTAAACATGACAAAGCTGGAGAAGGAAGCC
CAGGTGGAGGCCACAAGCAGGAAAGAAAAGGCCAAGCAGAGGCCCCTGGCCCTGAACAC
TGTGGAGATGCTGCGTGTGGCCAGCTCTTCTCTGGGCATGGGGCCGCAGCACGCCATGCA
GACGGCTGAGCGGCTCTACACGCAAGGCTACATCAGCTACCCACGGACAGAGACCACCCA
CTACCCTGAGAACTTTGACCTGAAGGGCTCTCTGCGGCAGCAGGCCAACCACCCCTACTG
GGCCGACACGGTGAAGCGGTTGTTAGCAGAAGGTATCAACCGCCCGCGGAAAGGCCATGA
CGCCGGCGACCATCCCCCCATCACCCCCATGAAGTCTGCCACAGAGGCCGAATTAGGGGG
TGACGCGTGGCGGCTCTATGAGTACATCACCAGACACTTCATCGCCACGGTCAGCCATGAC
TGCAAGTACCTGCAGAGCACCATCTCCTTCAGAATTGGGCCCGAGCTCTTCACCTGCTCCG
GGAAGACCGTCCTCTCACCAGGCTTCACGGAGGTCATGCCCTGGCAGAGCGTGCCCCTG
GAGGAGAGCCTGCCCACTTGCCAGCGGGGTGATGCCTTCCCTGTGGGCGAGGTGAAGAT
GCTGGAGAAGCAGACGAACCCACCCGACTACCTGACGGAGGCCGAGCTCATCACGCTCAT
GGAGAAGCATGGCATCGGCACGGATGCCAGCATCCCTGTGCATATCAACAACATCTGCCAG
CGCAACTATGTCACGGTGGAGAGCGGGCGCCGGCTCAAGCCCACCAACCTCGGCATCGT
CCTGGTGCACGGCTACTATAAGATTGATGCAGAGCTGGTGCTCCCCACCATCCGCAGTGCA
GTGGAGAAGCAGCTGAACCTGATCGCCCAGGGCAAGGCCGACTACCGCCAGGTCCTGGG
CCACACCCTGGACGTGTTCAAGAGGAAGTTCCACTACTTTGTCGACTCCATTGCTGGCATG
GATGAGTTGATGGAGGTGTCTTTCTCGCCCCTGGCGGCCACAGGCAAGCCCCTCTCACGC
TGTGGGAAGTGCCACCGCTTCATGAAGTACATCCAGGCCAAGCCAAGCCGCCTGCACTGC
TCCCACTGCGATGAGACCTACACGCTCCCCCAGAACGGCACCATCAAGCTCTACAAGGAG
CTCCGCTGCCCTCTGGATGACTTCGAGCTGGTCCTGTGGTCATCAGGCTCTCGGGGCAAG
AGCTACCCGCTGTGCCCCTACTGCTACAACCACCCACCCTTCCGAGACATGAAGAAAGGCA
TGGGCgccAACGAGgccACGCACCCCTCCTGCCAGCACTCGCTGAGCATGCTGGGCATCGG
CCAGTGCGTGGAATGTGAGAGCGGGGTGCTGGTGCTGGACCCCACCTCGGCCCCAAGT
GGAAGGTGGCCTGCAACAAGTGCAACGTGGTAGCGCACTGCTTCGAGAACGCCCACCGC
GTGCGGGTGTCCGCCGACACCTGCAGTGTCTGTGAGGCCGCCTTGCTTGATGTGGACTTC

AACAAGGCCAAGTCCCCACTCCCGGGCGATGAGACGCAGCACATGGGCTGCGTCTTTTGT
GACCCCGTCTTCCAGGAGCTGGTGGAGCTGAAGCATGCGGCCTCCTGCCACCCCATGCAC
CGCGGTGGACCAGGGAGAAGGCAGGGTCGAGGGCGGGGCCGGGCCAGGAGGCCCCCT
GGGAAGCCCAACCCCAGACGGCCCAAGGACAAGATGTCAGCCCTGGCCGCCTACTTTGTA
AGCGGACCGACGCGTACGCGGCCGCTCGAGCAGAAACTCATCTCAGAAGAGGATCTGGCA
GCAAATGATATCCTGgactacaaagaccatgacggtgattataaagatcatgacatcgattacaaggatgacgatgacaag
GTTTAA

**mEGFP (used in pcDNA3.1(+))**
atggtgagcaagggcgaggagctgttcaccggggtggtgcccatcctggtcgagctggacggcgacgtaaacggccacaagttca
gcgtgtccggcgagggcgagggcgatgccacctacggcaagctgaccctgaagttcatctgcaccaccggcaagctgcccgtgcc
ctggcccaccctcgtgaccaccctgacctacggcgtgcagtgcttcagccgctaccccgaccacatgaagcagcacgacttcttcaa
gtccgccatgcccgaaggctacgtccaggagcgcaccatcttcttcaaggacgacggcaactacaagacccgcgccgaggtgaa
gttcgagggcgacaccctggtgaaccgcatcgagctgaagggcatcgacttcaaggaggacggcaacatcctggggcacaagct
ggagtacaactacaacagccacaacgtctatatcatggccgacaagcagaagaacggcatcaaggtgaacttcaagatccgcca
caacatcgaggacggcagcgtgcagctcgccgaccactaccagcagaacacccccatcggcgacggccccgtgctgctgcccga
caaccactacctgagcacccagtccaagctgagcaaagaccccaacgagaagcgcgatcacatggtcctgctggagttcgtgacc
gccgccgggatcactctcggcatggacgagctgtacaagtaa

**3xV5-Ubiquitn (used in pcDNA3.1(+))**
atggggaagcctatacccaatcctctcctgggggttggacagcaccggcagtggtagcggtggcaagcccatccccaacccccctgctg
ggcctggactccaccggctccggcagtggtggaaaacctattcccaacccactgctgggactggattctacaagcggatccatgcag
atcttcgtgaaaacccttaccggcaagaccatcacccttgaggtggagcccagtgacaccatcgaaaatgtgaaggccaagatcca
ggataaggaaggcattccccccgaccagcagaggctcatctttgcaggcaagcagctggaagatggccgtactctttctgactacaa
catccagaaggagtcgaccctgcacctggtcctgcgtctgagaggtggttag

**3xV5-Ubiquitn K6R, K27R, K29R, K33R, K63R (used in pcDNA3.1(+))**
atggggaagcctatacccaatcctctcctgggggttggacagcaccggcagtggtagcggtggcaagcccatccccaacccccctgctg
ggcctggactccaccggctccggcagtggtggaaaacctattcccaacccactgctgggactggattctacaagcggatccatgcag
atcttcgtgCGCacccttaccggcaagaccatcacccttgaggtggagcccagtgacaccatcgaaaatgtgCGCgccCGCat
ccaggatCGCgaaggcattccccccgaccagcagaggctcatctttgcaggcaagcagctggaagatggccgtactctttctgact
acaacatccagCGCgagtcgaccctgcacctggtcctgcgtctgagaggtggttag

**3xV5-Ubiquitn K11R, K48R (used in pcDNA3.1(+))**
atggggaagcctatacccaatcctctcctgggggttggacagcaccggcagtggtagcggtggcaagcccatccccaacccccctgctg
ggcctggactccaccggctccggcagtggtggaaaacctattcccaacccactgctgggactggattctacaagcggatccatgcag
atcttcgtgaaaacccttaccggcCGCaccatcacccttgaggtggagcccagtgacaccatcgaaaatgtgaaggccaagatcc
aggataaggaaggcattccccccgaccagcagaggctcatctttgcaggcCGCcagctggaagatggccgtactctttctgactac
aacatccagaaggagtcgaccctgcacctggtcctgcgtctgagaggtggttag

### <u>ORFs in plasmids used in longitudinal fluorescence microscopy and to generate dox-inducible stable cell lines</u>

**3xFLAG-mEGFP-NES-TOP3B WT (used in pGW1 and pcDNA5/FRT/TO)**
atgggaagcagcgactacaaagaccatgacggtgattataaagatcatgacatcgattacaaggatgacgatgacaagggttctgg
tgtgagcaagggcgaggagctgttcaccggggtggtgcccatcctggtcgagctggacggcgacgtaaacggccacaagttcagc
gtgtccggcgagggcgagggcgatgccacctacggcaagctgaccctgaagttcatctgcaccaccggcaagctgcccgtgccctg
gcccaccctcgtgaccaccctgacctacggcgtgcagtgcttcagccgctaccccgaccacatgaagcagcacgacttcttcaagtc
cgccatgcccgaaggctacgtccaggagcgcaccatcttcttcaaggacgacggcaactacaagacccgcgccgaggtgaagttc
gagggcgacaccctggtgaaccgcatcgagctgaagggcatcgacttcaaggaggacggcaacatcctggggcacaagctgga

gtacaactacaacagccacaacgtctatatcatggccgacaagcagaagaacggcatcaaggtgaacttcaagatccgccacaac
atcgaggacggcagcgtgcagctcgccgaccactaccagcagaacacccccatcggcgacggccccgtgctgctgcccgacaac
cactacctgagcacccagtccaagctgagcaaagaccccaacgagaagcgcgatcacatggtcctgctggagttcgtgaccgccg
ccgggatcactctcggcatggacgagctgtacaagggttctggtctgcagctgcctcccctggagcgcctgaccctggacggttcttct
AAGACTGTGCTCATGGTTGCTGAAAAGCCGTCCTTGGCACAGTCAATTGCCAAAATCCTCT
CTAGAGGGAGCCTGTCCTCACACAAAGGGCTGAACGGGGCCTGCTCAGTCCACGAGTACA
CTGGGACCTTTGCTGGCCAGCCAGTGCGCTTCAAGATGACGTCTGTCTGTGGTCACGTGA
TGACCCTGGATTTCCTGGGAAAATACAACAAATGGGACAAAGTGGACCCCGCAGAACTGTT
CAGCCAAGCTCCCACGGAGAAGAAAGAAGCTAACCCCAAGCTGAACATGGTGAAGTTCCT
GCAGGTGGAGGGCAGAGGCTGCGACTACATCGTGCTGTGGCTGGACTGCGACAAGGAGG
GGGAGAACATCTGCTTTGAGGTTCTTGATGCTGTTCTGCCCGTCATGAACAAGGCCCATGG
TGGCGAGAAGACCGTGTTCCGGGCCAGGTTTAGCTCCATCACGGACACAGACATCTGTAAT
GCCATGGCCTGCCTAGGCGAGCCTGACCACAACGAGGCGCTCTCAGTGGATGCTCGCCA
GGAGCTGGACCTGCGAATCGGCTGTGCATTCACCAGGTTTCAGACTAAATATTTCCAGGGG
AAATACGGTGATTTAGACAGCTCTCTCATCTCCTTTGGGCCGTGTCAGACTCCAACCCTGG
GATTCTGTGTGGAGAGACATGATAAAATCCAGTCCTTCAAACCAGAGACCTACTGGGTGCT
GCAGGCCAAGGTTAACACTGACAAAGACAGATCTCTCCTTTTGGACTGGGACCGAGTAAGA
GTGTTTGACCGGGAGATCGCACAGATGTTTTTAAACATGACAAAGCTGGAGAAGGAAGCCC
AGGTGGAGGCCACAAGCAGGAAAGAAAAGGCCAAGCAGAGGCCCCTGGCCCTGAACACT
GTGGAGATGCTGCGTGTGGCCAGCTCTTCTCTGGGCATGGGGCCGCAGCACGCCATGCA
GACGGCTGAGCGGCTCTACACGCAAGGCTACATCAGCTACCCACGGACAGAGACCACCCA
CTACCCTGAGAACTTTGACCTGAAGGGCTCTCTGCGGCAGCAGGCCAACCACCCCTACTG
GGCCGACACGGTGAAGCGGTTGTTAGCAGAAGGTATCAACCGCCCGCGGAAAGGCCATGA
CGCCGGCGACCATCCCCCCATCACCCCCATGAAGTCTGCCACAGAGGCCGAATTAGGGGG
TGACGCGTGGCGGCTCTATGAGTACATCACCAGACACTTCATCGCCACGGTCAGCCATGAC
TGCAAGTACCTGCAGAGCACCATCTCCTTCAGAATTGGGCCCGAGCTCTTCACCTGCTCCG
GGAAGACCGTCCTCTCACCAGGCTTCACGGAGGTCATGCCCTGGCAGAGCGTGCCCCTG
GAGGAGAGCCTGCCCACTTGCCAGCGGGGTGATGCCTTCCCTGTGGGCGAGGTGAAGAT
GCTGGAGAAGCAGACGAACCCACCCGACTACCTGACGGAGGCCGAGCTCATCACGCTCAT
GGAGAAGCATGGCATCGGCACGGATGCCAGCATCCCTGTGCATATCAACAACATCTGCCAG
CGCAACTATGTCACGGTGGAGAGCGGGCGCCGGCTCAAGCCCACCAACCTCGGCATCGT
CCTGGTGCACGGCTACTATAAGATTGATGCAGAGCTGGTGCTCCCCACCATCCGCAGTGCA
GTGGAGAAGCAGCTGAACCTGATCGCCCAGGGCAAGGCCGACTACCGCCAGGTCCTGGG
CCACACCCTGGACGTGTTCAAGAGGAAGTTCCACTACTTTGTCGACTCCATTGCTGGCATG
GATGAGTTGATGGAGGTGTCTTTCTCGCCCCTGGCGGCCACAGGCAAGCCCCTCTCACGC
TGTGGGAAGTGCCACCGCTTCATGAAGTACATCCAGGCCAAGCCAAGCCGCCTGCACTGC
TCCCACTGCGATGAGACCTACACGCTCCCCCAGAACGGCACCATCAAGCTCTACAAGGAG
CTCCGCTGCCCTCTGGATGACTTCGAGCTGGTCCTGTGGTCATCAGGCTCTCGGGGCAAG
AGCTACCCGCTGTGCCCCTACTGCTACAACCACCCACCCTTCCGAGACATGAAGAAAGGCA
TGGGCTGCAACGAGTGTACGCACCCCTCCTGCCAGCACTCGCTGAGCATGCTGGGCATCG
GCCAGTGCGTGGAATGTGAGAGCGGGGTGCTGGTGCTGGACCCCACCTCGGGCCCCAAG
TGGAAGGTGGCCTGCAACAAGTGCAACGTGGTAGCGCACTGCTTCGAGAACGCCCACCGC
GTGCGGGTGTCCGCCGACACCTGCAGTGTCTGTGAGGCCGCCTTGCTTGATGTGGACTTC
AACAAGGCCAAGTCCCCACTCCCGGGCGATGAGACGCAGCACATGGGCTGCGTCTTTTGT
GACCCCGTCTTCCAGGAGCTGGTGGAGCTGAAGCATGCGGCCTCCTGCCACCCCATGCAC
CGCGGTGGACCAGGGAGAAGGCAGGGTCGAGGGCGGGGCCGGGCCAGGAGGCCCCCT
GGGAAGCCCAACCCCAGACGGCCCAAGGACAAGATGTCAGCCCTGGCCGCCTACTTTGTA
TAA

**3xFLAG-mEGFP-NES-TOP3B Y336F (used in pGW1 and pcDNA5/FRT/TO)**

atgggaagcagcgactacaaagaccatgacggtgattataaagatcatgacatcgattacaaggatgacgatgacaagggttctgg
tgtgagcaagggcgaggagctgttcaccggggtggtgcccatcctggtcgagctggacggcgacgtaaacggccacaagttcagc
gtgtccggcgagggcgagggcgatgccacctacggcaagctgaccctgaagttcatctgcaccaccggcaagctgcccgtgccctg
gcccaccctcgtgaccaccctgacctacggcgtgcagtgcttcagccgctaccccgaccacatgaagcagcacgacttcttcaagtc
cgccatgcccgaaggctacgtccaggagcgcaccatcttcttcaaggacgacggcaactacaagacccgcgccgaggtgaagttc
gagggcgacaccctggtgaaccgcatcgagctgaagggcatcgacttcaaggaggacggcaacatcctggggcacaagctgga
gtacaactacaacagccacaacgtctatatcatggccgacaagcagaagaacggcatcaaggtgaacttcaagatccgccacaac
atcgaggacggcagcgtgcagctcgccgaccactaccagcagaacacccccatcggcgacggccccgtgctgctgcccgacaac
cactacctgagcacccagtccaagctgagcaaagaccccaacgagaagcgcgatcacatggtcctgctggagttcgtgaccgccg
ccgggatcactctcggcatggacgagctgtacaagggttctggtctgcagctgcctcccctggagcgcctgaccctggacggttcttct
AAGACTGTGCTCATGGTTGCTGAAAAGCCGTCCTTGGCACAGTCAATTGCCAAAATCCTCT
CTAGAGGGAGCCTGTCCTCACACAAAGGGCTGAACGGGGCCTGCTCAGTCCACGAGTACA
CTGGGACCTTTGCTGGCCAGCCAGTGCGCTTCAAGATGACGTCTGTCTGTGGTCACGTGA
TGACCCTGGATTTCCTGGGAAAATACAACAAATGGGACAAAGTGGACCCCGCAGAACTGTT
CAGCCAAGCTCCCACGGAGAAGAAAGAAGCTAACCCCAAGCTGAACATGGTGAAGTTCCT
GCAGGTGGAGGGCAGAGGCTGCGACTACATCGTGCTGTGGCTGGACTGCGACAAGGAGG
GGGAGAACATCTGCTTTGAGGTTCTTGATGCTGTTCTGCCCGTCATGAACAAGGCCCATGG
TGGCGAGAAGACCGTGTTCCGGGCCAGGTTTAGCTCCATCACGGACACAGACATCTGTAAT
GCCATGGCCTGCCTAGGCGAGCCTGACCACAACGAGGCGCTCTCAGTGGATGCTCGCCA
GGAGCTGGACCTGCGAATCGGCTGTGCATTCACCAGGTTTCAGACTAAATATTTCCAGGGG
AAATACGGTGATTTAGACAGCTCTCTCATCTCCTTTGGGCCGTGTCAGACTCCAACCCTGG
GATTCTGTGTGGAGAGACATGATAAAATCCAGTCCTTCAAACCAGAGACCTACTGGGTGCT
GCAGGCCAAGGTTAACACTGACAAAGACAGATCTCTCCTTTTGGACTGGGACCGAGTAAGA
GTGTTTGACCGGGAGATCGCACAGATGTTTTTAAACATGACAAAGCTGGAGAAGGAAGCCC
AGGTGGAGGCCACAAGCAGGAAAGAAAAGGCCAAGCAGAGGCCCCTGGCCCTGAACACT
GTGGAGATGCTGCGTGTGGCCAGCTCTTCTCTGGGCATGGGGCCGCAGCACGCCATGCA
GACGGCTGAGCGGCTCTACACGCAAGGCTACATCAGCtttCCACGGACAGAGACCACCCACT
ACCCTGAGAACTTTGACCTGAAGGGCTCTCTGCGGCAGCAGGCCAACCACCCCTACTGGG
CCGACACGGTGAAGCGGTTGTTAGCAGAAGGTATCAACCGCCCGCGGAAAGGCCATGACG
CCGGCGACCATCCCCCCATCACCCCCATGAAGTCTGCCACAGAGGCCGAATTAGGGGGTG
ACGCGTGGCGGCTCTATGAGTACATCACCAGACACTTCATCGCCACGGTCAGCCATGACTG
CAAGTACCTGCAGAGCACCATCTCCTTCAGAATTGGGCCCGAGCTCTTCACCTGCTCCGGG
AAGACCGTCCTCTCACCAGGCTTCACGGAGGTCATGCCCTGGCAGAGCGTGCCCCTGGAG
GAGAGCCTGCCCACTTGCCAGCGGGGTGATGCCTTCCCTGTGGGCGAGGTGAAGATGCT
GGAGAAGCAGACGAACCCACCCGACTACCTGACGGAGGCCGAGCTCATCACGCTCATGGA
GAAGCATGGCATCGGCACGGATGCCAGCATCCCTGTGCATATCAACAACATCTGCCAGCGC
AACTATGTCACGGTGGAGAGCGGGCGCCGGCTCAAGCCCACCAACCTCGGCATCGTCCTG
GTGCACGGCTACTATAAGATTGATGCAGAGCTGGTGCTCCCCACCATCCGCAGTGCAGTGG
AGAAGCAGCTGAACCTGATCGCCCAGGGCAAGGCCGACTACCGCCAGGTCCTGGGCCAC
ACCCTGGACGTGTTCAAGAGGAAGTTCCACTACTTTGTCGACTCCATTGCTGGCATGGATG
AGTTGATGGAGGTGTCTTTCTCGCCCCTGGCGGCCACAGGCAAGCCCCTCTCACGCTGTG
GGAAGTGCCACCGCTTCATGAAGTACATCCAGGCCAAGCCAAGCCGCCTGCACTGCTCCC
ACTGCGATGAGACCTACACGCTCCCCCAGAACGGCACCATCAAGCTCTACAAGGAGCTCC
GCTGCCCTCTGGATGACTTCGAGCTGGTCCTGTGGTCATCAGGCTCTCGGGGCAAGAGCT
ACCCGCTGTGCCCCTACTGCTACAACCACCCACCCTTCCGAGACATGAAGAAAGGCATGG
GCTGCAACGAGTGTACGCACCCCTCCTGCCAGCACTCGCTGAGCATGCTGGGCATCGGCC
AGTGCGTGGAATGTGAGAGCGGGGTGCTGGTGCTGGACCCCACCTCGGGCCCCAAGTGG
AAGGTGGCCTGCAACAAGTGCAACGTGGTAGCGCACTGCTTCGAGAACGCCCACCGCGTG
CGGGTGTCCGCCGACACCTGCAGTGTCTGTGAGGCCGCCTTGCTTGATGTGGACTTCAAC
AAGGCCAAGTCCCCACTCCCGGGCGATGAGACGCAGCACATGGGCTGCGTCTTTTGTGAC
CCCGTCTTCCAGGAGCTGGTGGAGCTGAAGCATGCGGCCTCCTGCCACCCCATGCACCGC

GGTGGACCAGGGAGAAGGCAGGGTCGAGGGCGGGGCCGGGCCAGGAGGCCCCCTGGG
AAGCCCAACCCCAGACGGCCCAAGGACAAGATGTCAGCCCTGGCCGCCTACTTTGTATAA

**3xFLAG-mEGFP-NES-TOP3B R338W (used in pGW1 and pcDNA5/FRT/TO)**

atgggaagcagcgactacaaagaccatgacggtgattataaagatcatgacatcgattacaaggatgacgatgacaagggttctgg
tgtgagcaagggcgaggagctgttcaccggggtggtgcccatcctggtcgagctggacggcgacgtaaacggccacaagttcagc
gtgtccggcgagggcgagggcgatgccacctacggcaagctgaccctgaagttcatctgcaccaccggcaagctgcccgtgccctg
gcccaccctcgtgaccaccctgacctacggcgtgcagtgcttcagccgctaccccgaccacatgaagcagcacgacttcttcaagtc
cgccatgcccgaaggctacgtccaggagcgcaccatcttcttcaaggacgacggcaactacaagacccgcgccgaggtgaagttc
gagggcgacaccctggtgaaccgcatcgagctgaagggcatcgacttcaaggaggacggcaacatcctggggcacaagctgga
gtacaactacaacagccacaacgtctatatcatggccgacaagcagaagaacggcatcaaggtgaacttcaagatccgccacaac
atcgaggacggcagcgtgcagctcgccgaccactaccagcagaacacccccatcggcgacggccccgtgctgctgcccgacaac
cactacctgagcacccagtccaagctgagcaaagaccccaacgagaagcgcgatcacatggtcctgctggagttcgtgaccgccg
ccgggatcactctcggcatggacgagctgtacaagggttctggtctgcagctgcctcccctggagcgcctgaccctggacggttcttct
AAGACTGTGCTCATGGTTGCTGAAAAGCCGTCCTTGGCACAGTCAATTGCCAAAATCCTCT
CTAGAGGGAGCCTGTCCTCACACAAAGGGCTGAACGGGGCCTGCTCAGTCCACGAGTACA
CTGGGACCTTTGCTGGCCAGCCAGTGCGCTTCAAGATGACGTCTGTCTGTGGTCACGTGA
TGACCCTGGATTTCCTGGGAAAATACAACAAATGGGACAAAGTGGACCCCGCAGAACTGTT
CAGCCAAGCTCCCACGGAGAAGAAAGAAGCTAACCCCAAGCTGAACATGGTGAAGTTCCT
GCAGGTGGAGGGCAGAGGCTGCGACTACATCGTGCTGTGGCTGGACTGCGACAAGGAGG
GGGAGAACATCTGCTTTGAGGTTCTTGATGCTGTTCTGCCCGTCATGAACAAGGCCCATGG
TGGCGAGAAGACCGTGTTCCGGGCCAGGTTTAGCTCCATCACGGACACAGACATCTGTAAT
GCCATGGCCTGCCTAGGCGAGCCTGACCACAACGAGGCGCTCTCAGTGGATGCTCGCCA
GGAGCTGGACCTGCGAATCGGCTGTGCATTCACCAGGTTTCAGACTAAATATTTCCAGGGG
AAATACGGTGATTTAGACAGCTCTCTCATCTCCTTTGGGCCGTGTCAGACTCCAACCCTGG
GATTCTGTGTGGAGAGACATGATAAAATCCAGTCCTTCAAACCAGAGACCTACTGGGTGCT
GCAGGCCAAGGTTAACACTGACAAAGACAGATCTCTCCTTTTGGACTGGGACCGAGTAAGA
GTGTTTGACCGGGAGATCGCACAGATGTTTTTAAACATGACAAAGCTGGAGAAGGAAGCCC
AGGTGGAGGCCACAAGCAGGAAAGAAAAGGCCAAGCAGAGGCCCCTGGCCCTGAACACT
GTGGAGATGCTGCGTGTGGCCAGCTCTTCTCTGGGCATGGGGCCGCAGCACGCCATGCA
GACGGCTGAGCGGCTCTACACGCAAGGCTACATCAGCTACCCAtggACAGAGACCACCCAC
TACCCTGAGAACTTTGACCTGAAGGGCTCTCTGCGGCAGCAGGCCAACCACCCCTACTGG
GCCGACACGGTGAAGCGGTTGTTAGCAGAAGGTATCAACCGCCCGCGGAAAGGCCATGAC
GCCGGCGACCATCCCCCCATCACCCCCATGAAGTCTGCCACAGAGGCCGAATTAGGGGGT
GACGCGTGGCGGCTCTATGAGTACATCACCAGACACTTCATCGCCACGGTCAGCCATGACT
GCAAGTACCTGCAGAGCACCATCTCCTTCAGAATTGGGCCCGAGCTCTTCACCTGCTCCGG
GAAGACCGTCCTCTCACCAGGCTTCACGGAGGTCATGCCCTGGCAGAGCGTGCCCCTGGA
GGAGAGCCTGCCCACTTGCCAGCGGGGTGATGCCTTCCCTGTGGGCGAGGTGAAGATGC
TGGAGAAGCAGACGAACCCACCCGACTACCTGACGGAGGCCGAGCTCATCACGCTCATGG
AGAAGCATGGCATCGGCACGGATGCCAGCATCCCTGTGCATATCAACAACATCTGCCAGCG
CAACTATGTCACGGTGGAGAGCGGGCGCCGGCTCAAGCCCACCAACCTCGGCATCGTCCT
GGTGCACGGCTACTATAAGATTGATGCAGAGCTGGTGCTCCCCACCATCCGCAGTGCAGTG
GAGAAGCAGCTGAACCTGATCGCCCAGGGCAAGGCCGACTACCGCCAGGTCCTGGGCCA
CACCCTGGACGTGTTCAAGAGGAAGTTCCACTACTTTGTCGACTCCATTGCTGGCATGGAT
GAGTTGATGGAGGTGTCTTTCTCGCCCCTGGCGGCCACAGGCAAGCCCCTCTCACGCTGT
GGGAAGTGCCACCGCTTCATGAAGTACATCCAGGCCAAGCCAAGCCGCCTGCACTGCTCC
CACTGCGATGAGACCTACACGCTCCCCCAGAACGGCACCATCAAGCTCTACAAGGAGCTC
CGCTGCCCTCTGGATGACTTCGAGCTGGTCCTGTGGTCATCAGGCTCTCGGGGCAAGAGC
TACCCGCTGTGCCCCTACTGCTACAACCACCCACCCTTCCGAGACATGAAGAAAGGCATGG
GCTGCAACGAGTGTACGCACCCCTCCTGCCAGCACTCGCTGAGCATGCTGGGCATCGGCC
AGTGCGTGGAATGTGAGAGCGGGGTGCTGGTGCTGGACCCCACCTCGGGCCCCAAGTGG

AAGGTGGCCTGCAACAAGTGCAACGTGGTAGCGCACTGCTTCGAGAACGCCCACCGCGTG
CGGGTGTCCGCCGACACCTGCAGTGTCTGTGAGGCCGCCTTGCTTGATGTGGACTTCAAC
AAGGCCAAGTCCCCACTCCCGGGCGATGAGACGCAGCACATGGGCTGCGTCTTTTGTGAC
CCCGTCTTCCAGGAGCTGGTGGAGCTGAAGCATGCGGCCTCCTGCCACCCCATGCACCGC
GGTGGACCAGGGAGAAGGCAGGGTCGAGGGCGGGGCCGGGCCAGGAGGCCCCCTGGG
AAGCCCAACCCCAGACGGCCCAAGGACAAGATGTCAGCCCTGGCCGCCTACTTTGTATAA

**3xFLAG-mEGFP-NES-TOP3B C666R (used in pGW1 and pcDNA5/FRT/TO)**

atgggaagcagcgactacaaagaccatgacggtgattataaagatcatgacatcgattacaaggatgacgatgacaagggttctgg
tgtgagcaagggcgaggagctgttcaccggggtggtgcccatcctggtcgagctggacggcgacgtaaacggccacaagttcagc
gtgtccggcgagggcgagggcgatgccacctacggcaagctgaccctgaagttcatctgcaccaccggcaagctgcccgtgccctg
gcccaccctcgtgaccaccctgacctacggcgtgcagtgcttcagccgctaccccgaccacatgaagcagcacgacttcttcaagtc
cgccatgcccgaaggctacgtccaggagcgcaccatcttcttcaaggacgacggcaactacaagacccgcgccgaggtgaagttc
gagggcgacaccctggtgaaccgcatcgagctgaagggcatcgacttcaaggaggacggcaacatcctggggcacaagctgga
gtacaactacaacagccacaacgtctatatcatggccgacaagcagaagaacggcatcaaggtgaacttcaagatccgccacaac
atcgaggacggcagcgtgcagctcgccgaccactaccagcagaacacccccatcggcgacggccccgtgctgctgcccgacaac
cactacctgagcacccagtccaagctgagcaaagaccccaacgagaagcgcgatcacatggtcctgctggagttcgtgaccgccg
ccgggatcactctcggcatggacgagctgtacaagggttctggtctgcagctgcctcccctggagcgcctgaccctggacggttcttct
AAGACTGTGCTCATGGTTGCTGAAAAGCCGTCCTTGGCACAGTCAATTGCCAAAATCCTCT
CTAGAGGGAGCCTGTCCTCACACAAAGGGCTGAACGGGGCCTGCTCAGTCCACGAGTACA
CTGGGACCTTTGCTGGCCAGCCAGTGCGCTTCAAGATGACGTCTGTCTGTGGTCACGTGA
TGACCCTGGATTTCCTGGGAAAATACAACAAATGGGACAAAGTGGACCCCGCAGAACTGTT
CAGCCAAGCTCCCACGGAGAAGAAAGAAGCTAACCCCAAGCTGAACATGGTGAAGTTCCT
GCAGGTGGAGGGCAGAGGCTGCGACTACATCGTGCTGTGGCTGGACTGCGACAAGGAGG
GGGAGAACATCTGCTTTGAGGTTCTTGATGCTGTTCTGCCCGTCATGAACAAGGCCCATGG
TGGCGAGAAGACCGTGTTCCGGGCCAGGTTTAGCTCCATCACGGACACAGACATCTGTAAT
GCCATGGCCTGCCTAGGCGAGCCTGACCACAACGAGGCGCTCTCAGTGGATGCTCGCCA
GGAGCTGGACCTGCGAATCGGCTGTGCATTCACCAGGTTTCAGACTAAATATTTCCAGGGG
AAATACGGTGATTTAGACAGCTCTCTCATCTCCTTTGGGCCGTGTCAGACTCCAACCCTGG
GATTCTGTGTGGAGAGACATGATAAAATCCAGTCCTTCAAACCAGAGACCTACTGGGTGCT
GCAGGCCAAGGTTAACACTGACAAAGACAGATCTCTCCTTTTGGACTGGGACCGAGTAAGA
GTGTTTGACCGGGAGATCGCACAGATGTTTTTAAACATGACAAAGCTGGAGAAGGAAGCCC
AGGTGGAGGCCACAAGCAGGAAAGAAAAGGCCAAGCAGAGGCCCCTGGCCCTGAACACT
GTGGAGATGCTGCGTGTGGCCAGCTCTTCTCTGGGCATGGGGCCGCAGCACGCCATGCA
GACGGCTGAGCGGCTCTACACGCAAGGCTACATCAGCTACCCACGGACAGAGACCACCCA
CTACCCTGAGAACTTTGACCTGAAGGGCTCTCTGCGGCAGCAGGCCAACCACCCCTACTG
GGCCGACACGGTGAAGCGGTTGTTAGCAGAAGGTATCAACCGCCCGCGGAAAGGCCATGA
CGCCGGCGACCATCCCCCCATCACCCCCATGAAGTCTGCCACAGAGGCCGAATTAGGGGG
TGACGCGTGGCGGCTCTATGAGTACATCACCAGACACTTCATCGCCACGGTCAGCCATGAC
TGCAAGTACCTGCAGAGCACCATCTCCTTCAGAATTGGGCCCGAGCTCTTCACCTGCTCCG
GGAAGACCGTCCTCTCACCAGGCTTCACGGAGGTCATGCCCTGGCAGAGCGTGCCCCTG
GAGGAGAGCCTGCCCACTTGCCAGCGGGGTGATGCCTTCCCTGTGGGCGAGGTGAAGAT
GCTGGAGAAGCAGACGAACCCACCCGACTACCTGACGGAGGCCGAGCTCATCACGCTCAT
GGAGAAGCATGGCATCGGCACGGATGCCAGCATCCCTGTGCATATCAACAACATCTGCCAG
CGCAACTATGTCACGGTGGAGAGCGGGCGCCGGCTCAAGCCCACCAACCTCGGCATCGT
CCTGGTGCACGGCTACTATAAGATTGATGCAGAGCTGGTGCTCCCCACCATCCGCAGTGCA
GTGGAGAAGCAGCTGAACCTGATCGCCCAGGGCAAGGCCGACTACCGCCAGGTCCTGGG
CCACACCCTGGACGTGTTCAAGAGGAAGTTCCACTACTTTGTCGACTCCATTGCTGGCATG
GATGAGTTGATGGAGGTGTCTTTCTCGCCCCTGGCGGCCACAGGCAAGCCCCTCTCACGC
TGTGGGAAGTGCCACCGCTTCATGAAGTACATCCAGGCCAAGCCAAGCCGCCTGCACTGC
TCCCACTGCGATGAGACCTACACGCTCCCCCAGAACGGCACCATCAAGCTCTACAAGGAG

CTCCGCcgcCCTCTGGATGACTTCGAGCTGGTCCTGTGGTCATCAGGCTCTCGGGGCAAGA
GCTACCCGCTGTGCCCCTACTGCTACAACCACCCACCCTTCCGAGACATGAAGAAAGGCAT
GGGCTGCAACGAGTGTACGCACCCCTCCTGCCAGCACTCGCTGAGCATGCTGGGCATCGG
CCAGTGCGTGGAATGTGAGAGCGGGGTGCTGGTGCTGGACCCCACCTCGGGCCCCAAGT
GGAAGGTGGCCTGCAACAAGTGCAACGTGGTAGCGCACTGCTTCGAGAACGCCCACCGC
GTGCGGGTGTCCGCCGACACCTGCAGTGTCTGTGAGGCCGCCTTGCTTGATGTGGACTTC
AACAAGGCCAAGTCCCCACTCCCGGGCGATGAGACGCAGCACATGGGCTGCGTCTTTTGT
GACCCCGTCTTCCAGGAGCTGGTGGAGCTGAAGCATGCGGCCTCCTGCCACCCCATGCAC
CGCGGTGGACCAGGGAGAAGGCAGGGTCGAGGGCGGGGCCGGGCCAGGAGGCCCCCT
GGGAAGCCCAACCCCAGACGGCCCAAGGACAAGATGTCAGCCCTGGCCGCCTACTTTGTA
TAA

**ORFs in plasmids for recombinant protein expression and purification**

**In the plasmids below, mEGFP is under regulation of a separate promoter in the opposite direction than the MBP fusion.*

**mEGFP___FLAG-MBP-twinSTII**
TTACTTGTACAGCTCGTCCATGCCGAGAGTGATCCCGGCGGCGGTCACGAACTCCAGCAG
GACCATGTGATCGCGCTTCTCGTTGGGGTCTTTGCTCAGCTTGGACTGGGTGCTCAGGTAG
TGGTTGTCGGGCAGCAGCACGGGGCCGTCGCCGATGGGGGTGTTCTGCTGGTAGTGGTC
GGCGAGCTGCACGCTGCCGTCCTCGATGTTGTGGCGGATCTTGAAGTTCACCTTGATGCC
GTTCTTCTGCTTGTCGGCCATGATATAGACGTTGTGGCTGTTGTAGTTGTACTCCAGCTTGT
GCCCCAGGATGTTGCCGTCCTCCTTGAAGTCGATGCCCTTCAGCTCGATGCGGTTCACCA
GGGTGTCGCCCTCGAACTTCACCTCGGCGCGGGTCTTGTAGTTGCCGTCGTCCTTGAAGA
AGATGGTGCGCTCCTGGACGTAGCCTTCGGGCATGGCGGACTTGAAGAAGTCGTGCTGCT
TCATGTGGTCGGGGTAGCGGCTGAAGCACTGCACGCCGTAGGTCAGGGTGGTCACGAGG
GTGGGCCAGGGCACGGGCAGCTTGCCGGTGGTGCAGATGAACTTCAGGGTCAGCTTGCC
GTAGGTGGCATCGCCCTCGCCCTCGCCGGACACGCTGAACTTGTGGCCGTTTACGTCGCC
GTCCAGCTCGACCAGGATGGGCACCACCCCGGTGAACAGCTCCTCGCCCTTGCTCACCAT
ggtgGGTACCGCATGCTATGCATCAGCTGCTAGCACCATGGCTCGAGATCCCGGGTGATCAA
GTCTTCGTCGAGTGATTGTAAATAAAATGTAATTTACAGTATAGTATTTTAATTAATATACAAATG
ATTTGATAATAATTCTTATTTAACTATAATATATTGTGTTGGGTTGAATTAAAGGTCCGTATACTC
CGGAATATTAATAGATCATGGAGATAATTAAAATGATAACCATCTCGCAAATAAATAAGTATTTT
ACTGTTTTCGTAACAGTTTTGTAATAAAAAAACCTATAAATATTCCGGATTATTCATACCGTCCC
ACCATCGGGCGCGGATCCCGGTCCGAAGCGCGCGGAATTCcaccatgggcagcggcGATTACAA
GGATGACGACGATAAGggttcttctatgaaaatcgaagaaggtaaactggtaatctggattaacggcgataaaggctataa
cggtctcgctgaagtcggtaagaaattcgagaaagataccggaattaaagtcaccgttgagcatccggataaactggaagagaaatt
cccacaggttgcggcaactggcgatggccctgacattatcttctgggcacacgaccgctttggtggctacgctcaatctggcctgttggc
tgaaatcaccccggacaaagcgttccaggacaagctgtatccgtttacctgggatgccgtacgttacaacggcaagctgattgcttacc
cgatcgctgttgaagcgttatcgctgatttataacaaagatctgctgccgaacccgccaaaaacctgggaagagatcccggcgctgg
ataaagaactgaaagcgaaaggtaagagcgcgctgatgttcaacctgcaagaaccgtacttcacctggccgctgattgctgctgacg
gggttatgcgttcaagtatgaaaacggcaagtacgacattaaagacgtgggcgtggataacgctggcgcgaaagcgggtctgacc
ttcctggttgacctgattaaaaacaaacacatgaatgcagacaccgattactccatcgcagaagctgcctttaataaaggcgaaacag
cgatgaccatcaacggcccgtgggcatggtccaacatcgacaccagcaaagtgaattatggtgtaacggtactgccgaccttcaag
ggtcaaccatccaaaccgttcgttggcgtgctgagcgcaggtattaacgccgccagtccgaacaaagagctggcaaaagagttcctc
gaaaactatctgctgactgatgaaggtctggaagcggttaataaagacaaaccgctgggtgccgtagcgctgaagtcttacgaggaa
gagttggcgaaagatccacgtattgccgccactatggaaaacgcccagaaaggtgaaatcatgccgaacatcccgcagatgtccg
ctttctggtatgccgtgcgtactgcggtgatcaacgccgccagcggtcgtcagactgtcgatgaagccctgaaagacgcgcagactaa

tgggatcgaggaaaacctgtacttccaatccggcagcggctggagccatccgcagtttgaaaaaggcagctggagccatccgcagt
ttgaaaaataa

**mEGFP___FLAG-MBP-ZnF (WT)-twinSTII**
TTACTTGTACAGCTCGTCCATGCCGAGAGTGATCCCGGCGGCGGTCACGAACTCCAGCAG
GACCATGTGATCGCGCTTCTCGTTGGGGTCTTTGCTCAGCTTGGACTGGGTGCTCAGGTAG
TGGTTGTCGGGCAGCAGCACGGGGCCGTCGCCGATGGGGGTGTTCTGCTGGTAGTGGTC
GGCGAGCTGCACGCTGCCGTCCTCGATGTTGTGGCGGATCTTGAAGTTCACCTTGATGCC
GTTCTTCTGCTTGTCGGCCATGATATAGACGTTGTGGCTGTTGTAGTTGTACTCCAGCTTGT
GCCCCAGGATGTTGCCGTCCTCCTTGAAGTCGATGCCCTTCAGCTCGATGCGGTTCACCA
GGGTGTCGCCCTCGAACTTCACCTCGGCGCGGGTCTTGTAGTTGCCGTCGTCCTTGAAGA
AGATGGTGCGCTCCTGGACGTAGCCTTCGGGCATGGCGGACTTGAAGAAGTCGTGCTGCT
TCATGTGGTCGGGGTAGCGGCTGAAGCACTGCACGCCGTAGGTCAGGGTGGTCACGAGG
GTGGGCCAGGGCACGGGCAGCTTGCCGGTGGTGCAGATGAACTTCAGGGTCAGCTTGCC
GTAGGTGGCATCGCCCTCGCCCTCGCCGGACACGCTGAACTTGTGGCCGTTTACGTCGCC
GTCCAGCTCGACCAGGATGGGCACCACCCCGGTGAACAGCTCCTCGCCCTTGCTCACCAT
ggtgGGTACCGCATGCTATGCATCAGCTGCTAGCACCATGGCTCGAGATCCCGGGTGATCAA
GTCTTCGTCGAGTGATTGTAAATAAAATGTAATTTACAGTATAGTATTTTAATTAATATACAAATG
ATTTGATAATAATTCTTATTTAACTATAATATATTGTGTTGGGTTGAATTAAAGGTCCGTATACTC
CGGAATATTAATAGATCATGGAGATAATTAAAATGATAACCATCTCGCAAATAAATAAGTATTTT
ACTGTTTTCGTAACAGTTTTGTAATAAAAAAACCTATAAATATTCCGGATTATTCATACCGTCCC
ACCATCGGGCGCGGATCCCGGTCCGAAGCGCGCGGAATTCcaccatgggcagcggcGATTACAA
GGATGACGACGATAAGggttcttctatgaaaatcgaagaaggtaaactggtaatctggattaacggcgataaaggctataa
cggtctcgctgaagtcggtaagaaattcgagaaagataccggaattaaagtcaccgttgagcatccggataaactggaagagaaatt
cccacaggttgcggcaactggcgatggccctgacattatcttctgggcacacgaccgctttggtggctacgctcaatctggcctgttggc
tgaaatcaccccggacaaagcgttccaggacaagctgtatccgtttacctgggatgccgtacgttacaacggcaagctgattgcttacc
cgatcgctgttgaagcgttatcgctgatttataacaaagatctgctgccgaacccgccaaaaacctgggaagagatcccggcgctgg
ataaagaactgaaagcgaaaggtaagagcgcgctgatgttcaacctgcaagaaccgtacttcacctggccgctgattgctgctgacg
gggggttatgcgttcaagtatgaaaacggcaagtacgacattaaagacgtgggcgtggataacgctggcgcgaaagcgggtctgacc
ttcctggttgacctgattaaaaacaaacacatgaatgcagacaccgattactccatcgcagaagctgcctttaataaaggcgaaacag
cgatgaccatcaacggcccgtgggcatggtccaacatcgacaccagcaaagtgaattatggtgtaacggtactgccgaccttcaag
ggtcaaccatccaaaccgttcgttggcgtgctgagcgcaggtattaacgccgccagtccgaacaaagagctggcaaaagagttcctc
gaaaactatctgctgactgatgaaggtctggaagcggttaataaagacaaaccgctgggtgccgtagcgctgaagtcttacgaggaa
gagttggcgaaagatccacgtattgccgccactatggaaaacgcccagaaaggtgaaatcatgccgaacatcccgcagatgtccg
ctttctggtatgccgtgcgtactgcggtgatcaacgccgccagcggtcgtcagactgtcgatgaagccctgaaagacgcgcagactaa
tgggatcgaggaaaacctgtacttccaatccaatattggaagtggaggcaagcccctctcacgctgtgggaagtgccaccgcttcatg
aagtacatccaggccaagccaagccgcctgcactgctcccactgcgatgagacctacacgctcccccagaacggcaccatcaagc
tctacaaggagctccgctgccctctggatgacttcgagctggtcctgtggtcatcaggctctcggggcaagagctacccgctgtgcccct
actgctacaaccacccacccttccgagacatgaagaaaggcatgggctgcaacgagtgtacgcacccctcctgccagcactcgctg
agcatgctgggcatcggccagtgcgtggaatgtgagagcggggtgctggtgctggaccccacctcgggccccaagtggaaggtgg
cctgcaacaagtgcaacgtggtagcgcactgcttcgagaacgcccaccgcgtgcgggtgtccgccgacacctgcagtgtctgtgag
gccgccttgcttgatgtggacttcaacaaggccaagtccccactcccgggcgatgagacgcagcacatggctgcgtcttttgtgacc
ccgtcttccaggagctggtggagctgaagcatgcggcctccggcagcggctggagccatccgcagtttgaaaaaggcagctggag
ccatccgcagtttgaaaaataa

**mEGFP___FLAG-MBP-ZnF (C666R)-twinSTII**
TTACTTGTACAGCTCGTCCATGCCGAGAGTGATCCCGGCGGCGGTCACGAACTCCAGCAG
GACCATGTGATCGCGCTTCTCGTTGGGGTCTTTGCTCAGCTTGGACTGGGTGCTCAGGTAG
TGGTTGTCGGGCAGCAGCACGGGGCCGTCGCCGATGGGGGTGTTCTGCTGGTAGTGGTC
GGCGAGCTGCACGCTGCCGTCCTCGATGTTGTGGCGGATCTTGAAGTTCACCTTGATGCC
GTTCTTCTGCTTGTCGGCCATGATATAGACGTTGTGGCTGTTGTAGTTGTACTCCAGCTTGT

GCCCCAGGATGTTGCCGTCCTCCTTGAAGTCGATGCCCTTCAGCTCGATGCGGTTCACCA
GGGTGTCGCCCTCGAACTTCACCTCGGCGCGGGTCTTGTAGTTGCCGTCGTCCTTGAAGA
AGATGGTGCGCTCCTGGACGTAGCCTTCGGGCATGGCGGACTTGAAGAAGTCGTGCTGCT
TCATGTGGTCGGGGTAGCGGCTGAAGCACTGCACGCCGTAGGTCAGGGTGGTCACGAGG
GTGGGCCAGGGCACGGGCAGCTTGCCGGTGGTGCAGATGAACTTCAGGGTCAGCTTGCC
GTAGGTGGCATCGCCCTCGCCCTCGCCGGACACGCTGAACTTGTGGCCGTTTACGTCGCC
GTCCAGCTCGACCAGGATGGGCACCACCCCGGTGAACAGCTCCTCGCCCTTGCTCACCAT
ggtgGGTACCGCATGCTATGCATCAGCTGCTAGCACCATGGCTCGAGATCCCGGGTGATCAA
GTCTTCGTCGAGTGATTGTAAATAAAATGTAATTTACAGTATAGTATTTTAATTAATATACAAATG
ATTTGATAATAATTCTTATTTAACTATAATATATTGTGTTGGGTTGAATTAAAGGTCCGTATACTC
CGGAATATTAATAGATCATGGAGATAATTAAAATGATAACCATCTCGCAAATAAATAAGTATTTT
ACTGTTTTCGTAACAGTTTTGTAATAAAAAAACCTATAAATATTCCGGATTATTCATACCGTCCC
ACCATCGGGCGCGGATCCCGGTCCGAAGCGCGCGGAATTCcaccatgggcagcggcGATTACAA
GGATGACGACGATAAGggttcttctatgaaaatcgaagaaggtaaactggtaatctggattaacggcgataaaggctataa
cggtctcgctgaagtcggtaagaaattcgagaaagataccggaattaaagtcaccgttgagcatccggataaactggaagagaaatt
cccacaggttgcggcaactggcgatggccctgacattatcttctgggcacacgaccgctttggtggctacgctcaatctggcctgttggc
tgaaatcaccccggacaaagcgttccaggacaagctgtatccgtttacctgggatgccgtacgttacaacggcaagctgattgcttacc
cgatcgctgttgaagcgttatcgctgatttataacaaagatctgctgccgaacccgccaaaaacctgggaagagatcccggcgctgg
ataaagaactgaaagcgaaaggtaagagcgcgctgatgttcaacctgcaagaaccgtacttcacctggccgctgattgctgctgacg
gggttatgcgttcaagtatgaaaacggcaagtacgacattaaagacgtgggcgtggataacgctggcgcgaaagcgggtctgacc
ttcctggttgacctgattaaaaacaaacacatgaatgcagacaccgattactccatcgcagaagctgcctttaataaaggcgaaacag
cgatgaccatcaacggcccgtgggcatggtccaacatcgacaccagcaaagtgaattatggtgtaacggtactgccgaccttcaag
ggtcaaccatccaaaccgttcgttggcgtgctgagcgcaggtattaacgccgccagtccgaacaaagagctggcaaaagagttcctc
gaaaactatctgctgactgatgaaggtctggaagcggttaataaagacaaaccgctgggtgccgtagcgctgaagtcttacgaggaa
gagttggcgaaagatccacgtattgccgccactatggaaaacgcccagaaaggtgaaatcatgccgaacatcccgcagatgtccg
ctttctggtatgccgtgcgtactgcggtgatcaacgccgccagcggtcgtcagactgtcgatgaagccctgaaagacgcgcagactaa
tgggatcgaggaaaacctgtacttccaatccaatattggaagtggaggcaagcccctctcacgctgtgggaagtgccaccgcttcatg
aagtacatccaggccaagccaagccgcctgcactgctcccactgcgatgagacctacacgctcccccagaacggcaccatcaagc
tctacaaggagctccgccgccctctggatgacttcgagctggtcctgtggtcatcaggctctcggggcaagagctacccgctgtgcccc
tactgctacaaccacccacccttccgagacatgaagaaaggcatgggctgcaacgagtgtacgcacccctcctgccagcactcgct
gagcatgctgggcatcggccagtgcgtggaatgtgagagcggggtgctggtgctggaccccacctcgggccccaagtggaaggtg
gcctgcaacaagtgcaacgtggtagcgcactgcttcgagaacgcccaccgcgtgcgggtgtccgccgacacctgcagtgtctgtga
ggccgccttgcttgatgtggacttcaacaaggccaagtccccactcccgggcgatgagacgcagcacatggctgcgtcttttgtgac
cccgtcttccaggagctggtggagctgaagcatgcggcctccggcagcggctggagccatccgcagtttgaaaaaggcagctgga
gccatccgcagtttgaaaaataa

**mEGFP___FLAG-MBP-ZnF (D669A)-twinSTII**

TTACTTGTACAGCTCGTCCATGCCGAGAGTGATCCCGGCGGCGGTCACGAACTCCAGCAG
GACCATGTGATCGCGCTTCTCGTTGGGGTCTTTGCTCAGCTTGGACTGGGTGCTCAGGTAG
TGGTTGTCGGGCAGCAGCACGGGGCCGTCGCCGATGGGGGTGTTCTGCTGGTAGTGGTC
GGCGAGCTGCACGCTGCCGTCCTCGATGTTGTGGCGGATCTTGAAGTTCACCTTGATGCC
GTTCTTCTGCTTGTCGGCCATGATATAGACGTTGTGGCTGTTGTAGTTGTACTCCAGCTTGT
GCCCCAGGATGTTGCCGTCCTCCTTGAAGTCGATGCCCTTCAGCTCGATGCGGTTCACCA
GGGTGTCGCCCTCGAACTTCACCTCGGCGCGGGTCTTGTAGTTGCCGTCGTCCTTGAAGA
AGATGGTGCGCTCCTGGACGTAGCCTTCGGGCATGGCGGACTTGAAGAAGTCGTGCTGCT
TCATGTGGTCGGGGTAGCGGCTGAAGCACTGCACGCCGTAGGTCAGGGTGGTCACGAGG
GTGGGCCAGGGCACGGGCAGCTTGCCGGTGGTGCAGATGAACTTCAGGGTCAGCTTGCC
GTAGGTGGCATCGCCCTCGCCCTCGCCGGACACGCTGAACTTGTGGCCGTTTACGTCGCC
GTCCAGCTCGACCAGGATGGGCACCACCCCGGTGAACAGCTCCTCGCCCTTGCTCACCAT
ggtgGGTACCGCATGCTATGCATCAGCTGCTAGCACCATGGCTCGAGATCCCGGGTGATCAA
GTCTTCGTCGAGTGATTGTAAATAAAATGTAATTTACAGTATAGTATTTTAATTAATATACAAATG

ATTTGATAATAATTCTTATTTAACTATAATATATTGTGTTGGGTTGAATTAAAGGTCCGTATACTC
CGGAATATTAATAGATCATGGAGATAATTAAAATGATAACCATCTCGCAAATAAATAAGTATTTT
ACTGTTTTCGTAACAGTTTTGTAATAAAAAAACCTATAAATATTCCGGATTATTCATACCGTCCC
ACCATCGGGCGCGGATCCCGGTCCGAAGCGCGCGGAATTCcaccatgggcagcggcGATTACAA
GGATGACGACGATAAGggttcttctatgaaaatcgaagaaggtaaactggtaatctggattaacggcgataaaggctataa
cggtctcgctgaagtcggtaagaaattcgagaaagataccggaattaaagtcaccgttgagcatccggataaactggaagagaaatt
cccacaggttgcggcaactggcgatggccctgacattatcttctgggcacacgaccgctttggtggctacgctcaatctggcctgttggc
tgaaatcaccccggacaaagcgttccaggacaagctgtatccgtttacctgggatgccgtacgttacaacggcaagctgattgcttacc
cgatcgctgttgaagcgttatcgctgatttataacaaagatctgctgccgaacccgccaaaaacctgggaagagatcccggcgctgg
ataaagaactgaaagcgaaaggtaagagcgcgctgatgttcaacctgcaagaaccgtacttcacctggccgctgattgctgctgacg
gggttatgcgttcaagtatgaaaacggcaagtacgacattaaagacgtgggcgtggataacgctggcgcgaaagcgggtctgacc
ttcctggttgacctgattaaaaacaaacacatgaatgcagacaccgattactccatcgcagaagctgcctttaataaaggcgaaacag
cgatgaccatcaacggcccgtgggcatggtccaacatcgacaccagcaaagtgaattatggtgtaacggtactgccgaccttcaag
ggtcaaccatccaaaccgttcgttggcgtgctgagcgcaggtattaacgccgccagtccgaacaaagagctggcaaaagagttcctc
gaaaactatctgctgactgatgaaggtctggaagcggttaataaagacaaaccgctgggtgccgtagcgctgaagtcttacgaggaa
gagttggcgaaagatccacgtattgccgccactatggaaaacgcccagaaaggtgaaatcatgccgaacatcccgcagatgtccg
ctttctggtatgccgtgcgtactgcggtgatcaacgccgccagcggtcgtcagactgtcgatgaagccctgaaagacgcgcagactaa
tgggatcgaggaaaacctgtacttccaatccaatattggaagtggaggcaagcccctctcacgctgtgggaagtgccaccgcttcatg
aagtacatccaggccaagccaagccgcctgcactgctcccactgcgatgagacctacacgctcccccagaacggcaccatcaagc
tctacaaggagctccgctgccctctggccgacttcgagctggtcctgtggtcatcaggctctcggggcaagagctacccgctgtgcccc
tactgctacaaccacccacccttccgagacatgaagaaaggcatgggctgcaacgagtgtacgcacccctcctgccagcactcgct
gagcatgctgggcatcggccagtgcgtggaatgtgagagcggggtgctggtgctggaccccacctcgggccccaagtggaaggtg
gcctgcaacaagtgcaacgtggtagcgcactgcttcgagaacgcccaccgcgtgcgggtgtccgccgacacctgcagtgtctgtga
ggccgccttgcttgatgtggacttcaacaaggccaagtccccactcccgggcgatgagacgcagcacatggctgcgtcttttgtgac
cccgtcttccaggagctggtggagctgaagcatgcggcctccggcagcggctggagccatccgcagtttgaaaaaggcagctgga
gccatccgcagtttgaaaaataa

**mEGFP___FLAG-MBP-ZnF (C688A/C691A)-twinSTII**

TTACTTGTACAGCTCGTCCATGCCGAGAGTGATCCCGGCGGCGGTCACGAACTCCAGCAG
GACCATGTGATCGCGCTTCTCGTTGGGGTCTTTGCTCAGCTTGGACTGGGTGCTCAGGTAG
TGGTTGTCGGGCAGCAGCACGGGGCCGTCGCCGATGGGGGTGTTCTGCTGGTAGTGGTC
GGCGAGCTGCACGCTGCCGTCCTCGATGTTGTGGCGGATCTTGAAGTTCACCTTGATGCC
GTTCTTCTGCTTGTCGGCCATGATATAGACGTTGTGGCTGTTGTAGTTGTACTCCAGCTTGT
GCCCCAGGATGTTGCCGTCCTCCTTGAAGTCGATGCCCTTCAGCTCGATGCGGTTCACCA
GGGTGTCGCCCTCGAACTTCACCTCGGCGCGGGTCTTGTAGTTGCCGTCGTCCTTGAAGA
AGATGGTGCGCTCCTGGACGTAGCCTTCGGGCATGGCGGACTTGAAGAAGTCGTGCTGCT
TCATGTGGTCGGGGTAGCGGCTGAAGCACTGCACGCCGTAGGTCAGGGTGGTCACGAGG
GTGGGCCAGGGCACGGGCAGCTTGCCGGTGGTGCAGATGAACTTCAGGGTCAGCTTGCC
GTAGGTGGCATCGCCCTCGCCCTCGCCGGACACGCTGAACTTGTGGCCGTTTACGTCGCC
GTCCAGCTCGACCAGGATGGGCACCACCCCGGTGAACAGCTCCTCGCCCTTGCTCACCAT
ggtgGGTACCGCATGCTATGCATCAGCTGCTAGCACCATGGCTCGAGATCCCGGGTGATCAA
GTCTTCGTCGAGTGATTGTAAATAAAATGTAATTTACAGTATAGTATTTTAATTAATATACAAATG
ATTTGATAATAATTCTTATTTAACTATAATATATTGTGTTGGGTTGAATTAAAGGTCCGTATACTC
CGGAATATTAATAGATCATGGAGATAATTAAAATGATAACCATCTCGCAAATAAATAAGTATTTT
ACTGTTTTCGTAACAGTTTTGTAATAAAAAAACCTATAAATATTCCGGATTATTCATACCGTCCC
ACCATCGGGCGCGGATCCCGGTCCGAAGCGCGCGGAATTCcaccatgggcagcggcGATTACAA
GGATGACGACGATAAGggttcttctatgaaaatcgaagaaggtaaactggtaatctggattaacggcgataaaggctataa
cggtctcgctgaagtcggtaagaaattcgagaaagataccggaattaaagtcaccgttgagcatccggataaactggaagagaaatt
cccacaggttgcggcaactggcgatggccctgacattatcttctgggcacacgaccgctttggtggctacgctcaatctggcctgttggc
tgaaatcaccccggacaaagcgttccaggacaagctgtatccgtttacctgggatgccgtacgttacaacggcaagctgattgcttacc
cgatcgctgttgaagcgttatcgctgatttataacaaagatctgctgccgaacccgccaaaaacctgggaagagatcccggcgctgg

ataaagaactgaaagcgaaaggtaagagcgcgctgatgttcaacctgcaagaaccgtacttcacctggccgctgattgctgctgacg
ggggttatgcgttcaagtatgaaaacggcaagtacgacattaaagacgtgggcgtggataacgctggcgcgaaagcgggtctgacc
ttcctggttgacctgattaaaaacaaacacatgaatgcagacaccgattactccatcgcagaagctgcctttaataaaggcgaaacag
cgatgaccatcaacggcccgtgggcatggtccaacatcgacaccagcaaagtgaattatggtgtaacggtactgccgaccttcaag
ggtcaaccatccaaaccgttcgttggcgtgctgagcgcaggtattaacgccgccagtccgaacaaagagctggcaaaagagttcctc
gaaaactatctgctgactgatgaaggtctggaagcggttaataaagacaaaccgctgggtgccgtagcgctgaagtcttacgaggaa
gagttggcgaaagatccacgtattgccgccactatggaaaacgcccagaaaggtgaaatcatgccgaacatcccgcagatgtccg
ctttctggtatgccgtgcgtactgcggtgatcaacgccgccagcggtcgtcagactgtcgatgaagccctgaaagacgcgcagactaa
tgggatcgaggaaaacctgtacttccaatccaatattggaagtggaggcaagcccctctcacgctgtgggaagtgccaccgcttcatg
aagtacatccaggccaagccaagccgcctgcactgctcccactgcgatgagacctacacgctcccccagaacggcaccatcaagc
tctacaaggagctccgctgccctctggatgacttcgagctggtcctgtggtcatcaggctctcggggcaagagctacccgctggccccc
tacgcctacaaccacccacccttccgagacatgaagaaaggcatgggctgcaacgagtgtacgcacccctcctgccagcactcgct
gagcatgctgggcatcggccagtgcgtggaatgtgagagcggggtgctggtgctggaccccacctcgggccccaagtggaaggtg
gcctgcaacaagtgcaacgtggtagcgcactgcttcgagaacgcccaccgcgtgcgggtgtccgccgacacctgcagtgtctgtga
ggccgccttgcttgatgtggacttcaacaaggccaagtccccactcccgggcgatgagacgcagcacatgggctgcgtcttttgtgac
cccgtcttccaggagctggtggagctgaagcatgcggcctccggcagcggctggagccatccgcagtttgaaaaaggcagctggtgga
gccatccgcagtttgaaaaataa

**His6-TOP3B(WT)-FLAG**

ATGggttcttctcaccatcaccatcaccatggttcttctAAGACTGTGCTCATGGTTGCTGAAAAGCCGTCCTT
GGCACAGTCAATTGCCAAAATCCTCTCTAGAGGGAGCCTGTCCTCACACAAAGGGCTGAAC
GGGGCCTGCTCAGTCCACGAGTACACTGGGACCTTTGCTGGCCAGCCAGTGCGCTTCAAG
ATGACGTCTGTCTGTGGTCACGTGATGACCCTGGATTTCCTGGGAAAATACAACAAATGGG
ACAAAGTGGACCCCGCAGAACTGTTCAGCCAAGCTCCCACGGAGAAGAAAGAAGCTAACC
CCAAGCTGAACATGGTGAAGTTCCTGCAGGTGGAGGGCAGAGGCTGCGACTACATCGTGC
TGTGGCTGGACTGCGACAAGGAGGGGGAGAACATCTGCTTTGAGGTTCTTGATGCTGTTCT
GCCCGTCATGAACAAGGCCCATGGTGGCGAGAAGACCGTGTTCCGGGCCAGGTTTAGCTC
CATCACGGACACAGACATCTGTAATGCCATGGCCTGCCTAGGCGAGCCTGACCACAACGAG
GCGCTCTCAGTGGATGCTCGCCAGGAGCTGGACCTGCGAATCGGCTGTGCATTCACCAGG
TTTCAGACTAAATATTTCCAGGGGAAATACGGTGATTTAGACAGCTCTCTCATCTCCTTTGGG
CCGTGTCAGACTCCAACCCTGGGATTCTGTGTGGAGAGACATGATAAAATCCAGTCCTTCA
AACCAGAGACCTACTGGGTGCTGCAGGCCAAGGTTAACACTGACAAAGACAGATCTCTCCT
TTTGGACTGGGACCGAGTAAGAGTGTTTGACCGGGAGATCGCACAGATGTTTTTAAACATG
ACAAAGCTGGAGAAGGAAGCCCAGGTGGAGGCCACAAGCAGGAAAGAAAAGGCCAAGCA
GAGGCCCCTGGCCCTGAACACTGTGGAGATGCTGCGTGTGGCCAGCTCTTCTCTGGGCAT
GGGGCCGCAGCACGCCATGCAGACGGCTGAGCGGCTCTACACGCAAGGCTACATCAGCTA
CCCACGGACAGAGACCACCCACTACCCTGAGAACTTTGACCTGAAGGGCTCTCTGCGGCA
GCAGGCCAACCACCCCTACTGGGCCGACACGGTGAAGCGGTTGTTAGCAGAAGGTATCAA
CCGCCCGCGGAAAGGCCATGACGCCGGCGACCATCCCCCCATCACCCCCATGAAGTCTGC
CACAGAGGCCGAATTAGGGGGTGACGCGTGGCGGCTCTATGAGTACATCACCAGACACTT
CATCGCCACGGTCAGCCATGACTGCAAGTACCTGCAGAGCACCATCTCCTTCAGAATTGGG
CCCGAGCTCTTCACCTGCTCCGGGAAGACCGTCCTCTCACCAGGCTTCACGGAGGTCATG
CCCTGGCAGAGCGTGCCCCTGGAGGAGAGCCTGCCCACTTGCCAGCGGGGTGATGCCTT
CCCTGTGGGCGAGGTGAAGATGCTGGAGAAGCAGACGAACCCACCCGACTACCTGACGG
AGGCCGAGCTCATCACGCTCATGGAGAAGCATGGCATCGGCACGGATGCCAGCATCCCTG
TGCATATCAACAACATCTGCCAGCGCAACTATGTCACGGTGGAGAGCGGGCGCCGGCTCA
AGCCCACCAACCTCGGCATCGTCCTGGTGCACGGCTACTATAAGATTGATGCAGAGCTGGT
GCTCCCCACCATCCGCAGTGCAGTGGAGAAGCAGCTGAACCTGATCGCCCAGGGCAAGG
CCGACTACCGCCAGGTCCTGGGCCACACCCTGGACGTGTTCAAGAGGAAGTTCCACTACT
TTGTCGACTCCATTGCTGGCATGGATGAGTTGATGGAGGTGTCTTTCTCGCCCCTGGCGGC
CACAGGCAAGCCCCTCTCACGCTGTGGGAAGTGCCACCGCTTCATGAAGTACATCCAGGC

CAAGCCAAGCCGCCTGCACTGCTCCCACTGCGATGAGACCTACACGCTCCCCCAGAACGG
CACCATCAAGCTCTACAAGGAGCTCCGCTGCCCTCTGGATGACTTCGAGCTGGTCCTGTG
GTCATCAGGCTCTCGGGGCAAGAGCTACCCGCTGTGCCCCTACTGCTACAACCACCCACC
CTTCCGAGACATGAAGAAAGGCATGGGCTGCAACGAGTGTACGCACCCCTCCTGCCAGCA
CTCGCTGAGCATGCTGGGCATCGGCCAGTGCGTGGAATGTGAGAGCGGGGTGCTGGTGC
TGGACCCCACCTCGGGCCCCAAGTGGAAGGTGGCCTGCAACAAGTGCAACGTGGTAGCG
CACTGCTTCGAGAACGCCCACCGCGTGCGGGTGTCCGCCGACACCTGCAGTGTCTGTGA
GGCCGCCTTGCTTGATGTGGACTTCAACAAGGCCAAGTCCCCACTCCCGGGCGATGAGAC
GCAGCACATGGGCTGCGTCTTTTGTGACCCCGTCTTCCAGGAGCTGGTGGAGCTGAAGCA
TGCGGCCTCCTGCCACCCCATGCACCGCGGTGGACCAGGGAGAAGGCAGGGTCGAGGG
CGGGGCCGGGCCAGGAGGCCCCCTGGGAAGCCCAACCCCAGACGGCCCAAGGACAAGA
TGTCAGCCCTGGCCGCCTACTTTGTAAGCGGACCGACGCGTACGCGGCCGCTCGAGCAGA
AACTCATCTCAGAAGAGGATCTGGCAGCAAATGATATCCTGGATTACAAGGATGACGACGAT
AAGGTTTAA

**His6-TOP3B(Y336F)-FLAG**
ATGggttcttctcaccatcaccatcaccatggttcttctAAGACTGTGCTCATGGTTGCTGAAAAGCCGTCCTT
GGCACAGTCAATTGCCAAAATCCTCTCTAGAGGGAGCCTGTCCTCACACAAAGGGCTGAAC
GGGGCCTGCTCAGTCCACGAGTACACTGGGACCTTTGCTGGCCAGCCAGTGCGCTTCAAG
ATGACGTCTGTCTGTGGTCACGTGATGACCCTGGATTTCCTGGGAAAATACAACAAATGGG
ACAAAGTGGACCCCGCAGAACTGTTCAGCCAAGCTCCCACGGAGAAGAAAGAAGCTAACC
CCAAGCTGAACATGGTGAAGTTCCTGCAGGTGGAGGGCAGAGGCTGCGACTACATCGTGC
TGTGGCTGGACTGCGACAAGGAGGGGGAGAACATCTGCTTTGAGGTTCTTGATGCTGTTCT
GCCCGTCATGAACAAGGCCCATGGTGGCGAGAAGACCGTGTTCCGGGCCAGGTTTAGCTC
CATCACGGACACAGACATCTGTAATGCCATGGCCTGCCTAGGCGAGCCTGACCACAACGAG
GCGCTCTCAGTGGATGCTCGCCAGGAGCTGGACCTGCGAATCGGCTGTGCATTCACCAGG
TTTCAGACTAAATATTTCCAGGGGAAATACGGTGATTTAGACAGCTCTCTCATCTCCTTTGGG
CCGTGTCAGACTCCAACCCTGGGATTCTGTGTGGAGAGACATGATAAAATCCAGTCCTTCA
AACCAGAGACCTACTGGGTGCTGCAGGCCAAGGTTAACACTGACAAAGACAGATCTCTCCT
TTTGGACTGGGACCGAGTAAGAGTGTTTGACCGGGAGATCGCACAGATGTTTTTAAACATG
ACAAAGCTGGAGAAGGAAGCCCAGGTGGAGGCCACAAGCAGGAAAGAAAAGGCCAAGCA
GAGGCCCCTGGCCCTGAACACTGTGGAGATGCTGCGTGTGGCCAGCTCTTCTCTGGGCAT
GGGGCCGCAGCACGCCATGCAGACGGCTGAGCGGCTCTACACGCAAGGCTACATCAGCttt
CCACGGACAGAGACCACCCACTACCCTGAGAACTTTGACCTGAAGGGCTCTCTGCGGCAG
CAGGCCAACCACCCCTACTGGGCCGACACGGTGAAGCGGTTGTTAGCAGAAGGTATCAAC
CGCCCGCGGAAAGGCCATGACGCCGGCGACCATCCCCCCATCACCCCCATGAAGTCTGCC
ACAGAGGCCGAATTAGGGGGTGACGCGTGGCGGCTCTATGAGTACATCACCAGACACTTC
ATCGCCACGGTCAGCCATGACTGCAAGTACCTGCAGAGCACCATCTCCTTCAGAATTGGGC
CCGAGCTCTTCACCTGCTCCGGGAAGACCGTCCTCTCACCAGGCTTCACGGAGGTCATGC
CCTGGCAGAGCGTGCCCCTGGAGGAGAGCCTGCCCACTTGCCAGCGGGGTGATGCCTTC
CCTGTGGGCGAGGTGAAGATGCTGGAGAAGCAGACGAACCCACCCGACTACCTGACGGA
GGCCGAGCTCATCACGCTCATGGAGAAGCATGGCATCGGCACGGATGCCAGCATCCCTGT
GCATATCAACAACATCTGCCAGCGCAACTATGTCACGGTGGAGAGCGGGCGCCGGCTCAA
GCCCACCAACCTCGGCATCGTCCTGGTGCACGGCTACTATAAGATTGATGCAGAGCTGGTG
CTCCCCACCATCCGCAGTGCAGTGGAGAAGCAGCTGAACCTGATCGCCCAGGGCAAGGC
CGACTACCGCCAGGTCCTGGGCCACACCCTGGACGTGTTCAAGAGGAAGTTCCACTACTT
TGTCGACTCCATTGCTGGCATGGATGAGTTGATGGAGGTGTCTTTCTCGCCCCTGGCGGCC
ACAGGCAAGCCCCTCTCACGCTGTGGGAAGTGCCACCGCTTCATGAAGTACATCCAGGCC
AAGCCAAGCCGCCTGCACTGCTCCCACTGCGATGAGACCTACACGCTCCCCCAGAACGGC
ACCATCAAGCTCTACAAGGAGCTCCGCTGCCCTCTGGATGACTTCGAGCTGGTCCTGTGGT
CATCAGGCTCTCGGGGCAAGAGCTACCCGCTGTGCCCCTACTGCTACAACCACCCACCCTT

CCGAGACATGAAGAAAGGCATGGGCTGCAACGAGTGTACGCACCCCTCCTGCCAGCACTC
GCTGAGCATGCTGGGCATCGGCCAGTGCGTGGAATGTGAGAGCGGGGTGCTGGTGCTGG
ACCCCACCTCGGGCCCCAAGTGGAAGGTGGCCTGCAACAAGTGCAACGTGGTAGCGCAC
TGCTTCGAGAACGCCCACCGCGTGCGGGTGTCCGCCGACACCTGCAGTGTCTGTGAGGC
CGCCTTGCTTGATGTGGACTTCAACAAGGCCAAGTCCCCACTCCCGGGCGATGAGACGCA
GCACATGGGCTGCGTCTTTTGTGACCCCGTCTTCCAGGAGCTGGTGGAGCTGAAGCATGC
GGCCTCCTGCCACCCCATGCACCGCGGTGGACCAGGGAGAAGGCAGGGTCGAGGGCGG
GGCCGGGCCAGGAGGCCCCCTGGGAAGCCCAACCCCAGACGGCCCAAGGACAAGATGTC
AGCCCTGGCCGCCTACTTTGTAAGCGGACCGACGCGTACGCGGCCGCTCGAGCAGAAACT
CATCTCAGAAGAGGATCTGGCAGCAAATGATATCCTGGATTACAAGGATGACGACGATAAGG
TTTAA
